# Codon-usage optimization in the prokaryotic tree of life: How synonymous codons are differentially selected in sequence domains with different expression levels and degrees of conservation

**DOI:** 10.1101/2020.02.12.942524

**Authors:** José Luis López, Mauricio Javier Lozano, María Laura Fabre, Antonio Lagares

**Affiliations:** IBBM - Instituto de Biotecnología y Biología Molecular, CONICET, CCT-La Plata, Departamento de Ciencias Biológicas, Facultad de Ciencias Exactas, Universidad Nacional de La Plata, calles 47 y 115, 1900-La Plata, Argentina.

**Keywords:** codon usage selection, mutational bias, genome evolution, core genes, singletons, translation efficiency, translation accuracy

## Abstract

Prokaryote genomes exhibit a wide range of GC contents and codon usages, both resulting from an interaction between mutational bias and natural selection. In order to investigate the basis underlying specific codon changes, we performed a comprehensive analysis of 29-different prokaryote families. The analysis of core-gene sets with increasing ancestries in each family lineage revealed that the codon usages became progressively more adapted to the tRNA pools. While, as previously reported, highly-expressed genes presented the more optimized codon usage, the singletons contained the less selectively-favored codons. Results showed that usually codons with the highest translational adaptation were preferentially enriched. In agreement with previous reports, a C-bias in 2- to 3-fold codons, and a U-bias in 4-fold codons occurred in all families, irrespective of the global genomic-GC content. Furthermore, the U-biases suggested that U_3_-mRNA–U_34_-tRNA interactions were responsible for a prominent codon optimization in both the more ancestral core and the highly expressed genes. A comparative analysis of sequences that encode conserved-(*cr*) or variable-(v*r*) translated products, with each one being under high- (HEP) and low- (LEP) expression levels, demonstrated that the efficiency was more relevant (by a factor of 2) than accuracy to modelling codon usage. Finally, analysis of the third position of codons (GC3) revealed that, in genomes of global-GC contents higher than 35-40%, selection favored a GC3 increase; whereas in genomes with very low-GC contents, a decrease in GC3 occurred. A comprehensive final model is presented where all patterns of codon usage variations are condensed in five-distinct behavioral groups.

**IMPORTANCE:** The prokaryotic genomes—the current heritage of the more ancient life forms on earth— are comprised of diverse gene sets; all characterized by varied origins, ancestries, and spatial-temporal–expression patterns. Such genetic diversity has for a long time raised the question of how cells shape their coding strategies to optimize protein demands (*i.e.*, product abundance) and accuracy (*i.e.*, translation fidelity) through the use of the same genetic code in genomes with GC-contents that range from less than 20 to over 80%. In this work, we present evidence on how codon usage is adjusted in the prokaryote tree of life, and on how specific biases have operated to improve translation. Through the use of proteome data, we characterized conserved and variable sequence domains in genes of either high- or low-expression level, and quantitated the relative weight of efficiency and accuracy—as well as their interaction—in shaping codon usage in prokaryotes.

## INTRODUCTION

The wide range of GC contents exhibited by prokaryote genomes—*i. e.*, from less than 20 to 80%—are believed to be primarily caused by interspecies differences in mutational processes that operate on both the coding and the noncoding regions (1–6). Since prokaryote genomes consist mainly of coding regions that tightly reflect the genomic GC content, mutational bias is a main force that shapes the codon usage of the majority of the genes (7, 8). Thus, understanding how selection is coupled to mutational processes to model codon usage under such diverse GC contents is an essential issue (9–11). Recent evidence suggests that prokaryotic genomes with intermediate-to-high GC contents are affected by mutations that are universally biased in favor of AT replacements (12, 13). That process is counterbalanced by selection-based constraints that, in turn, increase the GC content and fine-tune codon usage—*i. e.*, the so-called mutation-selection-drift model (14–16). Intragenomic codon-usage heterogeneities, however, are always present among different gene sets—*i. e.*, between core genes that are shared throughout a given lineage, and singletons (unique accessory genes) that are taxa- and/or strain-specific (17, 18). Furthermore, in a multipartite genome, the linkage between the physical patterns of heterogeneity in codon usage and the replicon location of the different core genes has also been recently demonstrated (19). The analysis of intragenomic codon usage heterogeneities by different authors (20, 21) has served to identify at least the following three distinctive gene groups: The first comprises the majority of the coding sequences that are associated with the so-called typical codon usage, while the second consists of the putative highly expressed (PHE) genes involving codon usages that are the best adapted to the translational machinery (20, 22–26). The third contains genes that encode the accessory information including the singletons (unique genes) that are present in mobile genetic elements as well as in the more stable replicons (27–31). The intracellular variations in codon usage can be explained on the basis of selective pressures that operate with different strengths depending on gene function and the resulting impact on cellular fitness (32). A search for the biochemical basis associated with the heterogeneity in codon usage among different gene sets has been the focus of numerous studies. Several lines of evidence have indicated that the biased codon usage in PHE genes correlates with the copy number of the specific tRNA species that decode the preferred codons (23, 33, 34) and with an optimal codon-anticodon interaction (35). The latter includes both the classical Watson-and-Crick interactions (WCIs) and a wobble-base pairing with the corresponding cognate tRNAs. All these interactions have been taken into consideration in order to define different numerical indices (36, 37) as estimators of the codon adaptation to the existing tRNA pool. Though not considered in currently used translation-adaptation indices, evidence has also been found for other nonstandard codon-anticodon interactions which, by improving the decoding capacity, are also relevant to codon-usage evolution (38–41).

The analysis of an extensive number of genes with different functions, ubiquitousness, and degrees of phylogenetic conservation has demonstrated that codon usage is related to gene-expression level (33, 42, 43), the degree of conservation (18, 32, 44, 45), the genomic location—*i. e.*, chromosome, chromid, plasmidome (19, 46, 47)—and different features such as codon ramps, and mRNA secondary structure, among others (48–50). Current evidence indicates that accessory genes involve atypical codon usages (21, 47, 51, 52) compared to the more conserved (ancestral) core genes in a given lineage. These latter genes, for their part, exhibit adaptational variations in codon usages ranging from typical to more biased as the one observed in genes that correspond to highly abundant proteins which are coded by PHE (53). Moreover, that core genes may also exhibit remarkable codon-usage heterogeneities has been recently demonstrated (19).

In the work reported here after examining 29 different prokaryote families, we performed a consolidated analysis aimed at characterizing the specific intragenomic codon variations that lead to differences in codon usage between gene sets with diverse expression levels and degrees of conservation in a given lineage. The evaluation of intragenic regions with different coding characteristics—compared to strategies based on the global analyses of complete genes—enabled the recognition of different patterns of codon usages within a message to be translated. Thus the questions emerged of (i) whether the codon-usage patterns associated with highly expressed amino-acid sequences (*i. e.*, affecting efficiency) were the same as those associated with genes encoding highly conserved sequences (*i. e.*, affecting accuracy), and (ii) whether the requirements for translation efficiency and accuracy were fully independent or whether those two types of demands interacted. The results have indicated how, even in organisms with quite different GC contents, alterations in specific codons are associated with a selective adaptation of the more ancestral genes compared to the adaptation of those genes that are newer in the phylogeny. Through an independent analysis of sequences associated with variable or conserved regions having different expression levels (*i. e.*, low *versus* high), we were able to identify the specific codon usages associated with translation efficiency and accuracy as well as quantitatively estimate their relative relevance to codon usage.

## MATERIALS AND METHODS

### Prokaryote families selected for analysis in this work and identification of core genes and singletons

We screened the EDGAR public-projects database (54, 55) available at https://edgar.computational.bio.uni-giessen.de/cgi-bin/edgar_login.cgi, chose several prokaryote families that included at least 20 complete genomes each, and finally selected 27 bacterial and 2 archaeal families (Table S1_a, tab 1). A specific core-gene set was defined as a group of genes whose orthologs are present in a given set of species under investigation. For each of the families selected sequential core-gene sets with increasing ancestry (C1 through Cn) were calculated. To that end, first the phylogenetic tree for each family was extracted from EDGAR and one species per family chosen as a reference. Next, the different core-gene sets were obtained by incorporating into the analysis new species having sequentially increasing phylogenetic distances from the reference strain (accordingly, by following the tree from the branches to the root). Table S1_a-c, lists the phylogenetic trees used for these calculations as well as the particular species that were included in each core-gene set (C1 to Cn) for the different prokaryote families. The phylogenetic trees were edited with the Figtree (56) and Inkskape programs (TEAM-Inkscape). At least six core-gene sets differing in size from *ca*. 50 to 100 genes each were calculated per family. In each prokaryote family, the most ancestral core-gene set (Cn) consisted of 100 to 500 orthologs. Table S2, tabs 2 to 30 lists the singletons— those corresponding to genes that were specific to the reference strains with no orthologs within the family—as calculated with EDGAR.

### PHE genes

For each of the selected reference genomes, we obtained a set of genes encoding ribosomal proteins and tRNA synthetases (24, 57). Table S2, tab 1 itemizes the PHE genes whose orthologs were obtained and analyzed in each reference genome.

### Highly (HEP) and lowly (LEP) expressed proteins within the same core-gene set

Integrated expression data from the Protein Abundance Database (PaxDB; (58)) were retrieved for the bacterial strains *Yersinia pestis* CO92, *Streptococcus pyogenes* M1 GAS, *Campylobacter jejuni* subsp. jejuni NCTC 11168, *Bacillus subtilis* subsp. subtilis str. 168, *Bacteroides thetaiotaomicron* VPI-5482, and *Mycobacterium tuberculosis* H37Rv. Assuming that orthologs have comparable expression levels within the same—or closely related—species and using the PaxDB data from the above indicated 6 strains, we inferred putative expression data for the proteomes of the microorganisms presented in Figs. 4 to 6, and listed in Table S3. Then, for selected core fractions, we obtained one subset of genes encoding HEP plus another subset codifying LEP. For 23 out of the 29 prokaryotic genomes that we analyzed here, no proteome data were available, nor were any in phylogenetically related microorganisms.

### Analysis of codon usage in gene-sequence regions that encode either conserved (*cr*) or variable (*vr*) amino-acid positions in the HEP and LEP subsets

Individual genes that belonged to the HEP and LEP groups were aligned with the corresponding orthologs. Then codons corresponding to conserved and variable amino-acid positions in the HEP genes were separated and each concatenated to generate the HEP_*cr*, HEP_*vr* sequence groups. Through the use of a similar procedure with the LEP genes, the LEP_*cr*, and LEP_*vr* sequences were also generated. Codons categorized as belonging to the *cr* group were those associated with positions with fully conserved amino acids throughout the alignment. Codons categorized as belonging to the *vr* group were those associated with positions where none of the amino acids aligned (at that specific point) reached a proportion higher than 0.5. The modal codon usage (47) of each collection of *cr* and *vr* sequences were calculated and used for further analysis.

### Raw codon counts (RCC)-based Correspondence analyses (CAs)

The RCC-based CAs were performed using bash and R-software homemade scripts which can be found at CUBES software page (this work, available at https://github.com/maurijlozano/CUBES). Briefly, G. Olsen codon usage software was used to count codons on coding sequences (available at http://www.life.illinois.edu/gary/programs.html), data were loaded on R, and the correspondence analyses were run using FactoMiner (59) and Factoextra (https://CRAN.R-project.org/package=factoextra) packages. Plots were made using ggplot2, ggrepel, ggthemes and gtools R packages. For each core-gene set the CA coordinates were calculated as the arithmetic mean of the first and second dimensions of all the genes present in that set (centroids). Then, a plot was generated containing all the coding sequences, together with the projections of the core-gene sets (C1 to Cn), the singletons and PHE genes.

### Relative synonymous-codon usages (RSCUs)-based CAs, and calculation of modal codon usages

The RSCUs (60) of all individual genes from a given genome were calculated by CodonW with DNA sequences as input data (61) and then used to perform the 59-variable–based correspondence analysis (CA)—*i. e.*, with all the codons except those for Met (AUG), Trp (UGG), and the three stop codons (UAA, UAG, and UGA). The modal codon usages (47) were calculated for the core genes, singletons, and PHE genes. Artificial sequences representing modal codon usages (*i. e.*, modal sequences) and the amino-acid composition present in each core fraction (Cn) were generated through the use of a homemade Perl script (calculate_modals2.pl) from the CUBES package. In order to accurately represent the modal codon usage, particularly for synonymous codons from low-abundance amino acids, modal sequences were designed with a length of at least ten thousand codons. These modal sequences were used as an additional input in their respective CAs. CA plots were generated through the use of Ggplot2 program (62) and edited with Inkskape (TEAM-Inkscape).

### tRNA-gene–copy number and modal species-specific tRNA-adaptation index (m-tAI)

The gene-copy number of each tRNA in the different prokaryote species studied here were downloaded from the GtRNAdb server (http://gtrnadb.ucsc.edu), which website uses predictions made by the program tRNAscan-SE (63). For each reference genome, the copy number for the tRNAs and the sequences of all the open reading frames were used to train the *S_ij_* weights as previously reported, with that parameter estimating the efficiency of the interaction between the *i*th codon and the *j*th anticodon in a given organism (36, 37). The procedure stated in brief: With a given *n*, and randomly generated *S_ij_* starting points—*i. e.*, having values that range between 0 and 1—the tAI was calculated for each coding sequence by means of the tAI package ((36), https://github.com/mariodosreis/tai). Next, the directional codon-bias score (DCBS; (37)) was calculated through the use of the script seq2DCBS.pl (CUBES package). Finally, the Nelder-Mead optimization algorithm from R project was used (instead of the hill-climbing algorithm) to search for the *S_ij_* value that maximized the Spearman rank correlation between the DCBS and the tAI. A script for bulk *S_ij_* estimation is available in the CUBES package (https://github.com/maurijlozano/CUBES, calculate_sopt_DCBS_GNM_f.sh). The estimated sets of *S_ij_* weights were used to calculate the modal tRNA-adaptation index (m-tAI) for different species and gene sets (*i. e.*, core and PHE genes plus singletons) as a measure of their efficiency in being recognized by the intracellular tRNA pool. The m-tAIs were calculated from the previously generated modal sequences by means of the tAI_Modal_g.sh script from the CUBES package.

## RESULTS

### Ancestry-dependent codon-usage bias as revealed by the analysis of core genes from diverse prokaryotic families

López et al. (19) have recently demonstrated that, in a model proteobacterium, the more ancestral the core genes were the better adapted their codon usages were to the translational machinery. In order to investigate if such correlation was associated with a general phenomenon in different prokaryote taxa, we assembled different core-gene sets that progressed deeper into the phylogenies of 27 Gram-negative and -positive eubacterial families spanning the phyla Proteobacteria, Actinobacteria, Firmicutes, and Bacteroidetes along with 2 archaeal families from the phylum Euryarchaeota. Table S1_a (tab 1) itemizes for each taxon the number of genes in each gene set from the most recent core 1 (C1), to the most ancestral core n (Cn). The codon-usage variation with gene ancestry within a given prokaryote family was evaluated through a correspondence analysis (CA) that used as variables the raw codon counts (RCC) of the individual genes in each genome analyzed (see Materials and Methods). The average values of the first two components for the core-gene sets C1 to Cn were projected on the CA plots. Fig. 1 (left panels) depicts the CAs for four different genomes specifically selected to represent groups of organisms with different types of CA plots and GC contents, namely Groups A to D. CAs were also calculated using RSCUs as input variables instead of RCC as presented in Fig S1A. In agreement with a recent study in *Sinorhizobium meliloti* (19), in all instances a directional shift in the codon usage positions was evident from the most recent (C1) towards the most ancestral (Cn) core-gene set. That this ancestry-dependent pattern of codon-usage variation had been observed in even quite distant prokaryote families among those analyzed here was remarkable (*cf*. the CA plots for all other species in Fig. S1B, left panels). In the evolution of core codon usages, however, the extent of the observed shifts and the type of synonymous codons enriched in each taxon (*i. e.*, the direction of change) varied markedly among different families (Fig. 1, Fig S1A and Fig S1B, right panels).

**Fig. 1.**
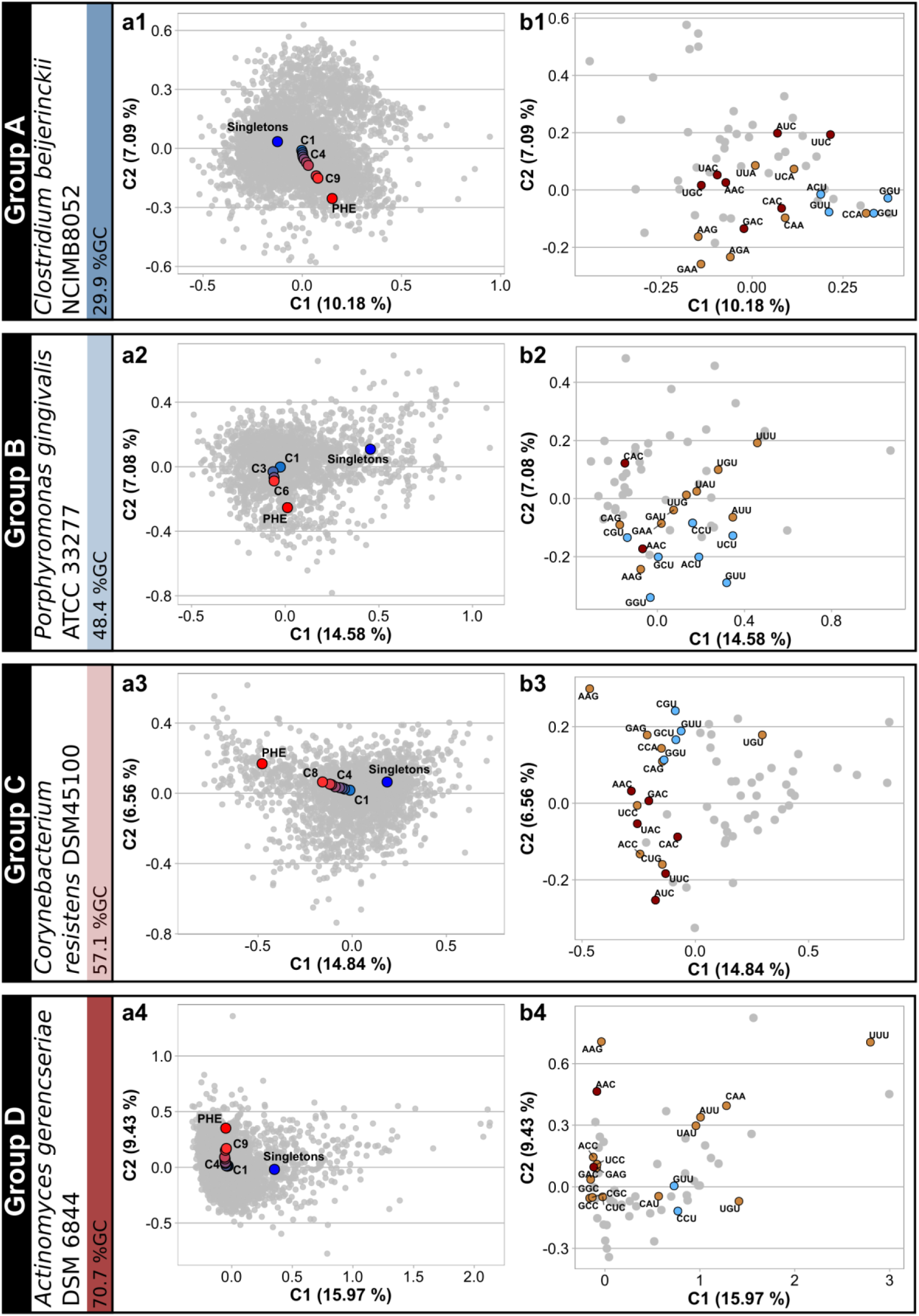
Raw codon counts-based Correspondence Analysis (CA) plots of core-gene sets with different degrees of conservation throughout the phylogeny of selected prokaryote families (Groups A to D). **Panels a1 to a4:** In 4 reference strains with different GC contents, individual genes (in gray) are represented in the space of the first-two CA components, with the percent variation of components 1 and 2 being indicated on the axes. CAs were computed using raw codon counts (RCC) as the input variables. Average coordinates (centroids) for different gene sets (i.e. singletons in blue, C1 to Cn in a gradient from blue to red, and PHE in red) were projected on the CA space. In C1 to Cn the higher number denote a more ancestral core-gene set within the phylogeny. Table S1_a (tab 1) lists the prokaryote species that were used to construct each Ci gene set by means of the EDGAR software (54, 55). **Panel b1 to b4:** Plots describing codon relative weight in the first two principal-component positions of the CA. Codons with the highest CUF enrichment for each amino acid from C1 to PHE (i.e. those codons that better represent translational adaptation) were colored in brown, except when those same codons corresponded also to a C- or to a U-bias in which cases they were colored in red and light blue, respectively.

The general features that characterized the bias in codon usages can be summarized as follows. First, a general pattern indicated that in bacteria from Groups B, C and D the PHE genes are enriched in codons with higher GC3 when compared with singletons (Fig. 1 and Fig. S1, right panels). Conversely, an AU enrichment in the third position of codons was observed in the ancestral core fractions of organisms from Group A which have extremely low GC contents. Second, from C1 to Cn in the CA plot, the codon usages gradually shifted away from the position of the singletons (the unique genes) to approach the region where the PHE genes were located (Fig. 1 and Fig S1, left panels). Similar results were obtained when PHE genes were subtracted from the different Cn cores (see Fig. S2). Thus, the overall evidence suggested that gene ancestry correlated with a codon-usage optimization that resembled the one observed in the PHE genes. Nonetheless, the more ancestral core genes (*i. e.*, the Cn gene sets) never overlapped with the position of the PHE genes in the CA plots. In most prokaryote species, the order of positions in the CA plot followed the sequence singletons-C1-Cn, which series was associated with an enrichment in some of the C-ending 2-/3-fold degenerate codon families (*i. e.*, the 2-/3-fold C-bias; *cf*. the distribution of red circles in Fig. 1 and Fig. S1, right panels); whereas the position of the PHE genes compared to that of Cn was characterized by an additional enrichment in U-ending 4-fold degenerate codon families (*i. e.* the 4-fold U bias; *cf*. the distribution of light-blue circles). Each of the previous effects varied in relative intensity among the different prokaryote families, where other specific codon changes (brown circles in Fig. 1 and Fig. S1) also occurred from C1 to Cn to PHE and accompanied the above-mentioned 2-/3-fold C, and 4-fold U biases. Wald et al. (41) have previously reported that the C and the U biases are associated with an improved codon-usage correspondence to the anticodons of the tRNA pool. The combined effects of the C and U biases are the basis for the “rabbit head” distribution of genes that we observe in most of the CA plots (gray dots), an effect that was originally described in *E. coli* (21). Contrasting with the codon usage of core and PHE genes, the singleton genes tend to be enriched in A/U-ending codons.

### Indication from m-tAI values that the codon usages of more ancestral genes are better adapted to the cellular translational machinery

In order to explore how extensive the correlations between codon usage, gene ancestry, and translation efficiency were, we calculated the m-tAI values for the C1 to the Cn core genes for a given strain and used those indices to estimate the adaptation of each gene set to the tRNA pool. Each m-tAI takes into consideration both the copy number of each tRNA structural gene as an estimation of that tRNA’s cellular concentration and the codon-anticodon interactions including the classical Watson-and-Crick interactions (WCIs) along with the wobble rules (see Materials and Methods). Unfortunately, nonstandard forms of base pairing, such as U:U interactions and others, are not included in the m-tAI calculations. In Fig. 2, the left panels illustrate how with progressive gene ancestry the m-tAI generally increases to often approach that of the PHE genes, thus evidencing that the more ancestral cores are enriched in genes that displayed adaptive—*i. e.*, selection-dependent—changes in their codon usage. That such m-tAI increases with progressive ancestry had been observed in strains from 18 prokaryote families (17 eubacteria, 1 archaea) was indeed remarkable (*cf.* Fig. 2 and Fig. S3, left panels a1, a3, a5, a6, a8, a10, and a12 to a19). In the reference strains from these prokaryote families, the PHE genes (red dashed lines) were always associated with higher m-tAI values than those of the core-gene sets from the same genome. Conversely, singletons (blue-dashed lines) were always the gene sets with the lower m-tAIs, thus suggesting that accessory genes (*i. e.*, those present in plasmids, phages, and the unique genes in chromosomes) involve codon usages that—most likely due to their non-essential character—is far from being optimized with respect to the host-translation machinery. Strains with the characteristics described above have genomes with quite diverse GC contents, ranging from *ca*. 30% to over 70%. Exceptions to the general increase in the m-tAI values with ancestry are likely due to m-tAI deficiencies to quantitate non-standard codon-anticodon interactions (i.e. those different from WCIs, and wobble base pairing) (36).

**Fig. 2.**
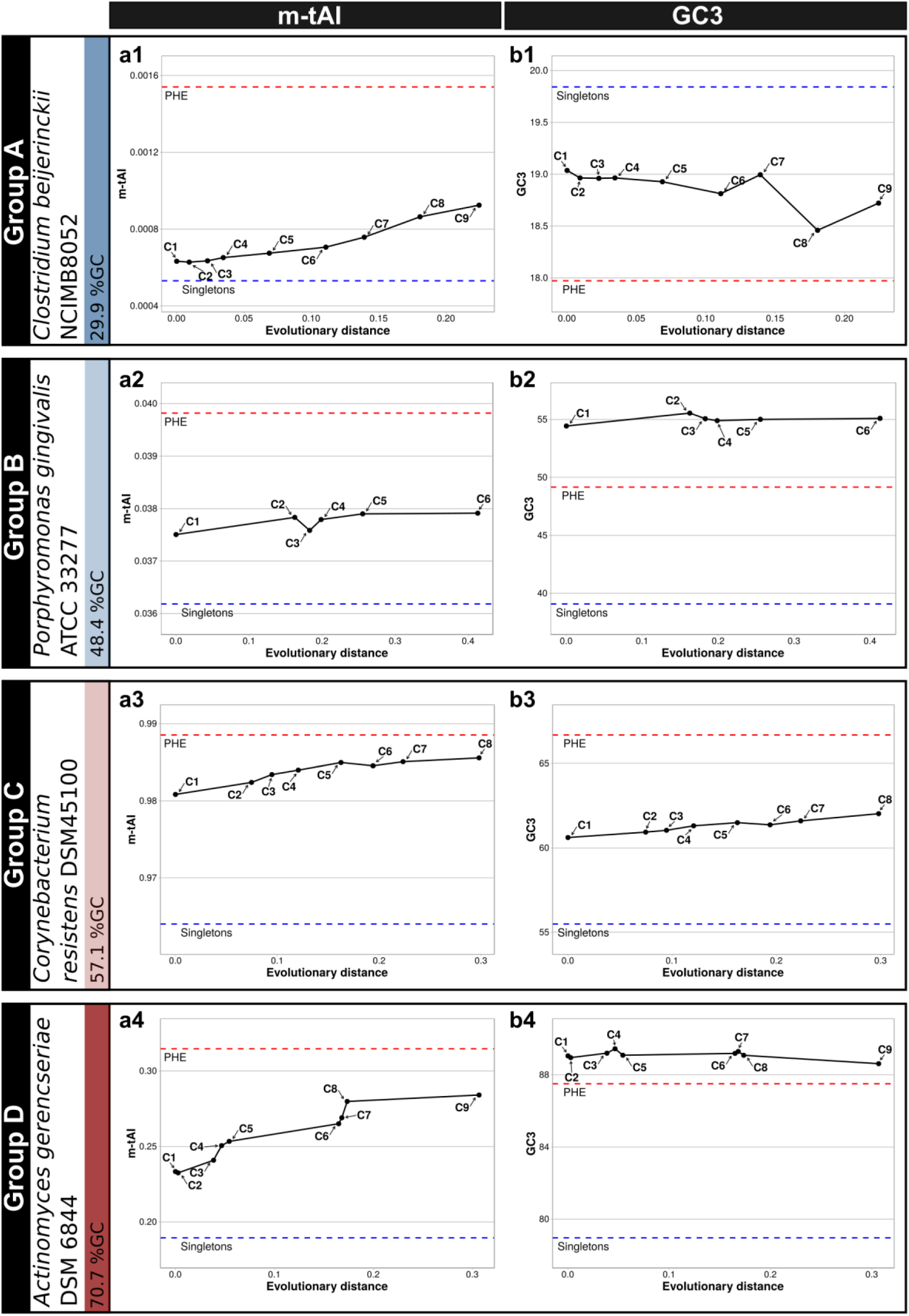
Codon-usage adaptations to the cellular tRNA-pool, and changes in the GC3 content of different prokaryote core genes. The reference strains represented here are the same five as in Fig. 1. **Panels a1 to a4:** In each panel, the modal tRNA-adaptation index (m-tAI) calculated for each of the Ci gene-sets as described in Materials and Methods is plotted on the *ordinate* as a function of the evolutionary distance indicated on the *abscissa* (Table S1_a, tab 2) as inferred from the corresponding phylogenetic trees included in Table S1_a-c. Higher values of m-tAI indicate an enrichment in the codon usage frequencies of those synonymous codons better adapted to the host-cell tRNA pool. The C1 to Cn gene sets plotted here are the same as those presented in Fig. 1. The red and blue horizontal dashed lines correspond to the respective m-tAI values calculated for the PHE genes and the singletons. **Panels b1 to b4:** In each panel, the average GC3 content in each core-gene set of increasing ancestry is plotted on the *ordinate* as a function of the evolutionary distance indicated on the *abscissa* as in panels a1 to a4. The PHE genes and the singletons are represented as red and blue horizontal dashed lines, respectively.

### Effect of codon optimization on the GC content

An analysis of the prokaryote genomes with different GC contents enabled us to explore how the GC composition at the third base of codons (*i. e.*, the GC3) changed in the core-gene sets over ancestry, and to compare the results with the GC3 in PHE genes and singletons. Since the first two positions in codons are constrained by the protein-coding information, most of the GC changes result in variations in synonymous codons (2). As we have seen in the two previous sections, core genes adjust their codon usages in the direction of the PHE genes (Fig. 1 and Fig. S1, left panels) in order to improve translation (Fig. 2 and Fig. S3, left panels). The question thus became raised as to how bacteria with different GC contents changed their GC3 composition in the process of adapting their codon usage. The results presented in Fig. 2 and Fig. S3 (right panels) show that changes in GC3 in genomes from Groups A to D each follows a distinctive pattern from singletons-to-Ci-to-PHE. Whereas in genomes that belong to Group A (overall GC content lower than ca. 35%) the GC3 decreases from singletons to Ci to PHE (*cf*. Fig. 2, panel b1), in the genomes included in Group C the GC3 either increases from singletons to Ci-to PHE (*cf*. Fig. 2, panel b3) or plateaus in Ci to PHE at a high level (*cf*. Fig. S3, panel b17). In contrast, genomes pertaining to Group B; exhibited a biphasic pattern with an initial GC3 increase from the level of the singletons up to the contents of the Ci series (with i varying from 1 to n) followed by a later decrease from the Cn values down to those of the PHE genes (*cf*. Fig. 2, panel b2). Those changes in the Group-B genomes were reflected in pronounced forward and backward movements in the position of the core genes in the CA plots, first from singletons to Ci and then from Cn to the PHE genes (*cf*. Fig. S1, organisms in Group B). A similar biphasic pattern in the CA plots could also be recognized, though softened, in certain species that were included in Group C or even Group D where the PHE genes did not evidence a decrease in GC3 levels when compared to those of the core genes. The genomes in Group D had extremely high global GC contents and had GC3 values in all their core-gene sets (C1 to Cn) that were comparable—though slightly higher—than the corresponding values in their PHE genes. In the next section we will describe how individual codons for a given amino acid are selected in the more ancestral core-gene sets.

### Characterization of codons that improve adaptation to the tRNA pool

The variations in the use of individual codons when progressing from the C1 to the Cn gene sets were analyzed in the different prokaryote genomes, together with the tRNA-gene–copy numbers and the codon-adaptation indices (*Wi*(s); *cf*. Materials and Methods). Figs. 3 and S4 illustrate the CUFs (codon-usage frequencies, *cf*. Materials and Methods) for singletons, PHE genes, and core genes with increasing ancestry together with the tRNA-gene copies and the *Wis* (Fig. S5 summarizes the *Wis*, ΔCn-C1, and ΔPHE-Cn in the different genomes studied). In agreement with previous reports (10), our results demonstrated that the CUF values among synonymous codons were strongly influenced by the global GC content in each genome—*i. e.*, codons with G and C in the 3’ position (N_3_) were the more abundant synonymous codons in the GC-rich genomes, whereas A and U become predominant in that position in the genomes with low GC content (Figs. 3 and S4). An inspection of the proportion of codon usage for each amino acid in ancestral cores compared to the more recent ones (curves in Figs.3, S4, and S5) revealed that in most genomes a C-bias enrichment occurred with increased ancestry at the 3’ position of the 2-fold pyrimidine-ending codons—for Asp (GAU, GAC), Phe (UUU, UUC), His (CAU, CAC), Asn (AAU, AAC), and Tyr (UAU, UAC)—as well as in the unique 3-fold codons for Ile (AUU, AUC, AUA), which three included the pyrimidine-ending pair AUY (Figs. 3, S4, and S5). Corresponding to the observed C bias, in all these examples high *Wi* values (shown in parenthesis in the figure) were observed for the C-ending codons, which triplets were decoded through exact WCIs with the cognate tRNA species (*i. e.*, with the anticodon G_34_N_35_N_36_). Because of the absence of tRNA species bearing anticodons A_34_N_35_N_36_ for these six amino acids, lower *Wi* values were obtained for the U-ending codons as the consequence of a weaker wobble codon-anticodon non-WCI recognition. Especially noteworthy was the observation that, though to a lesser extent, the bacteria with extremely low GC contents likewise exhibited a C bias in the 2- to 3-fold–codon family, irrespective of a global decrease in the GC3 value, as in the example of *Clostridium beijerinckii* (*cf*. Figs. 2 and 3).

**Fig. 3.**
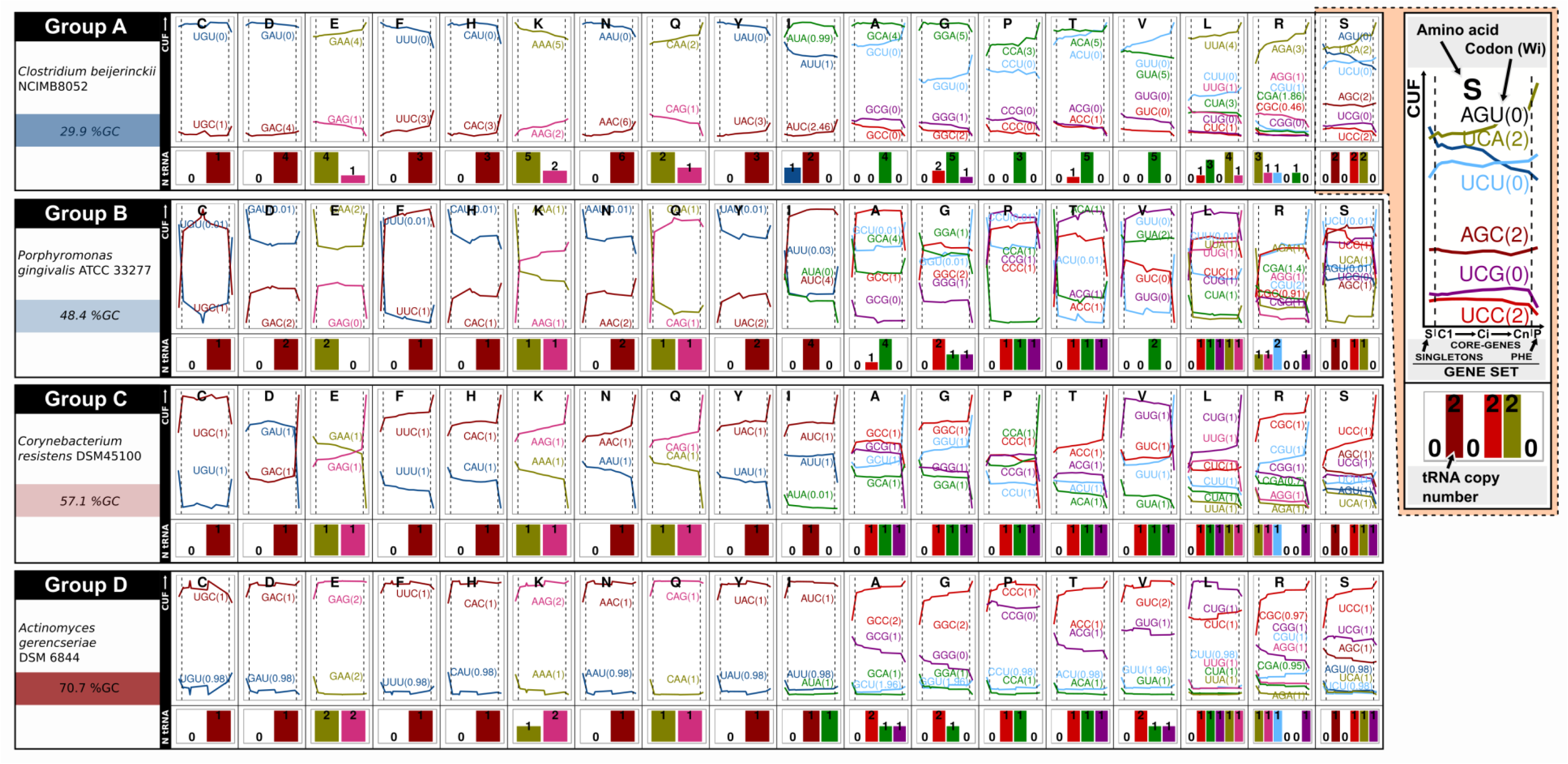
Codon usage frequencies and adaptation indices (*Wi*s) of the gene sets analyzed in this work, together with the tRNA-gene–copy numbers for strains of the four reference Groups A to D. For the amino acid denoted by the corresponding single-letter identification code located above each panel, the change in the modal codon-usage frequencies (CUFs; see Materials and Methods) of the core-gene sets with increasing ancestries (left to right, the C1 to Cn), the PHE genes, and the singletons are plotted in the upper panels as solid horizontal curves for each of the indicated codon triplets between the two vertical broken lines, for the singletons to the left of the first of those lines, and for the PHE genes to the right of the second (with singletons and PHE genes being located at the beginning and the end of the curves, respectively). The CUFs are represented by different colors with the associated absolute codon-adaptation index (*Wi*, (36)) being denoted within parentheses beside each triplet. Finally, the presence and gene-copy number (N tRNA) of the cognate tRNA species of a given synonymous codon bearing the exact complementary anticodon is depicted with a number and a bar of proportional height in the lower panel in the same color as the corresponding triplet and curve in the upper panel.

In the instance of the 2-fold purine-ending codons—that is GAA and GAG for Glu, AAA and AAG for Lys, and CAA and CAG for Gln—we observed that the codons with G or A in the 3’ position were enriched from C1 to Cn and from Cn to PHE (*i. e.*, ΔCn-C1 and/or ΔPHE-Cn in Fig. S5, respectively) depending upon which tRNA species (anticodons) were present. In those examples where only the tRNAs bearing the U_34_N_35_N_36_ anticodons were present, the cognate A-ending codons recognized by WCIs were the ones that became enriched in the more ancestral core and/or PHE genes (*cf*. in Fig. S5, the GA<u>A</u> triplet for Glu in *C. violaceum*, *P. graminis*, *Bacillus subtilis*, *Bordetella holmesii*, and *Leisingera methylohalidivorans*; the AA<u>A</u> for Lys in *M. smithii* and *Bacillus subtilis*; and the CA<u>A</u> for Gln in *M. smithii, Streptococcus equii*, and *B. subtilis*). Accordingly, these 3’-A–ending codons were associated with higher *Wi* values than the corresponding codons ending in G, as the latter were recognized only by wobble-base pairing (*i. e.*, G_3_-U_34_ interaction). In a second circumstance, where both tRNA species for the same amino acid (*i. e.*, those bearing anticodons U_34_N_35_N_36_ or C_34_N_35_N_36_) were present, a more frequent enrichment in G–ending codons was observed (with few exceptions) since such codons can be decoded by either Watson-Crick or wobble interactions with the tRNA anticodons C_34_N_35_N_36_ or U_34_N_35_N_36_, respectively. In those few examples where the A-ending were more enriched than the G-ending codons, a higher copy number of the tRNA genes was always observed with anticodons U_34_N_35_N_36_ than that obtained with the anticodons C_34_N_35_N_36_ (*cf*. in Figs. 3 and S4, the GA<u>A</u> triplets for Glu in *Bacteroides vulgatus* and *C. beijerinckii*; the AA<u>A</u> triplets for Lys in *S. multivorans*; and the CA<u>A</u> triplets for Gln in *C. beijerinckii* and *S. multivorans*).

A different codon usage bias—in a pattern not found in the 2-/3-fold–degenerate amino acids—was observed in codons encoded by 4-fold–degenerate amino acids (Val, Thr, Pro, Gly, Ala) or by the 4-fold boxes of the 6-fold degenerate amino acids (Ser, Leu, Arg). In these 4-fold groups a U-bias enrichment (*i. e.*, a NNU-codon enrichment) was observed in the PHE genes from most of the genomes irrespective of their GC content (Figs. 3, S4, and Fig. S5). This enrichment in U-ending codons, previously reported as a U bias (41), could not be explained by WCIs with A_34_N_35_N_36_ tRNAs because these latter species were not present in prokaryotes, except in the example of Arg. The observed U bias likely occurred through the previously proposed nonconventional codon-U_3_:anticodon-U_34_ interaction that was known to exist in bacteria (64). The presence of U_34_N_35_N_36_ tRNA species might then lead to an increase in both NNA and NNU codons as a consequence of positive WCIs and U_3_-U_34_ interactions, respectively.

All the codon adaptations that we have described in this section referring to core genes proved to be more prominent in the PHE genes, whose triplets were even better adapted to the translational machinery. Contrasting with such a strong pattern of selection-associated codon bias, the singletons displayed codon usages that were in general the most distant from those observed in the PHE genes (as exemplified in the CUFs in Figs. 3, S4, and in the CA plots from Figs. 1 and S1). These observations are also in agreement with variations in the m-tAIs for the different gene sets presented in the previous section.

### Search for coding signatures for translation efficiency and accuracy: Codon-usage profiles associated with sequences encoding highly-expressed–variable (HEP_*vr*) and -conserved (HEP_*cr*) translated domains

Expression level and amino-acid–sequence conservation are both parameters that positively correlate with codon-usage optimization (65). Nevertheless, the relative relevance of efficiency and accuracy to translation plus the way in which either one of those parameters affects the other have not yet been investigated in detail. A central limitation that made such studies difficult was associated with the natural genomic heterogeneity in gene ancestry along with the expression level and the sequence conservation (structural constraints) in the translated products. In order to reduce the degrees of freedom in the analysis; for each of six different bacterial species, we created two distinct gene sets based on the experimental proteome data. One of those gene sets consisted of genes encoding proteins with the highest expression levels in the cell (*i. e.*, the HEP), while the other was associated with proteins with low cellular abundance (*i. e.*, the LEP). Then, the conserved (*cr*) and variable (*vr*) sequences among the orthologs were collected from each individual gene (*cf*. Materials and Methods), the corresponding HEP_*cr*, HEP_*vr*, LEP_*cr*, and LEP_*vr* modal codon usages were used to calculate the relative distances illustrated in the neighbor-joining tree presented in Fig. 4. In five out of the six species present in the trees (Fig. 4A to 4E), the HEP_*cr* and HEP_*vr* sequences separated from those of the singletons, the core genes, and all the LEPs as the result of a strong codon-usage adaptation (also reflected in the low effective number of codons (Ncs) associated with the HEPs, indicated in parentheses following labels in the tree). Furthermore, the large distance in the tree between HEP_*cr* and LEP_*cr* (where both sequences encode regions with conserved amino acids, but with different expression levels) compared to the much shorter distance between HEP_*cr* and HEP_*vr* (where both encode highly expressed products with different degrees of conservation) pointed to the quantitatively stronger effect of efficiency over accuracy in shaping codon-usage bias. However, codons that were optimized as a result of accuracy under high and under low expression—*i. e.*, [HEP_*cr*–HEP_*vr*] and [LEP_*cr*–LEP_*vr*], respectively, labelled ***A*** for accuracy at the bottom of Fig. 5—were highly coincident with the codons that were optimized through efficiency—*i. e.*, [HEP_*cr*–LEP_*cr*] and [HEP_*vr*–LEP_*vr*], labelled ***E*** for efficiency. In some organisms, the greater distance between HEP_*cr* and HEP_*vr* than between LEP_*cr* and LEP_*vr* (Fig. 4) indicates a stronger influence of accuracy in codon-usage optimization when operating under high-expression conditions, thus pointing to an interaction between the simultaneous requirements of high fidelity and efficiency. The more relevant contributions to the global difference in codon usage between HEP and LEP resulted to be efficiency (both in conserved and in variable regions) (***E*** columns in Fig. 5) followed by accuracy under high expression (first ***A*** column in Fig 5)(the stronger the contribution of each factor, either ***E*** or ***A***, the shorter the distance in brackets to HEP-LEP in the figure). The heat maps display the complete profiles of preferred codons for sequences requiring high translational accuracy and/or efficiency (protein demands). As expected, the preferred codon for each amino was in agreement with the C and U bias and the tRNA-copy number described in the previous sections. In light of these results, the highly-expressed variable and conserved domains (HEP-*vr*/*cr*) constitute the basis for explaining the observed codon-usage optimization in the more ancestral core-gene sets (Cn), which concentrate HEPs (Table S3). Fig. 6 illustrates that HEP sequences (red dots) are those under the highest selective pressure to optimize codon usage because of both their expression level and their degree of conservation.

**Fig. 4.**
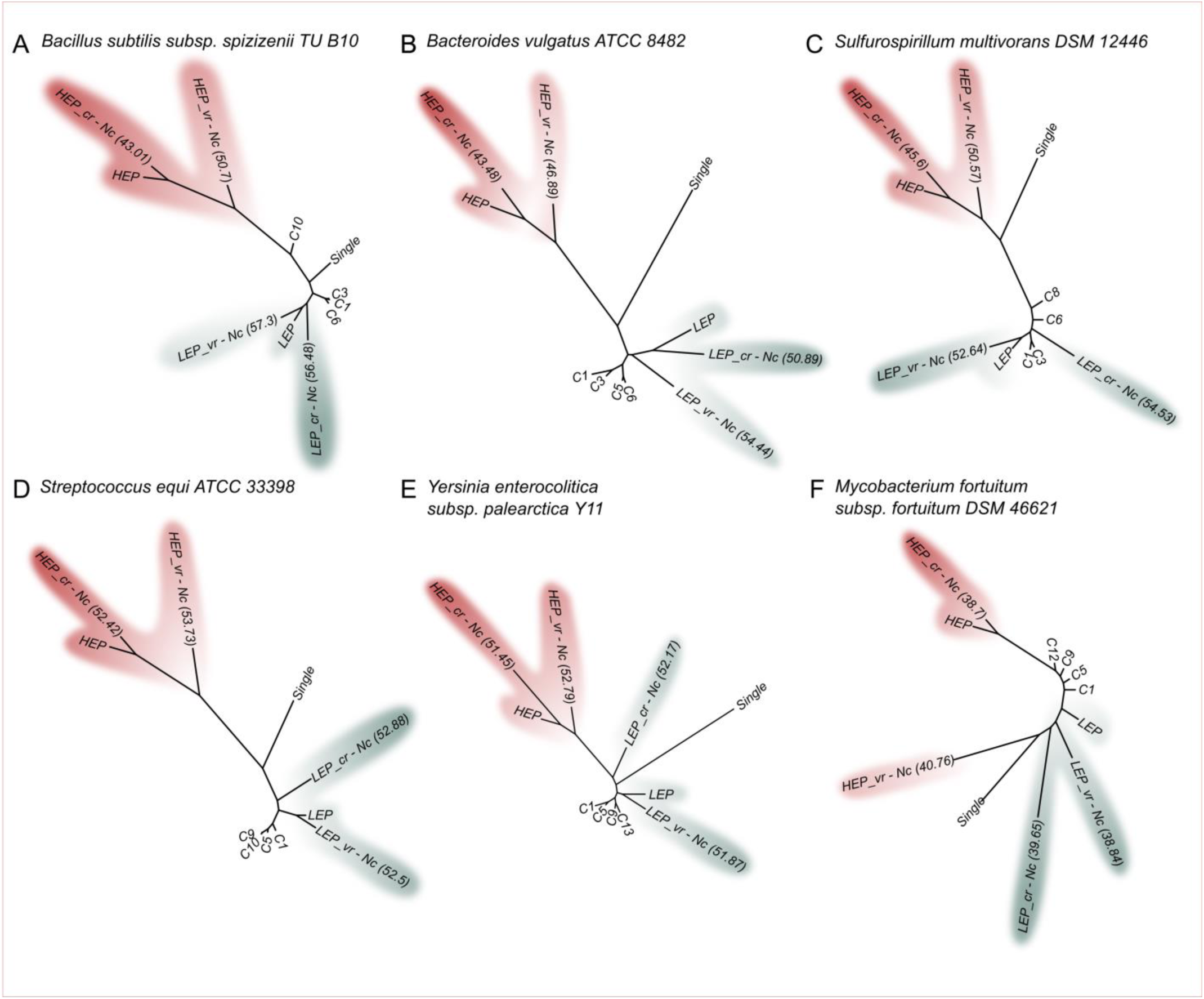
Neighbor-joining–distance trees of different gene sets encoding HEP, LEP, and their associated conserved (*cr*) and variable (*vr*) regions based on the corresponding modal codon usage. Modal codon usage-based neighbor-joining trees were constructed for the indicated gene sets and intragenic regions (*cr* and *vr*) following the method described by Karberg et al. (17) along with the neighbor-joining program of the Phylip package (69). Phylogenetic trees were drawn through the use of the Figtree application (70). The figure abbreviations are as follows: C1 to Ci, core-gene sets with increasing ancestry; single, singletons; HEP, genes encoding proteins with the highest expression level; LEP, genes encoding proteins with the lowest expression level; HEP_*cr*, conserved HEP sequences (dark red); HEP_*vr*, variable HEP sequences (light red); LEP_*cr*, conserved LEP sequences (dark blue); and LEP_*vr*, variable LEP sequences (light blue). HEP and LEP *cr* and *vr* subfractions were recovered as indicated in Materials and Methods through the use of the polypeptide sequences included in C13 for *Yersinia enterocolitica* subsp. *palearctica* Y11, C10 for *Streptococcus equi* ATCC 33398, C8 for *Sulfurospirillum multivorans* DSM 12446, C9 for *Bacillus subtilis* subsp. *spizizenii* TU B 10, C6 for *Bacteroides vulgatus* ATCC 8482, and C12 for *Mycobacterium fortuitum* subsp. *fortuitum* DSM 46621 (ATCC 6841). The effective number of codons (Nc_s_) as previously defined by Wright (71) are indicated in brackets for the *cr* and *vr* subset of sequences.

**Fig. 5.**
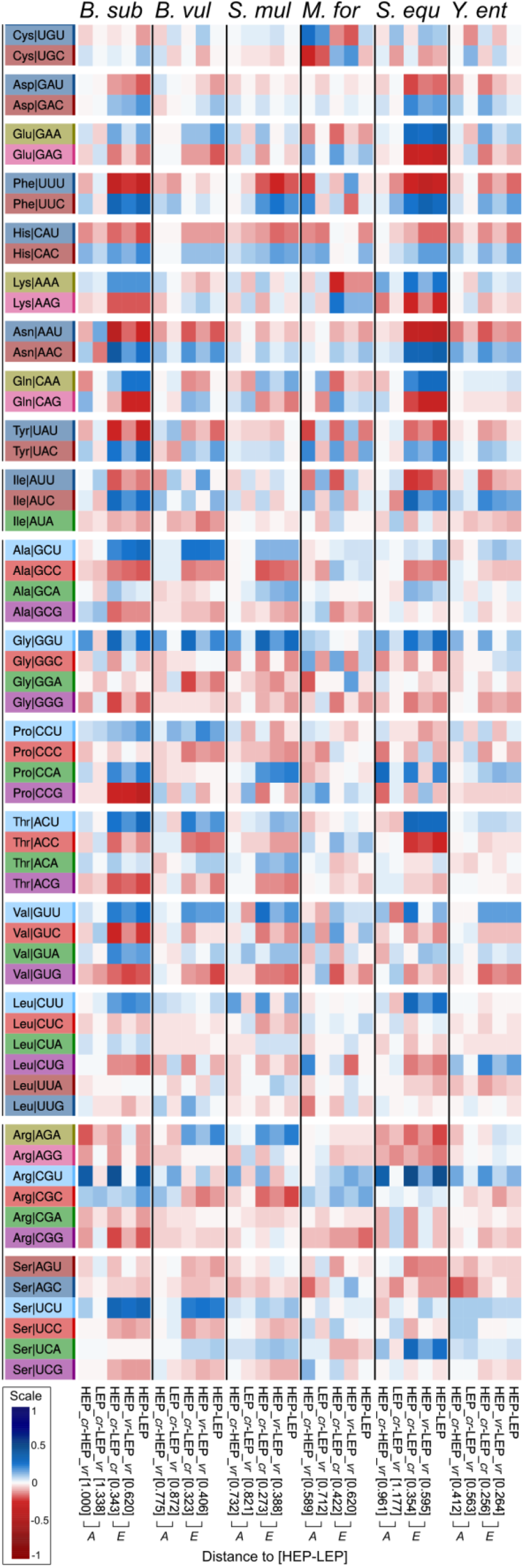
Heat-map representation expressing differences in modal-codon-usage profiles between the indicated gene sets. The color scale from red to blue indicates the relative level of use of each particular codon in a gene set compared to that of another (*i. e.*, gene set 1 *versus* gene set 2). The blue color corresponds to the dominant use of a particular codon in Gene set 1 over the use of the same codon in Gene set 2 (and *vice versa* for the red color). Amino acids are indicated in the standard three-letter code. The heat map was constructed through the use of the phytools R package (72). The abbreviations are as follows: HEP, genes encoding proteins with the highest expression level; LEP, genes encoding proteins with the lowest expression level; HEP_*cr*, conserved HEP sequences; HEP_*vr*, variable HEP sequences; LEP_*cr*, conserved LEP sequences; and LEP_*vr*, variable LEP sequences. The [HEP (gene set 1) – LEP (gene set 2)] column represents the profile of the optimized codons when comparing the coding strategies in high- *versus* low-expression genes (*i. e.*, reflecting differences in their modal codon usages). The columns indicated by ***A*** correspond to the profiles of codons optimized as a result of accuracy (*i. e.*, differences between [HEP_*cr* – HEP_*vr*] and [LEP_*cr* – LEP_*vr*]). The columns indicated by ***E*** correspond to the profiles of optimized codons through high expression (*i. e.*, reflecting differences in efficiency between [HEP_cr – LEP_*cr*] and [HEP_*vr* – LEP_*vr*]). The numbers in brackets indicate the extent to which changes induced by either efficiency or accuracy approach the differences in codon usage between HEP and LEP (*i. e.*, distances from each column to the column HEP – LEP). The shorter the distance in brackets the stronger the contribution of the indicated factor (*i. e.*, A = accuracy or E = expression level) to codon optimization in the HEP.

**Fig. 6.**
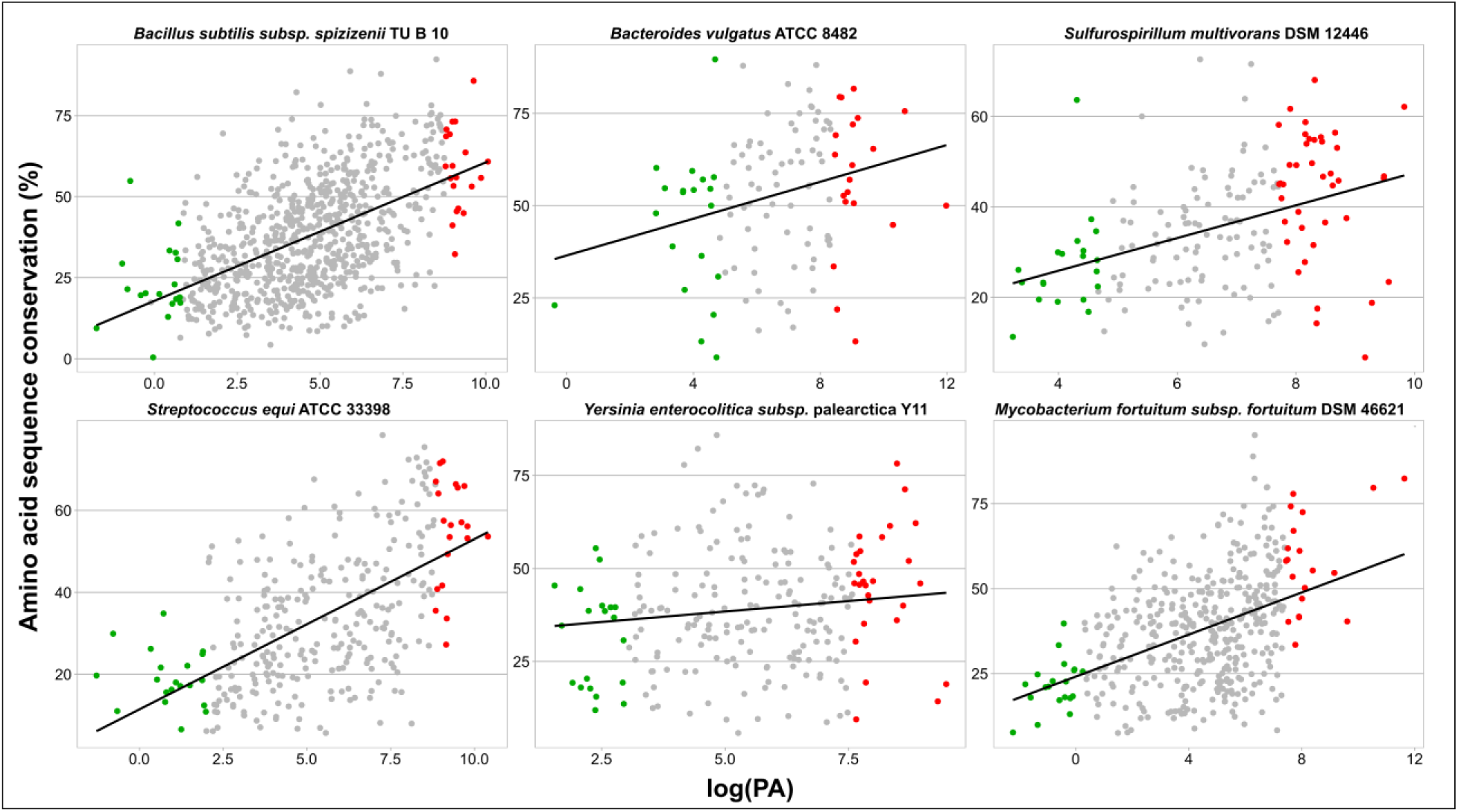
Amino-acid–sequence conservation in proteins with different cellular abundances. The amino-acid–sequence conservation calculated for proteins of the indicated bacterial species and core fractions (Materials and Methods) are plotted on the *ordinate* as a function of the logarithm of the corresponding protein abundance (logPA) on the *abscissa*. The red and blue dots correspond to HEP and LEP, respectively, with all the other proteins of the same core represented in gray. The linear regressions and graphs were all performed with the ggplot2 library from the R package.

## DISCUSSION

Since gene adaptation to a host cell is expected to be associated with an improved codon selection for translation efficiency and accuracy (43, 66), we investigated correlations between core-gene ancestry and their modal codon usage within a given prokaryote family. In order to ascertain if the adaptation of the more ancestral core genes was an extensive phenomenon among prokaryotes, we analyzed core modal codon usages in 27 different species of Bacteria and 2 of Archaea. That in the CA plots the more ancestral core genes had been the ones with the closest location to the PHE genes in all families was remarkable and strongly indicated a core codon-usage adaptation that likely operated to improve translation. In agreement with the position of the different gene sets in the CA plots, the m-tAI values served to confirm that the PHE genes were the best adapted gene set, followed by the Cn to C1 core genes, in that order, and finally by the singletons, with those being the least adapted genes with the lowest m-tAIs in the genome. Studies made by others have previously demonstrated that the level of gene expression together with the need to preserve accuracy during the translation of conserved amino-acid regions are both among the main parameters that govern codon-usage selection (65). The bioinformatics isolation of conserved (*cr*) and variable (*vr*) coding-sequence domains from genes under high- (HEP) and low-expression (LEP) regimes served in this work to ascertain quantitatively the relative contribution of efficiency (expression level) *versus* accuracy during the selection-based codon-usage optimization. According to the observed neighbor-joining distances (Table S3 worksheet “distances”, and tree in Fig. 4), changes in codon usage derived from differences in gene-expression levels—*i. e.*, the efficiency in terms of the distance from the LEP to the HEP— were between 1.25 to 2.35 times greater than the changes in codon usage resulting in increased accuracy—*i. e.*, the distance from *vr* to *cr*—. The increasing proportion of highly expressed-variable and specially-conserved sequences (*i. e.*, HEP_*vr* and HEP_*cr*) in the more ancestral gene sets constituted the basis for explaining the corresponding high degree of codon-usage optimization that gradually increased from C1 to Cn.

The central question therefore was how adaptive changes in codon usage—which alterations become reflected in m-tAI values—occurred in prokaryotes with quite diverse GC contents (10). Because of the small amount of intergenic DNA in prokaryotes, genomic differences in base composition must mainly derive from changes in the coding regions. Within the alterations in the open reading frames, changes in GC are preferentially associated with modifications in the GC3, and only to a lower extent with alterations in the GC content of the first two codon positions (2, 4). How mutational bias (12) competes with selection (15) to drive all these changes is not yet fully understood. The codon-usage biases described here were associated with the four different patterns of GC3 changes summarized in the schemes presented in Fig. 7 (*i. e.* the genome Groups A, B, C, and D depicted in the figure). The Group-A genomes, those having an extremely low GC content and with their GC3 frequency decreasing from C1 to Cn, proved to have only the tRNA-U_34_ to recognize 4-fold synonymous codons in one or more amino acids. In such instances, the observed core-gene AT enrichment over ancestry appeared to be directly affected by selection (as with the PHE genes), where codons NNA (via WCIs with the tRNA-U_34_) and NNU (via nonconventional U-U interactions) were preferentially enriched over NNC and NNG codons. Though both of those changes were probably related to improvements in translation efficiency, such increases are not always reflected in the m-tAIs since, as stated earlier, U-U interactions are not considered in the calculation of that index. Unfortunately, when we (not shown) and others (37) have attempted to improve the tRNA-adaptation index by including additional nonstandard base pairings, we obtained no better results. Nonetheless, under the assumption that the PHE genes are the best adapted to the translational machinery, in genomes with extremely low GC content—such as those belonging to Group-A—the observed AT3 enrichment from C1 to Cn to PHE (Fig. 7, right panel) should mainly result from selection. According to Hildebrand et al. (15), the mutational processes in very low-GC organisms favor a GC3 enrichment. That the core and PHE genes in bacteria that belonged to Group A had been selected to bear lower GC3 values than singletons in order to improve translation in view of the previous pattern of increasing GC content was remarkable, with this circumstance being a result of the above-mentioned enrichment in NNA and NNU triplets compared to their proportion in the synonymous codons (Fig. 7, right panel). In Group-B genomes, the biphasic pattern observed from singletons to PHE genes could be explained by an initial increase in GC3-rich codons from singletons to core genes, followed by a later U bias from core genes to PHE genes. That initial GC3 enrichment followed by a U3 increase was sufficient to explain the basis of the previously reported “rabbit head” distribution of codon usages that characterizes most prokaryote genomes (21, 67). What should be also especially noted is that the PHE genes separated from the Cn (in both the CA, and the GC3 plots) because of a much more intense U bias likely associated with the difference in expression levels between the two gene sets. In the type-C genomes, in which the GC3 always increased, the absence of a strong U bias from the Cn to the PHE genes led to a less pronounced—*i. e.*, more linear—“rabbit-head” distribution of genes in the CA plot. In addition to that general trend, *Yersinia enterocolítica*, *Methanolacinia petrolearia*, and *Sphingomonas parapaucimobilis* could be considered as having an intermediate behavior between the bacteria in Group C and those in Group B. Finally, the Group-D genomes, which had extremely-high GC contents, were the most restricted with respect to GC3 variations. The quite small compositional variations in that group of genomes became apparent in the compacted location of the different core and PHE genes in the CA plots. What was remarkable is that in Group-D genomes a U bias (though much less intense than in the genomes of Groups A, B, and C) was still a visible variable that differentiated codon usages between the core and the PHE genes. As stated above, the noninclusion of U:U interactions in the m-tAI calculation limited the use of this parameter to expresses the translational adaptation of those gene sets in which a U bias was dominant. Pouyet et al (11) present a model to predict and separate the relative contribution of mutational bias (N-layer), codon selection (C-layer), and amino acid composition (A-layer) on the global GC and the GC3 content. Our analysis is fully consistent with the results reported by Pouyet et al (11) where the C-layer (codon selection / translational selection) has a stronger influence on the GC3 of genes with low effective number of codons (Nc)(such as Cn and PHE) compared to the influence on genes with the highest Nc (such as the C1).

**Fig. 7.**
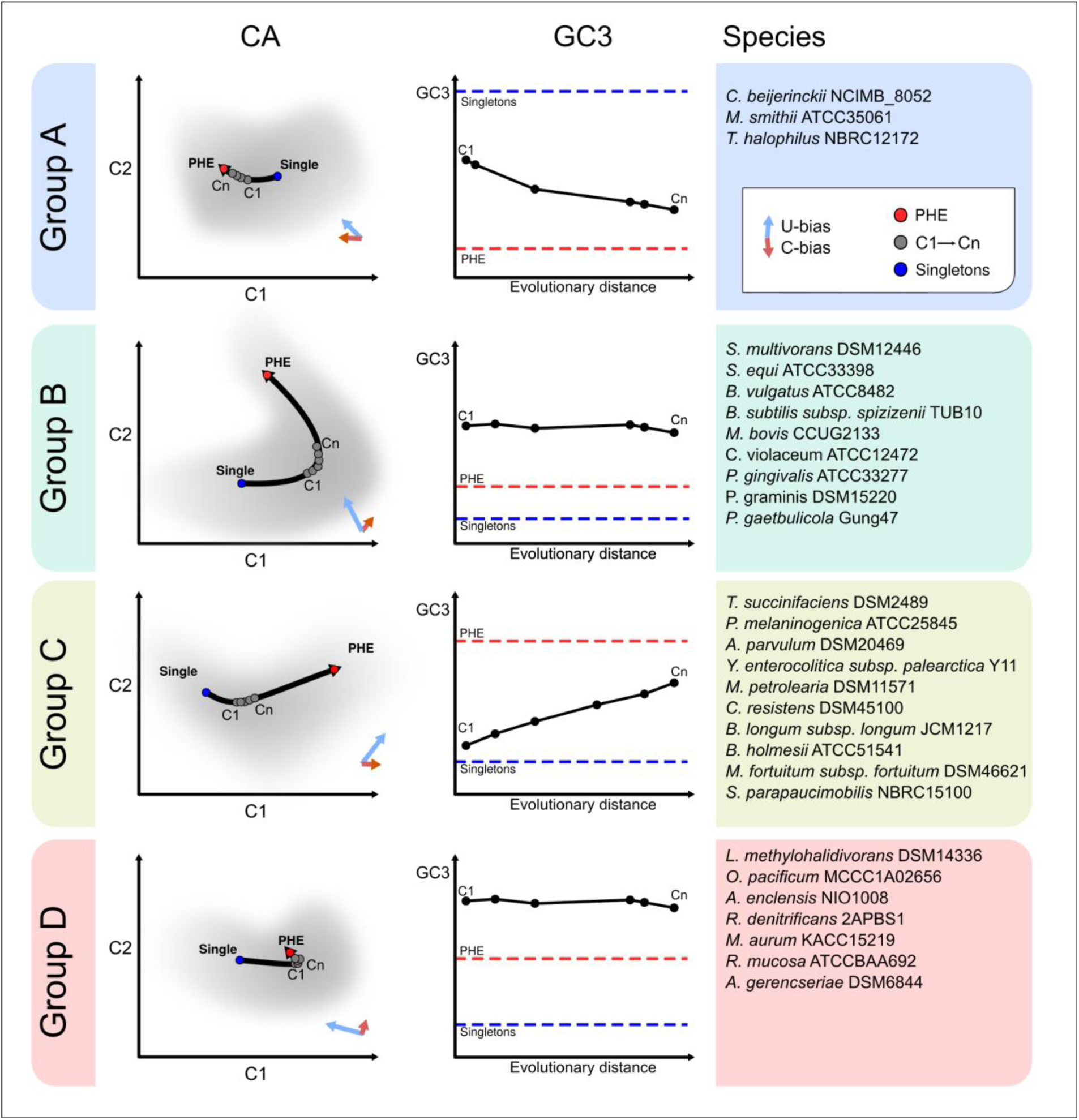
Schematic representation of general codon-usage patterns observed in different prokaryote families. For the prokaryote strains whose genomes were classified as belonging to Groups A, B, C, and D and listed to the right of each panel, cartoons with the associated correspondence-analysis (CA) and GC3-variation pattern among the core-gene sets of increasing ancestry (light gray) are presented, along with the corresponding singletons (blue) and PHE genes (red). The light-blue arrow indicates the direction of the U bias and the red arrow that of the C bias. The right panel is a plot of the GC3 content on the *ordinate* as a function of increasing evolutionary distance on the *abscissa* with the red horizontal dashed line denoting the PHE genes and the blue the singletons.

The results presented here together with previous evidence from other authors have enabled a comprehensive analysis of the principal basis underlying the changes associated with the optimization of codon usage in prokaryotes in different gene sets and in organisms with different GC contents. As stated previously, the overall codon usage is known to be constrained by genome-wide mutational processes (7, 8, 10) that are considered as a main force in shaping the global GC content. The intragenomic codon usage will concurrently become accommodated through selection-driven processes, as has also been extensively reported (35, 42, 48, 68). In order to further our knowledge of the relevance of those factors/forces generating intragenomic variations, we investigated the different nucleotide-base changes underlying the selection of preferred codons in the core and PHE genes of representative prokaryote species. The analysis of gene sets with different expression levels and degrees of conservation in organisms with diverse global GC contents enabled a description of how core codon usage approaches that existing in the PHE genes and how nucleotide changes correlate with an improved compatibility between the genes and the coexisting tRNA pool. That C- and U-ending codons in 2-/3- fold- and in 4-fold–degenerate amino acids, respectively, were specifically enriched as a result of selection to improve translation has been previously reported for different prokaryotic genomes (41). Using separate analyses focused on different gene sets, we demonstrated here that similar selection-driven adaptations in codon usage has taken place from singletons to core genes to PHE genes. The intensity and relevance of the C and U bias was dependent on the particular genome—and especially on the genomic GC content—as well as on the gene fractions under consideration. In contrast to the codon-usage variations occasioned by selection in the core and PHE genes, the singletons constituted the gene set characterized by both the lowest GC3 content as a result of the AT mutation that is universally biased in prokaryotes (12) and a much more relaxed selection than that of the more ancestral genes, with the sole exception of the extremely low-GC–containing genomes of Group A. In addition to a description of the basic changes that together conform the intracellular-codon–usage variations, further investigation should be focused on the analysis of the time course required by the newly acquired information to be properly incorporated into the genetic language of the host cell (codon usage tuning).

## Supporting information

Suplemental Table 1 - part_a

Suplemental Table 1 - part_b

Suplemental Table 1 - part_c

## ACKNOWLEDGEMENTS

We are grateful to Paula Giménez, Silvana Tongiani (both members of CPA CONICET at IBBM), and to Ruben Bustos from UNLP for their technical assistance; and to Donald F. Haggerty for editing the final version of the manuscript. This research was supported by the National Science and Technology Research Council (Consejo Nacional de Investigaciones Científicas y Técnicas – CONICET, Argentina) PIP 2014-0420; the Ministry of Science Technology and Productive Innovation (Ministerio de Ciencia Tecnolología e Innovación Productiva – MinCyT, Argentina) PICT-2012-1719, PICT-2015-2452, and PICT-2017-2022; and CYTED (Ciencia y Tecnología para el Desarrollo) acción 115RT0492. JLL, MJL, and MLF were supported by CONICET, and AL was supported by CONICET and by the UNLP (Universidad Nacional de La Plata).

