## Supplementary material for "Codon-usage optimization in the prokaryotic tree of life: How synonymous codons are differentially selected in sequence domains with different expression levels and degrees of conservation": Suplemental Figure 1

#### FIGURE S1A

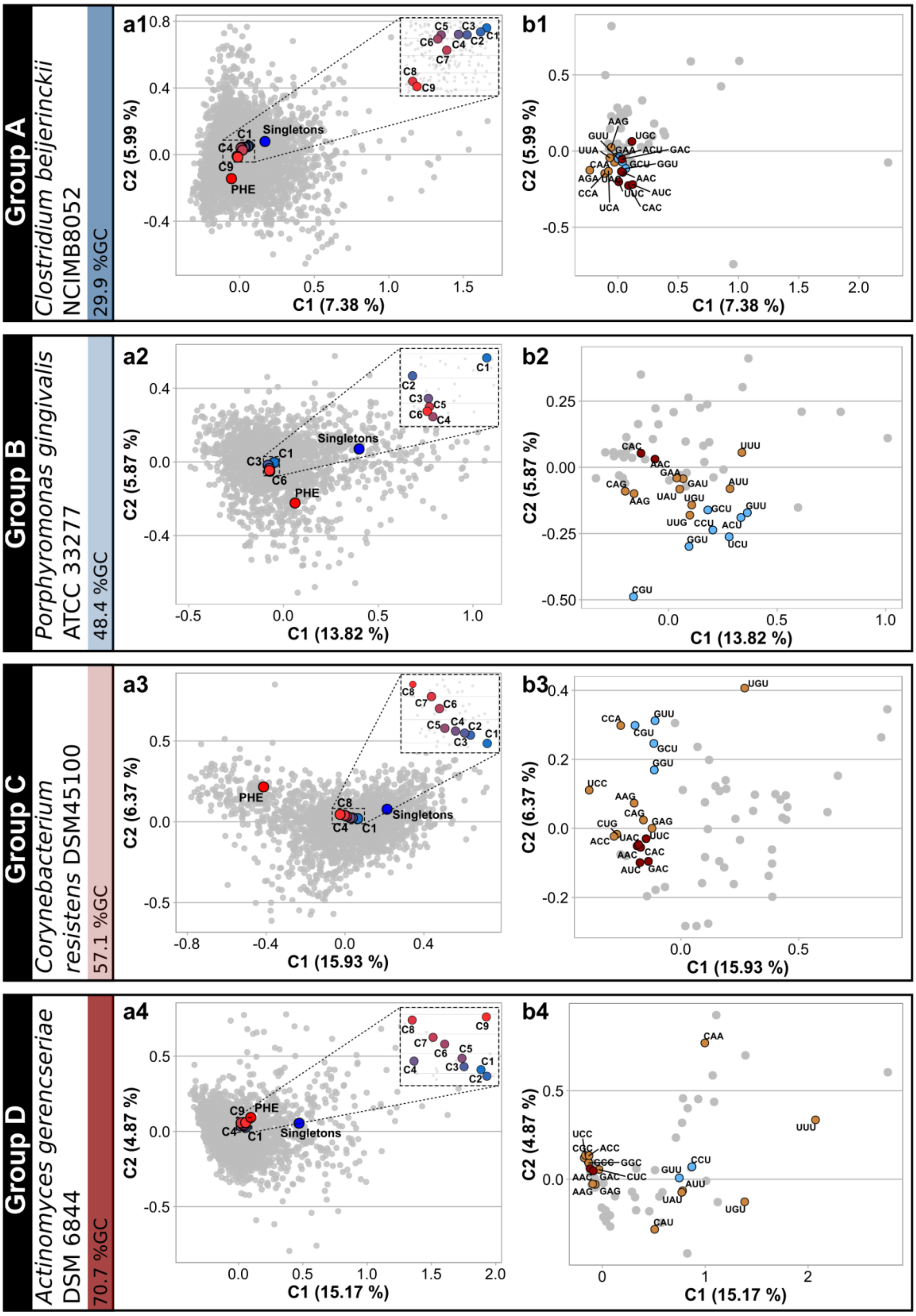


**Fig. S1A. RSCU-based Correspondence Analysis (CA) plots of core-gene sets with different degrees of conservation throughout the phylogeny of selected**

**prokaryote families (Groups A to D). Panels a1 to a4:** In 4 reference strains with different GC contents, individual genes (in gray) are represented in the space of the first-two CA components, with the percent variation of components 1 and 2 being indicated on the axes. CAs were computed using RSCUs as the input variables. Modal codon usages—calculated as indicated in Materials and Methods—for different gene sets (i.e. singletons in blue, C1 to Cn in a gradient from blue to red, and PHE in red) were projected on the CA space. In C1 to Cn the higher number denote a more ancestral core-gene set within the phylogeny. **Panel b1 to b4.** Plots describing codon relative weight in the first two principal-component positions of the CA. Codons that are colored in brown indicate those which present—for each amino acid—the highest CUF enrichment from C1 to PHE (i.e. those codons that better represent translational adaptation) except when some of those codons also correspond to a C-bias (colored in red) or a U-bias (colored in light blue).

### Fig. S1-B

**GROUP A**
*Methanobrevibacter smithii* ATCC 35061

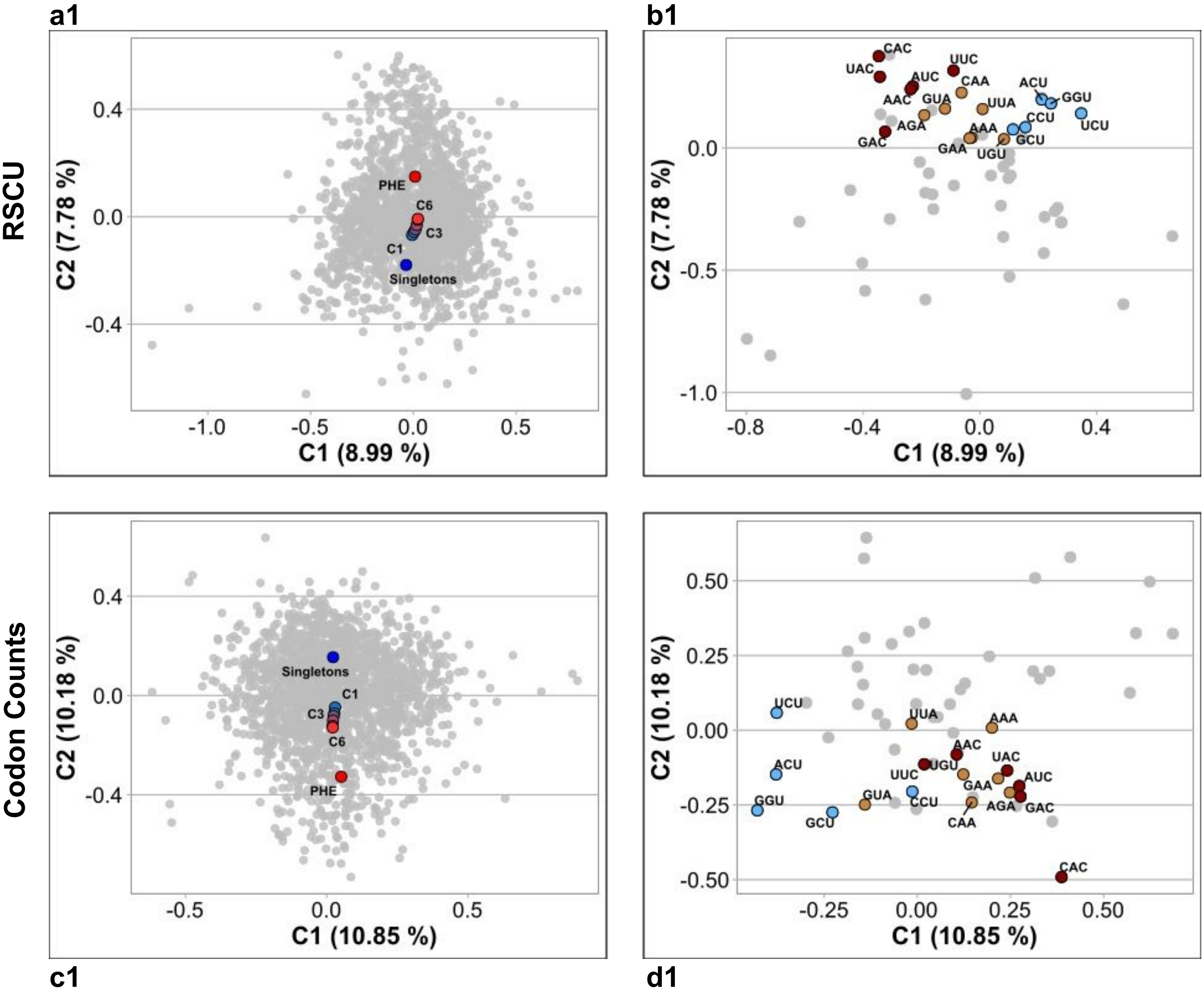

*Tetragenococcus halophilus* NBRC 12172 NC_016052

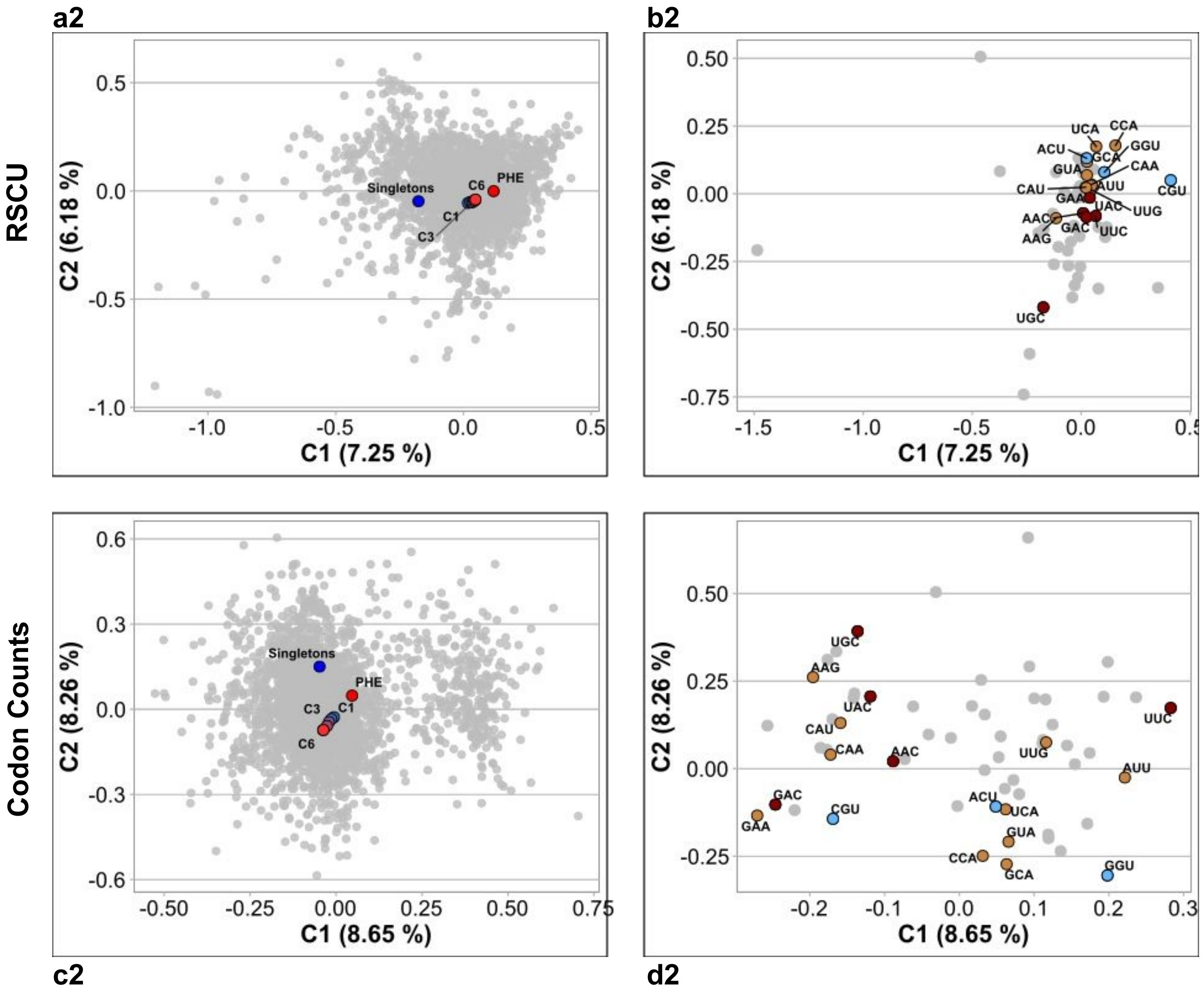

**GROUP B**

*Sulfurospirillum multivorans* DSM 12446 NZ_CP007201

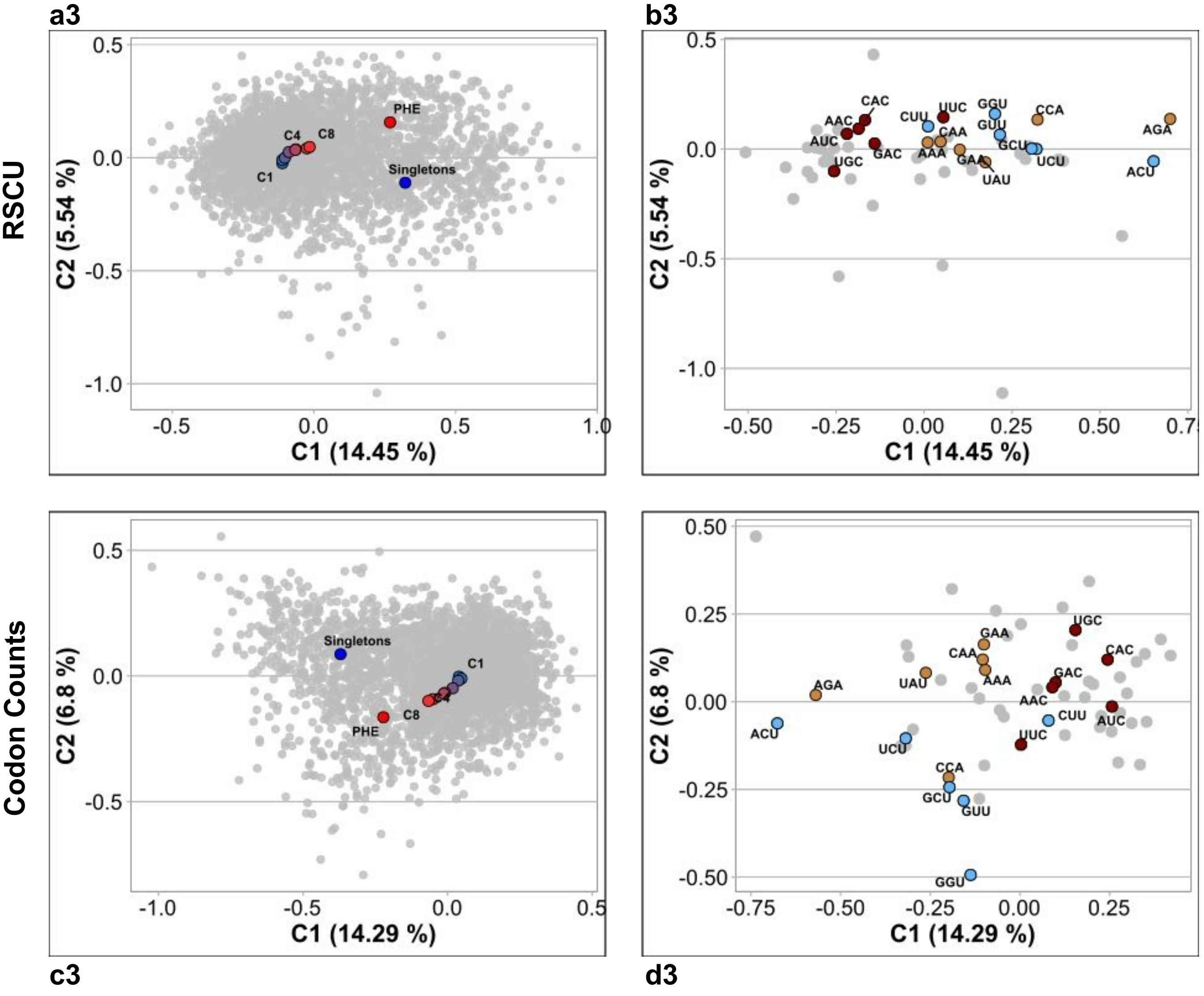

*Streptococcus equi* ATCC 33398 NZ_FTNH01000046

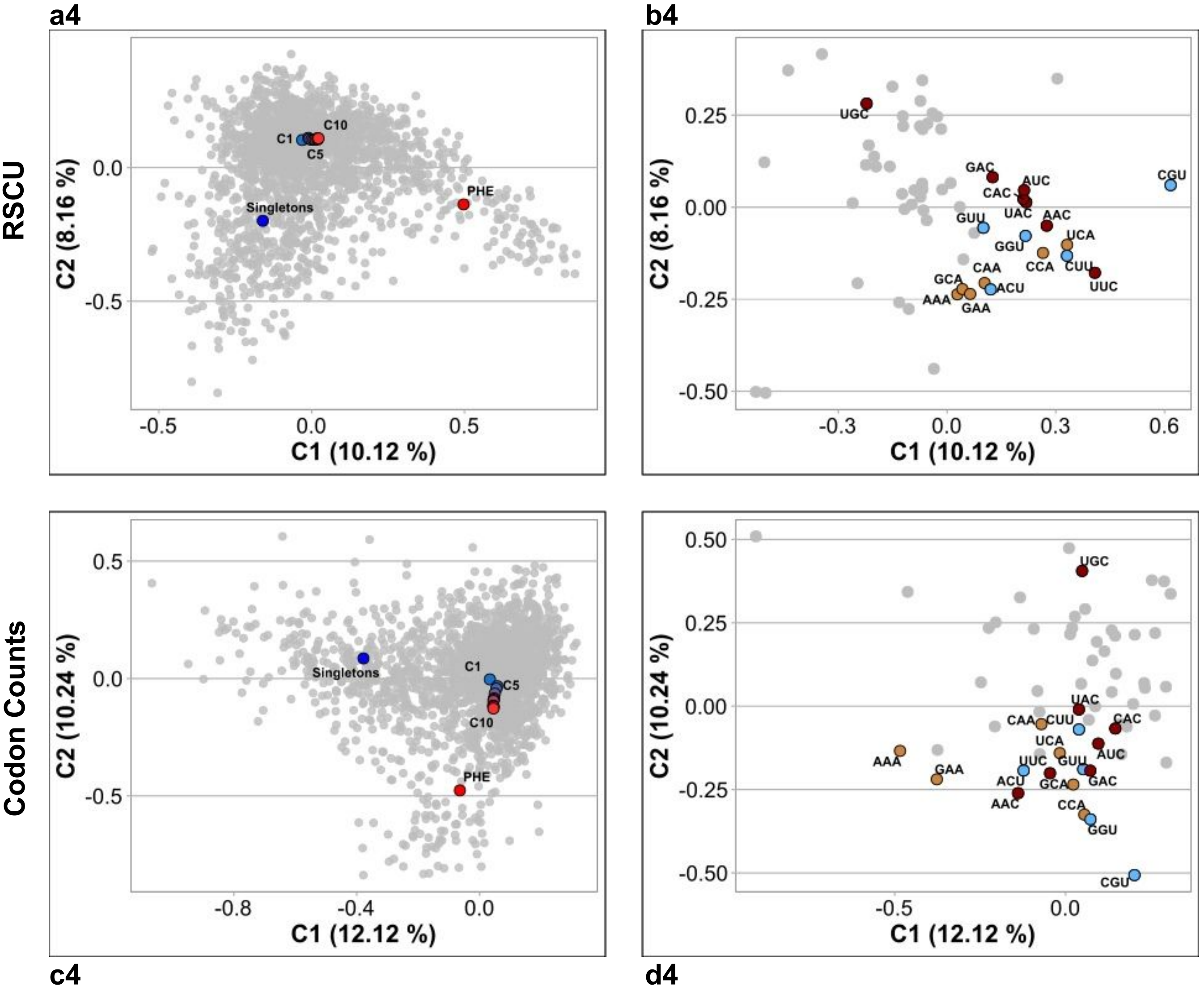

*Bacteroides vulgatus* ATCC 8482 NC_009614

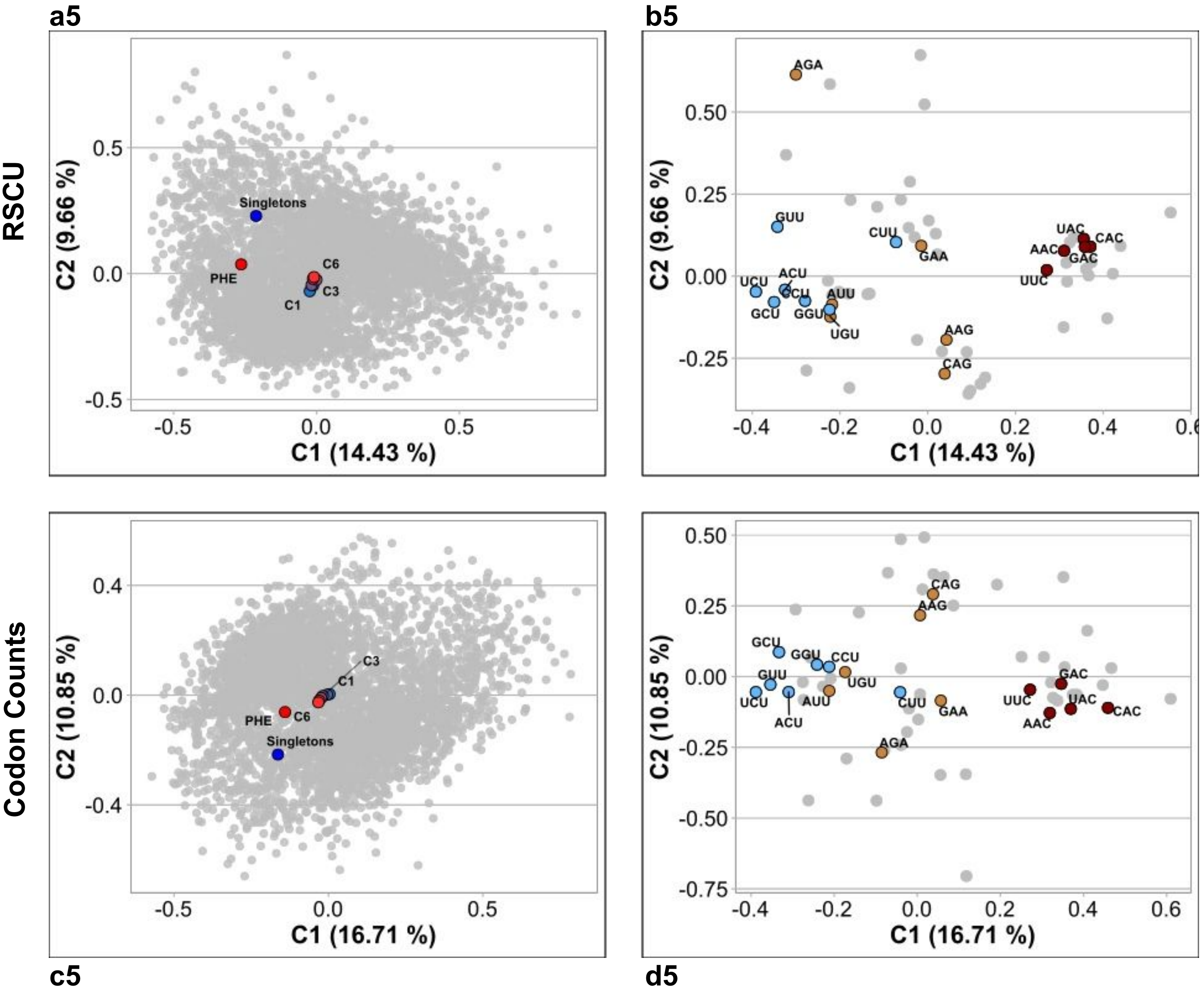

*Bacillus subtilis* subsp spizizenii TU B 10 NC_016047

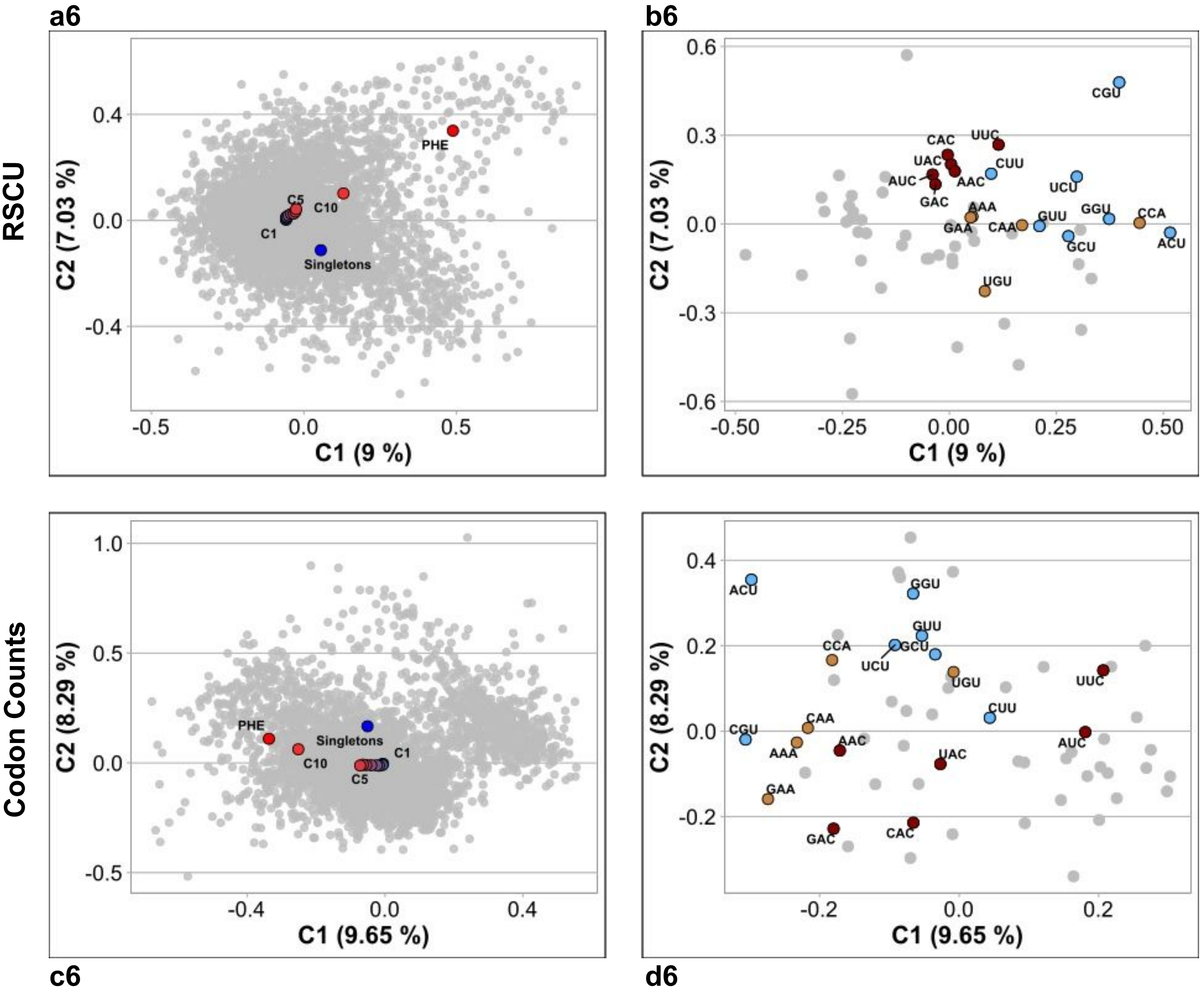

*Moraxella bovis* CCUG 2133 NZ_MUXV01000108

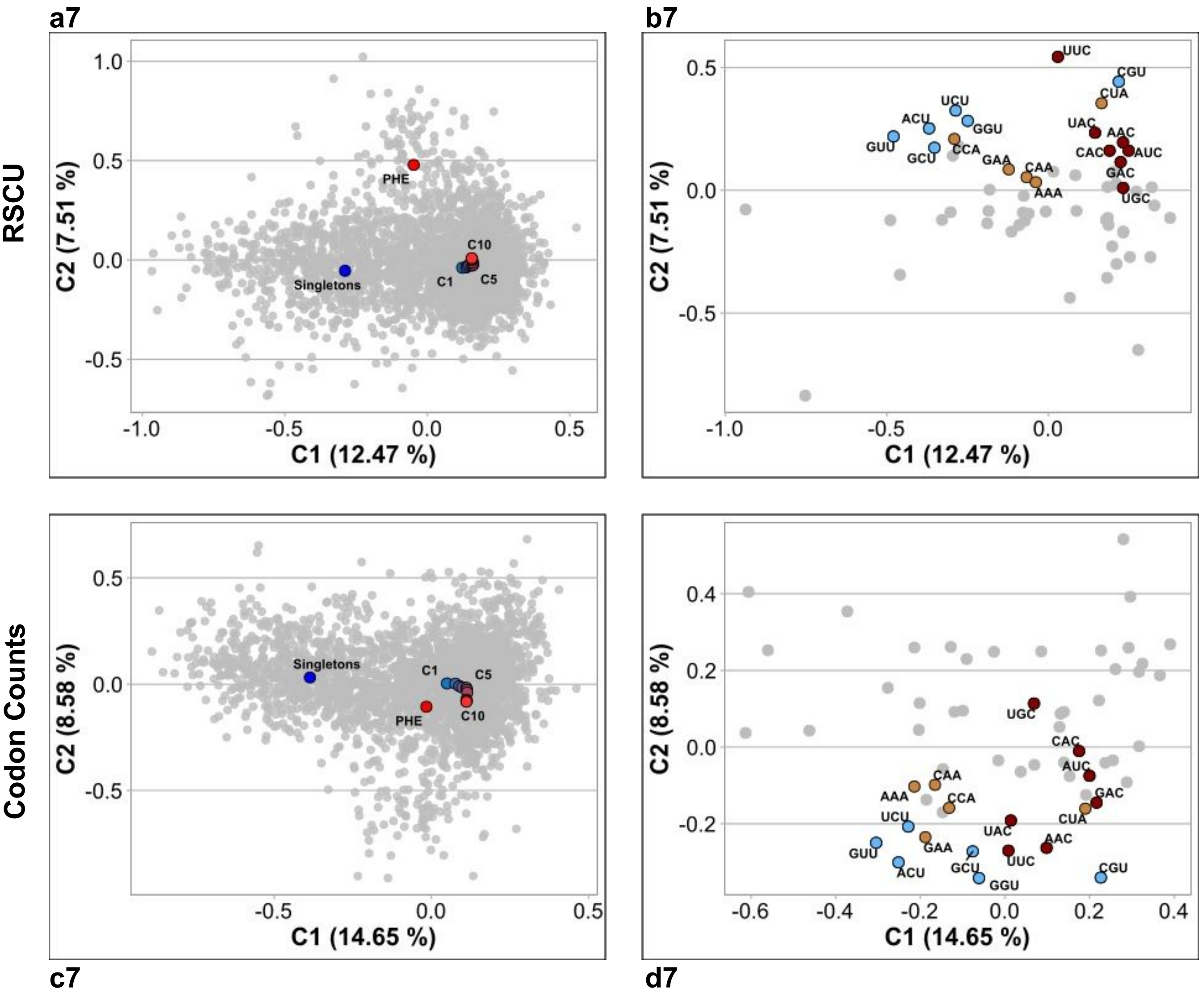

*Chromobacterium violaceum* ATCC12472

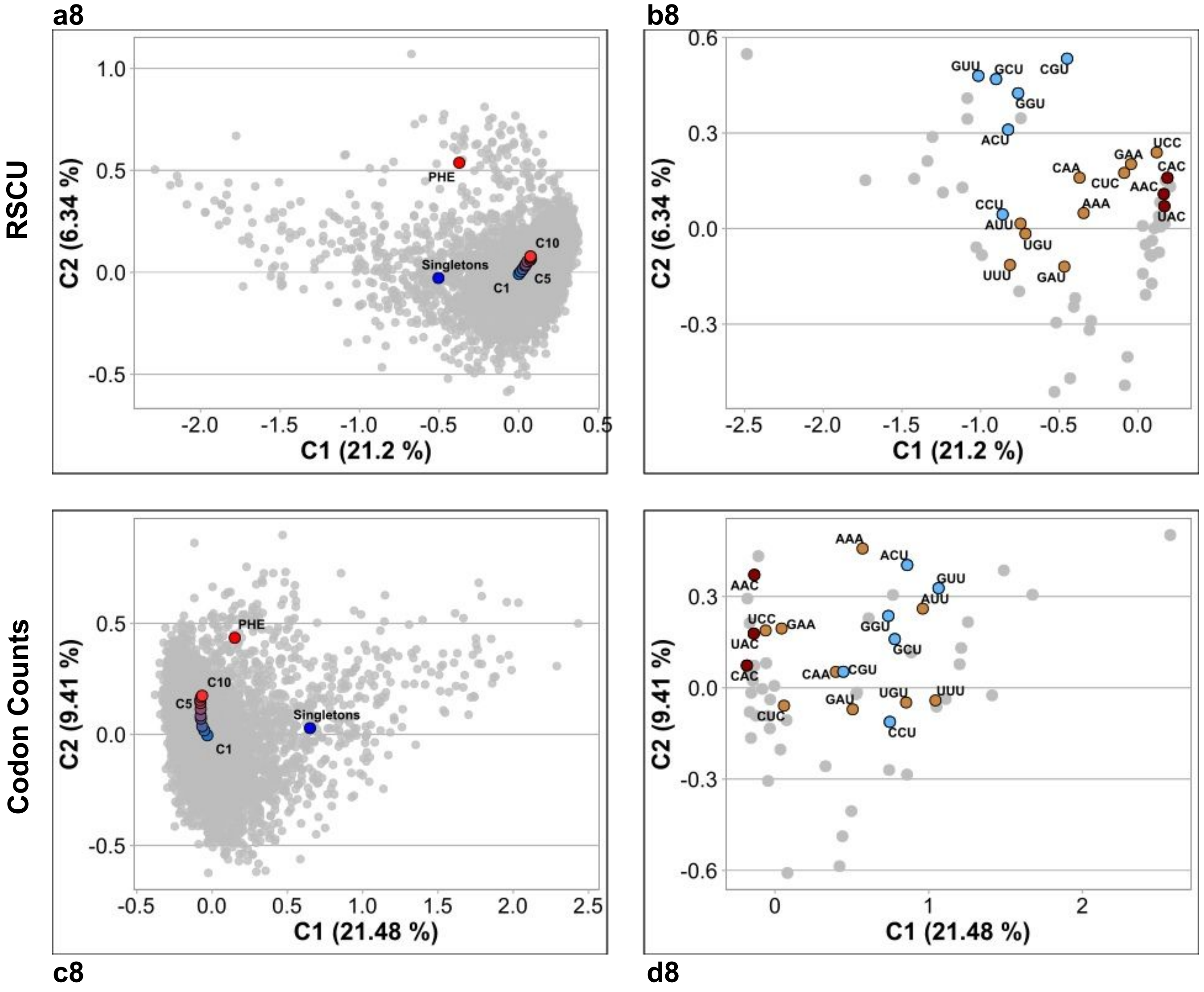

*Paenibacillus graminis* DSM 15220 NZ_CP009287

**a9** **b9**

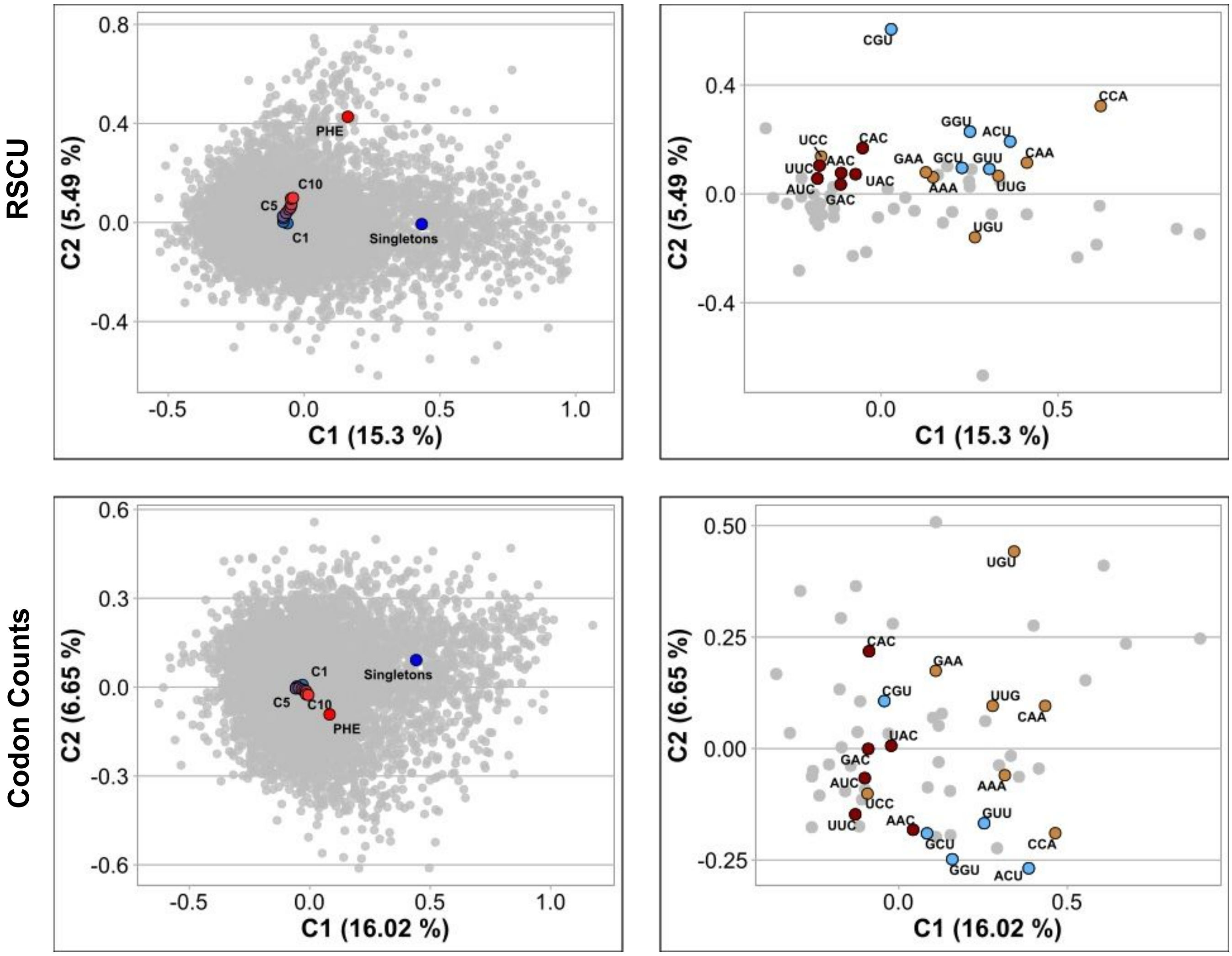


**c9** **d9**

*Photobacterium gaetbulicola* Gung47

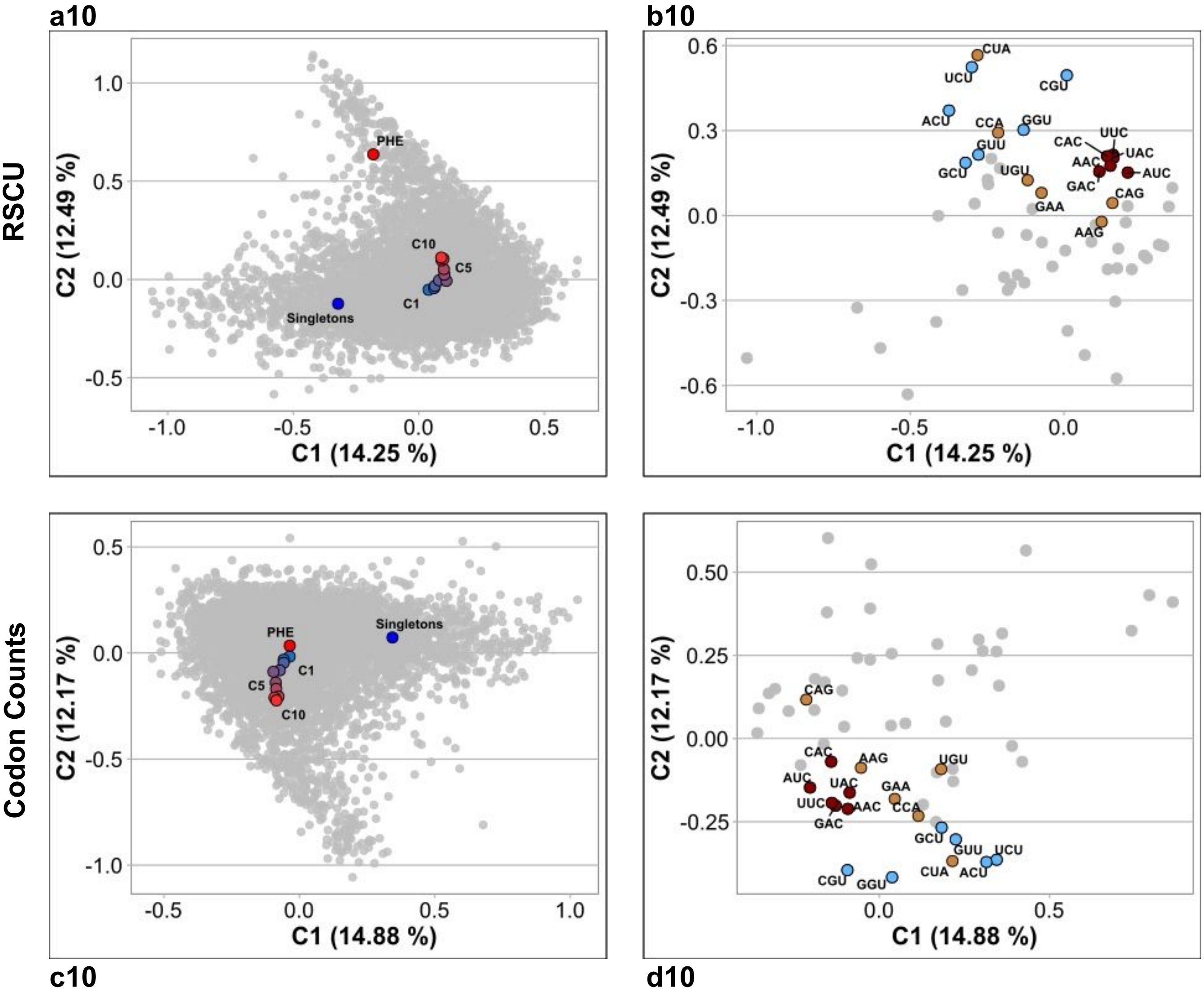

**GROUP C**

*Treponema succinifaciens* DSM 2489 NC_015385

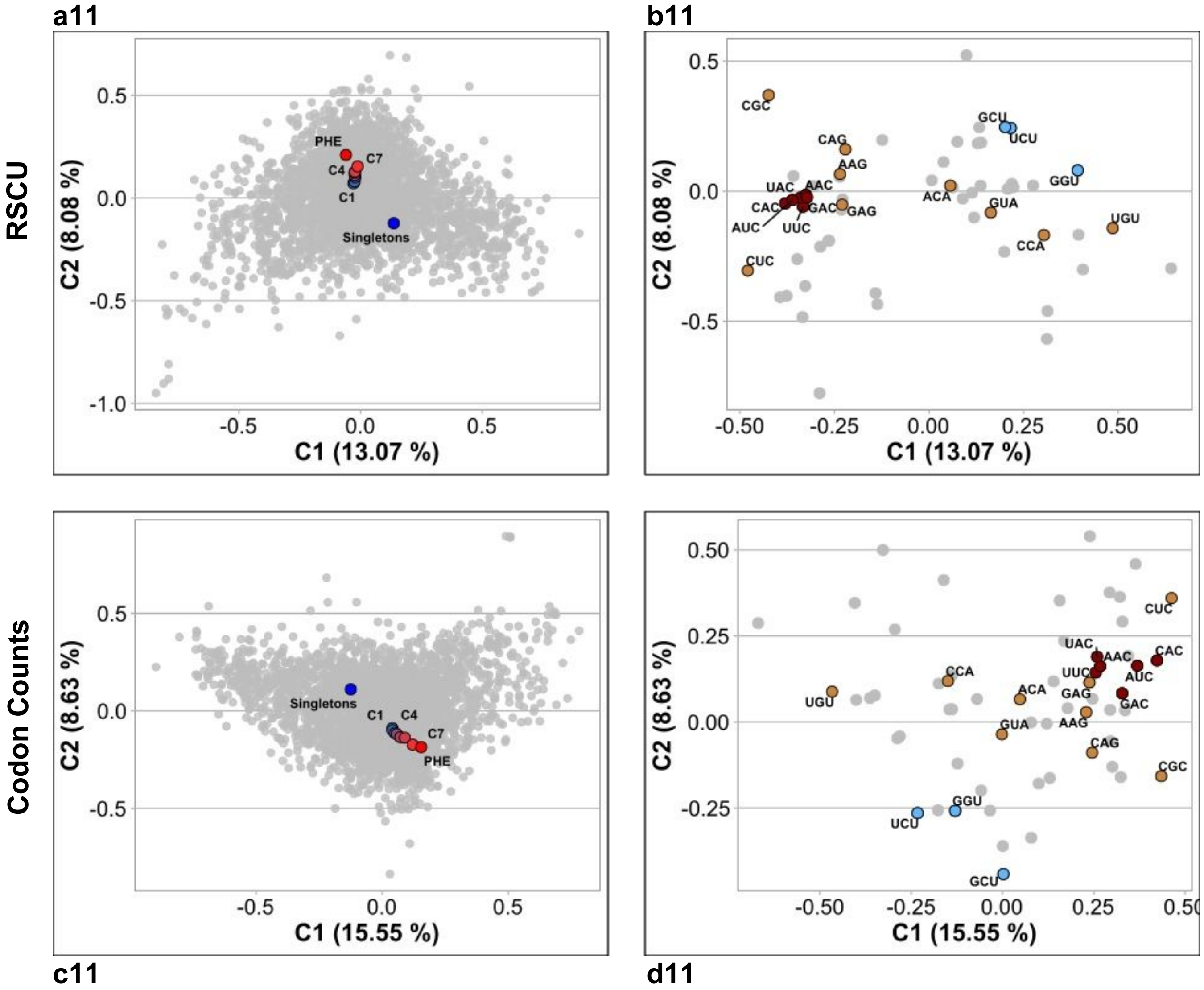

*Prevotella melaninogenica* ATCC 25845 NC_014371

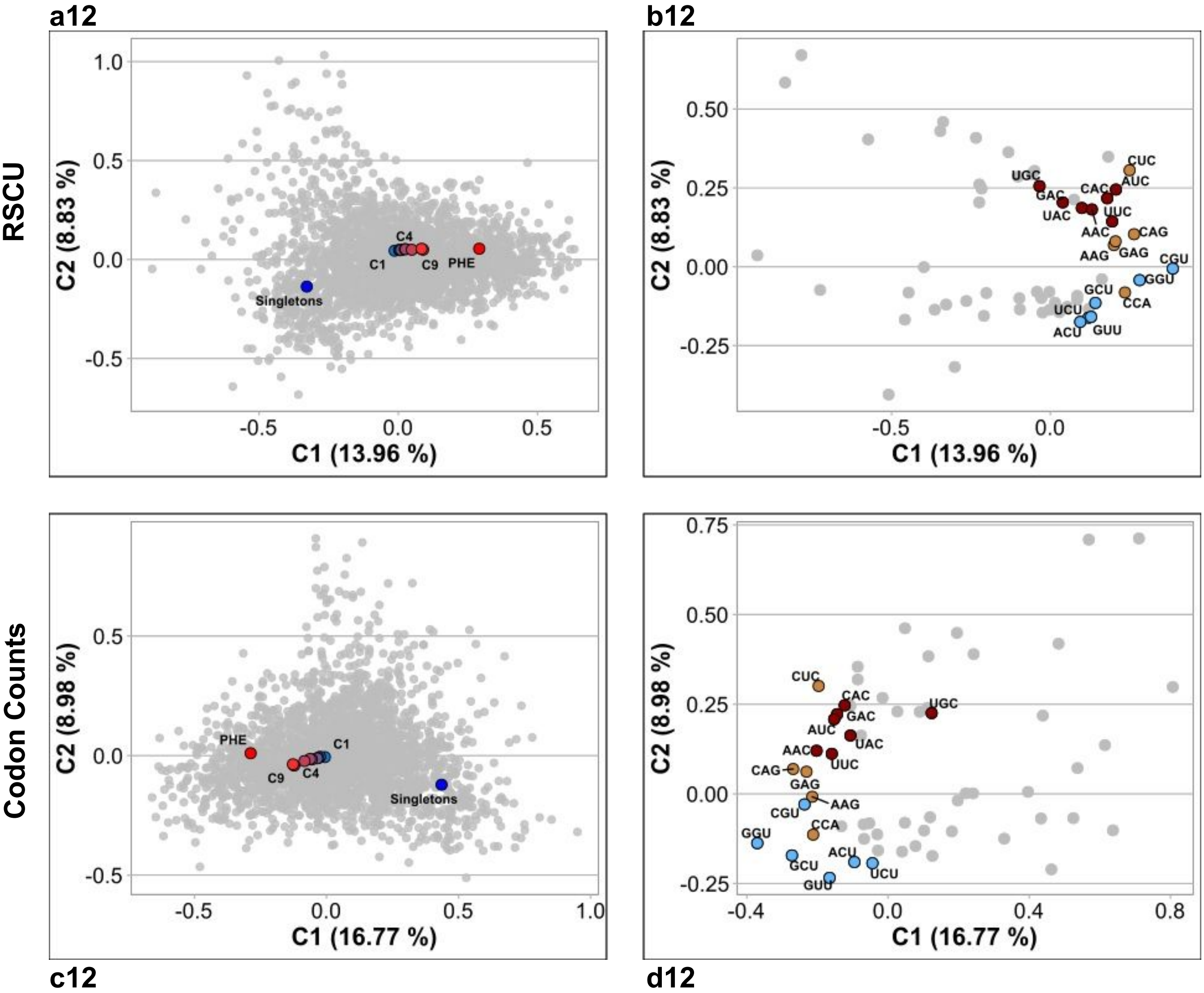

*Yersinia enterocolitica* subsp palearctica Y11 NC_017564

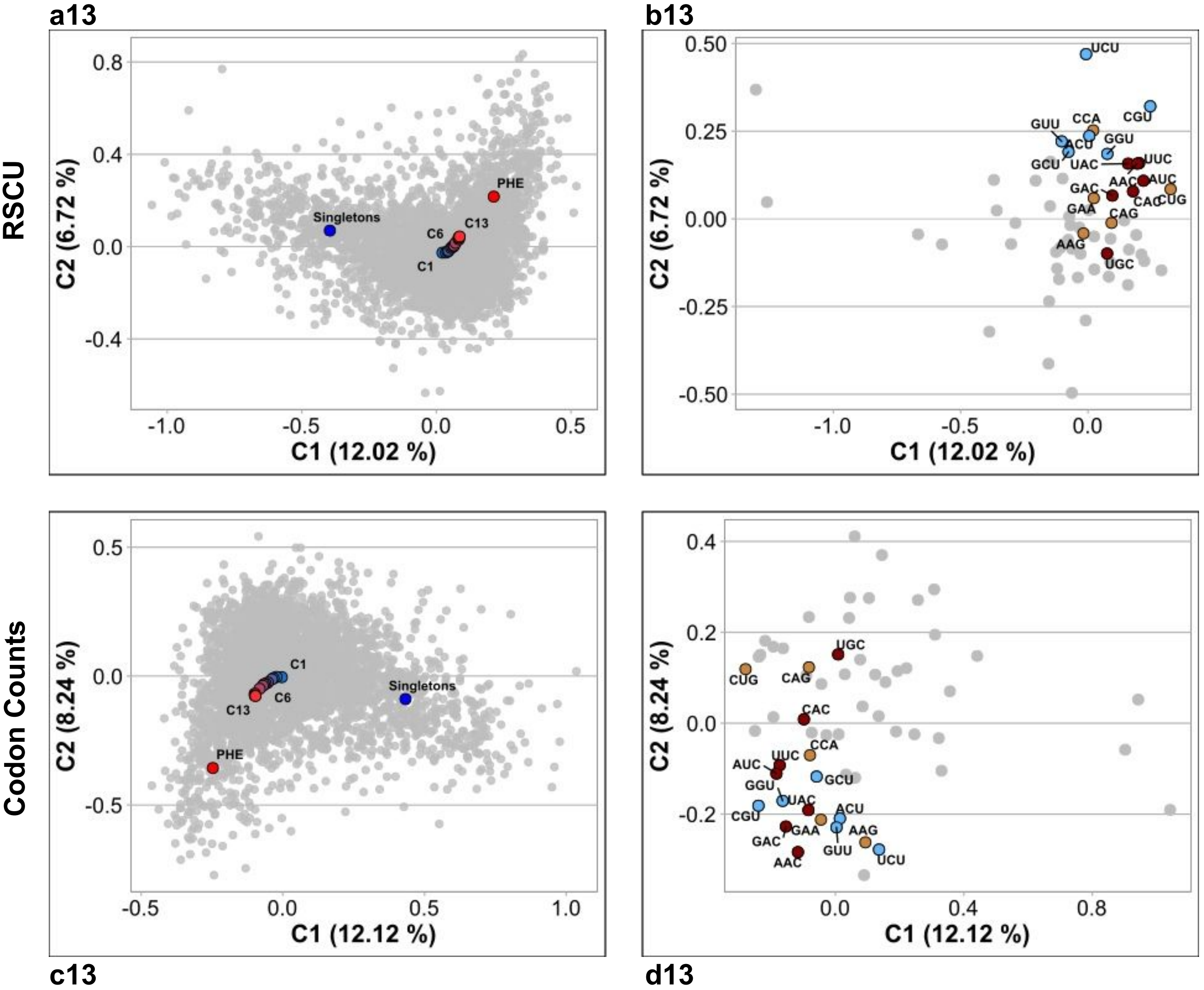

*Methanolacinia petrolearia* DSM 11571 NC_014507

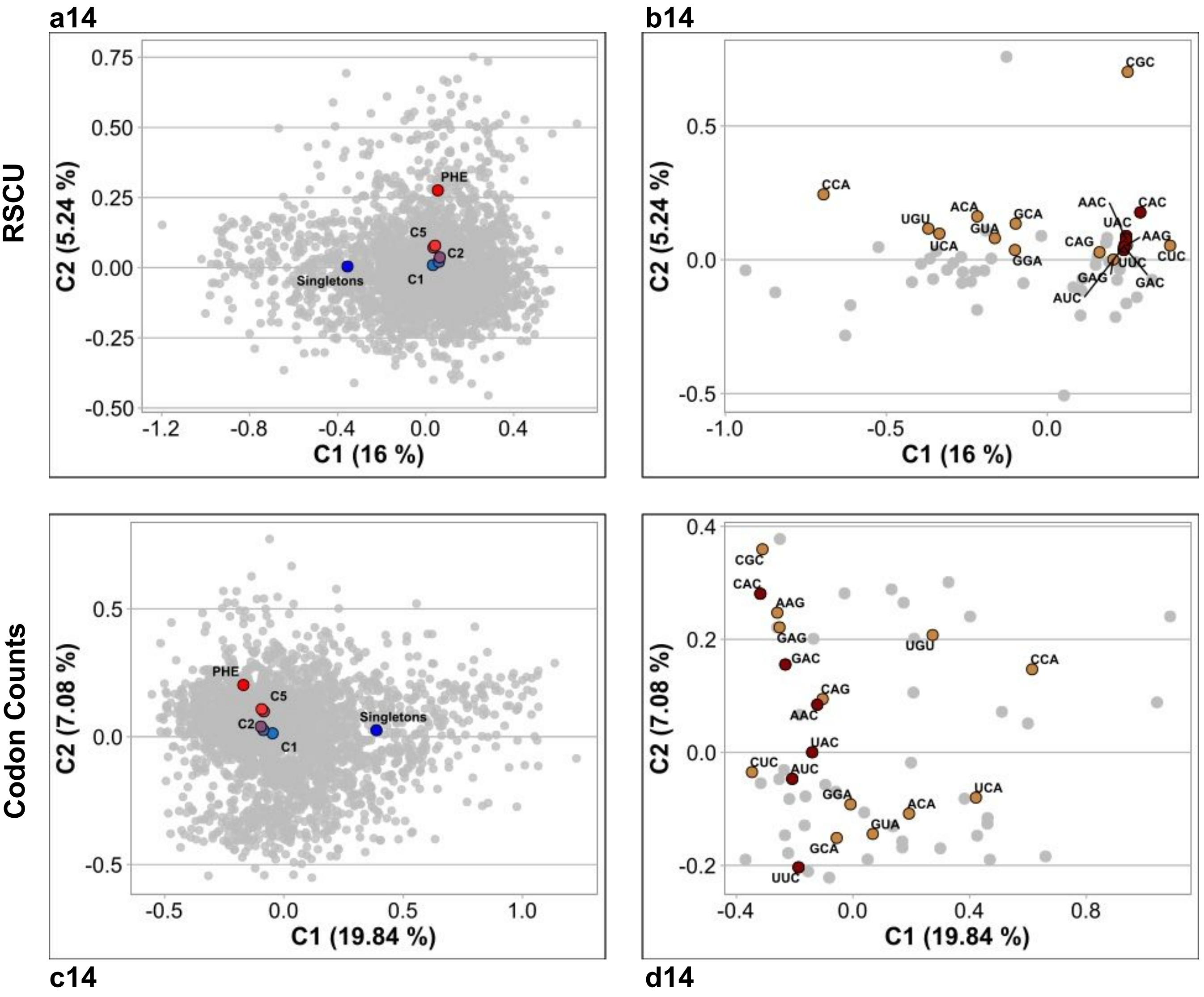

*Bifidobacterium longum* subsp longum JCM 1217 NC_015067

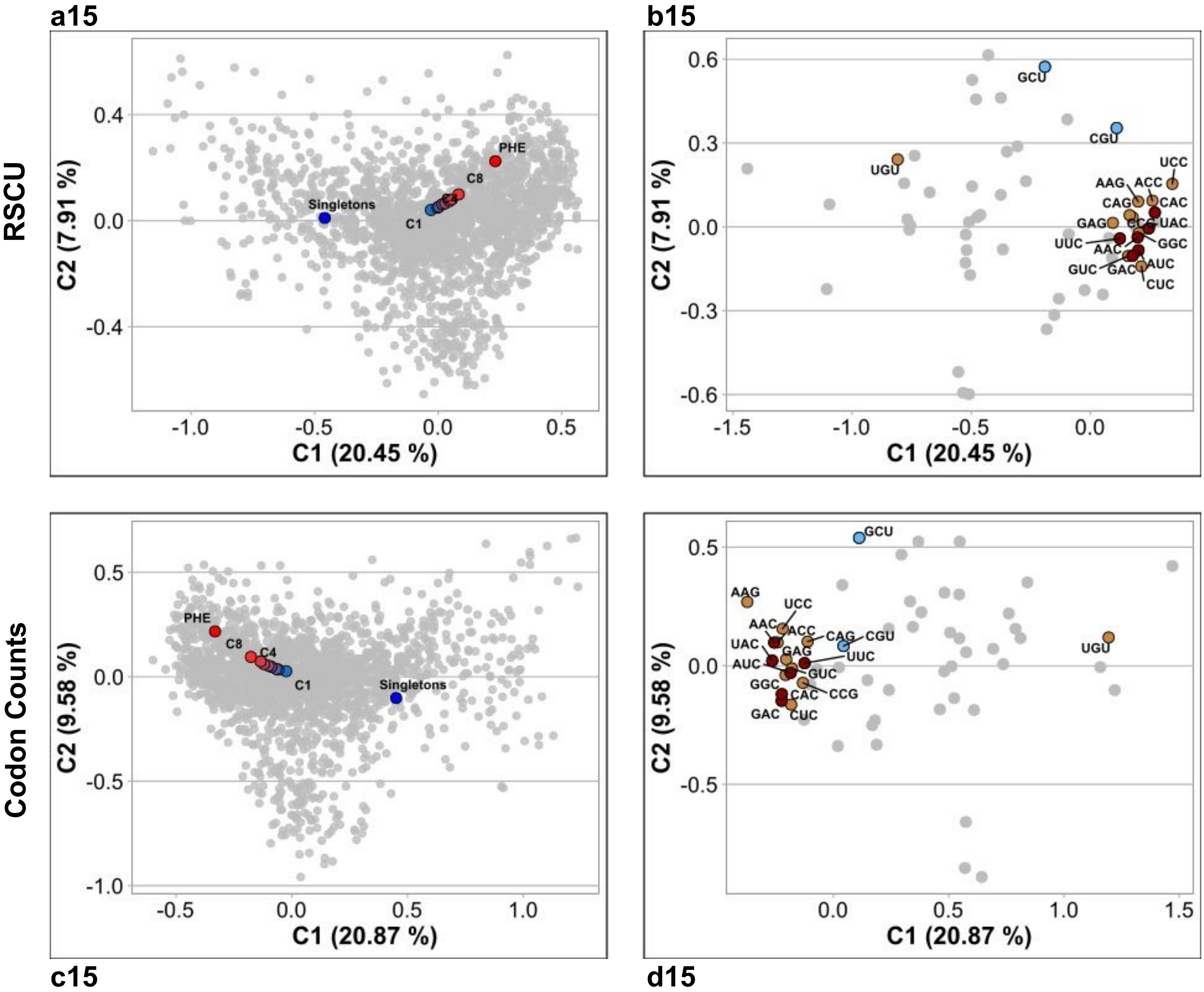

*Bordetella holmesii* ATCC 51541 NZ_CP007494

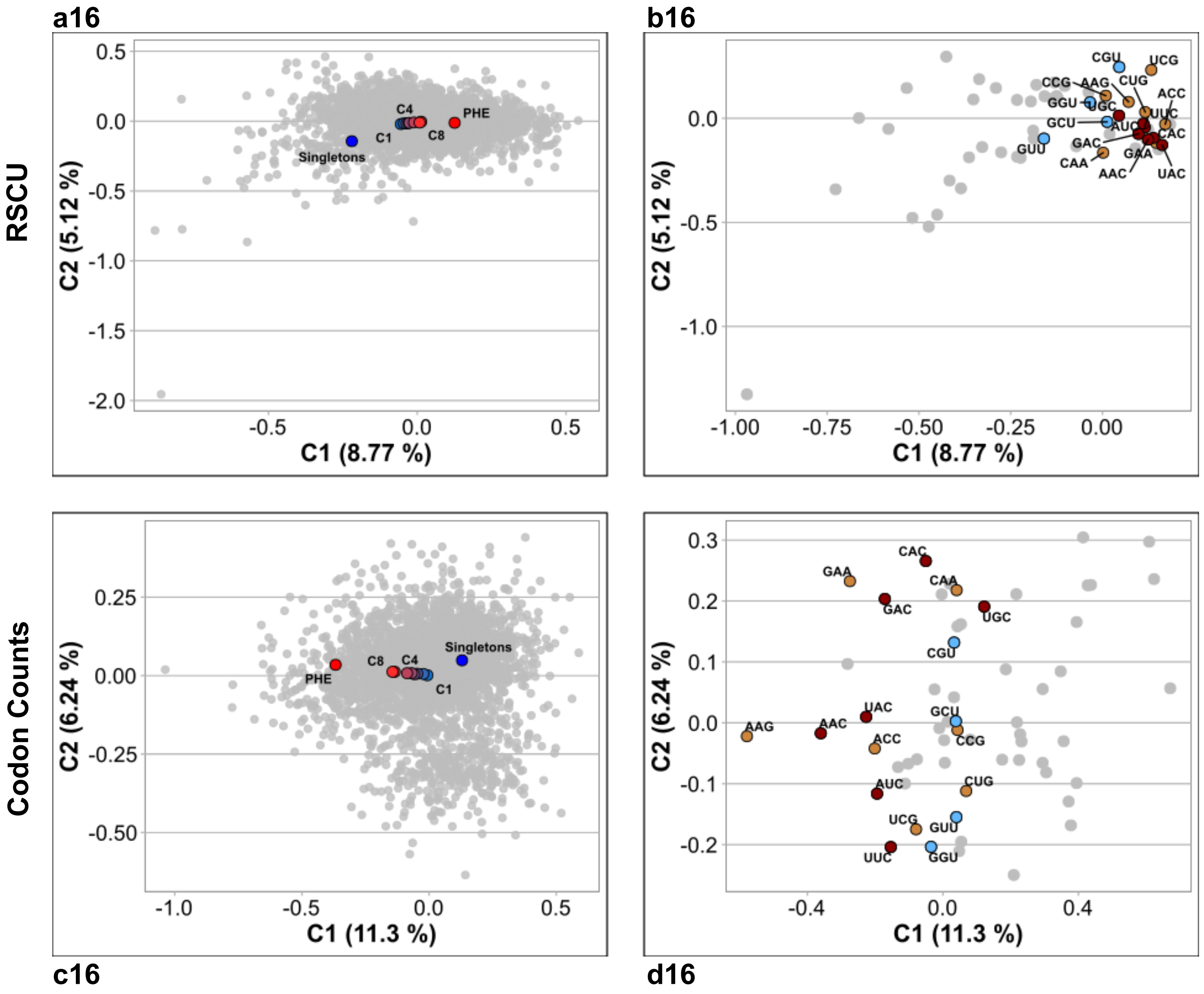

*Mycobacterium fortuitum* subsp fortuitum DSM 46621  ATCC 6841 CP014258

**a17** **b17**

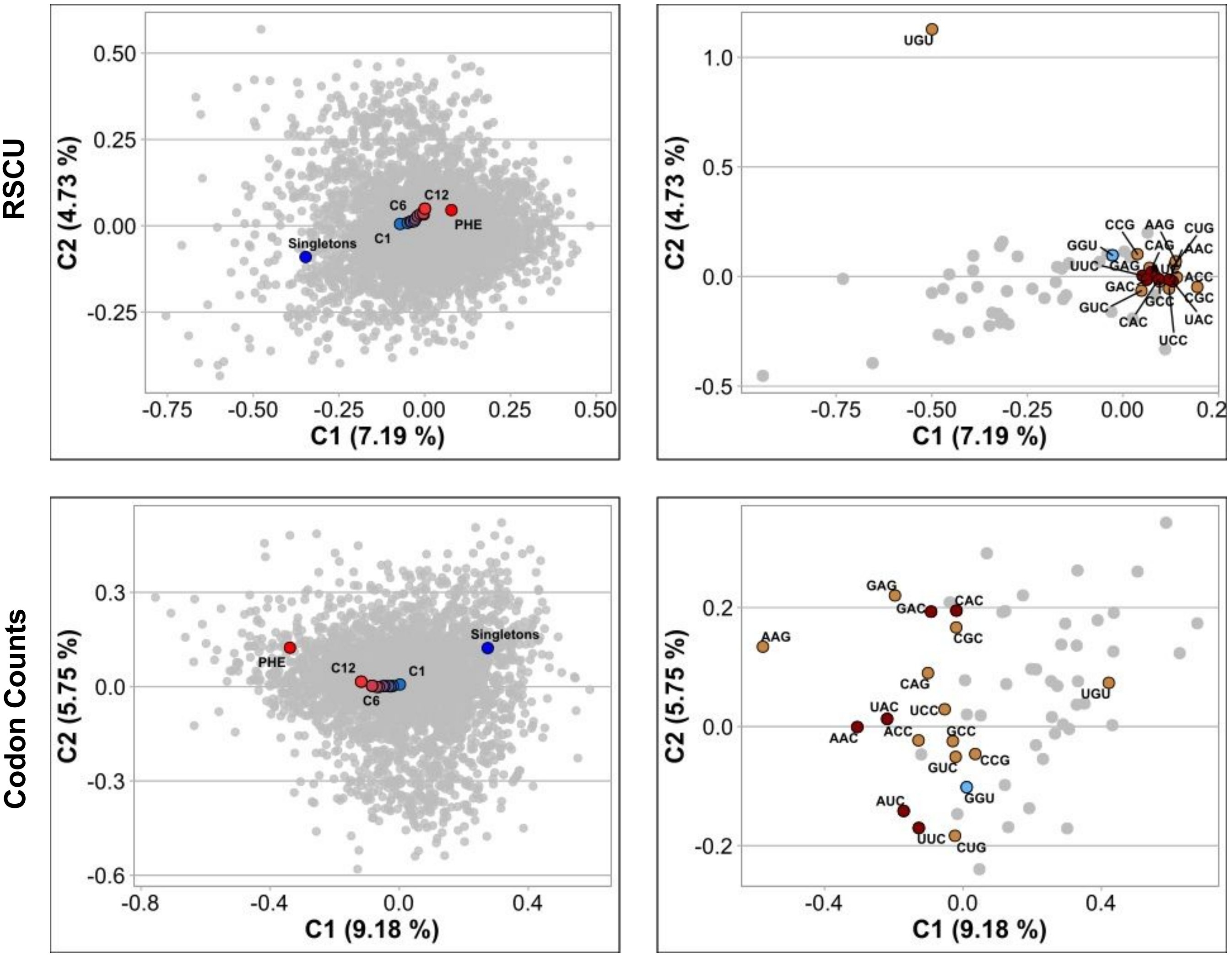


**c17** **d17**

*Sphingomonas parapaucimobilis* NBRC 15100 NZ_BBPI01000001

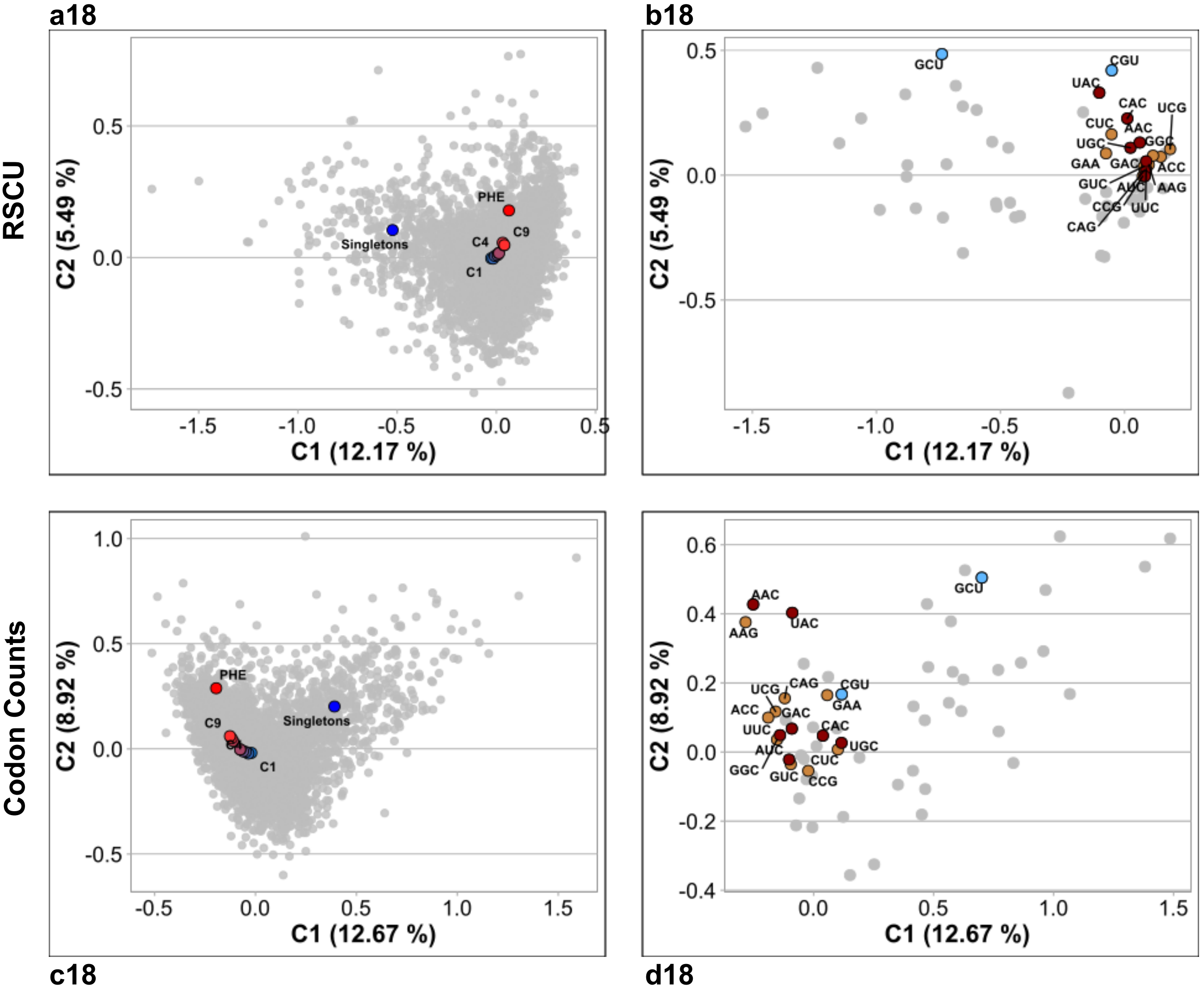

*Atopobium parvulum* DSM20469

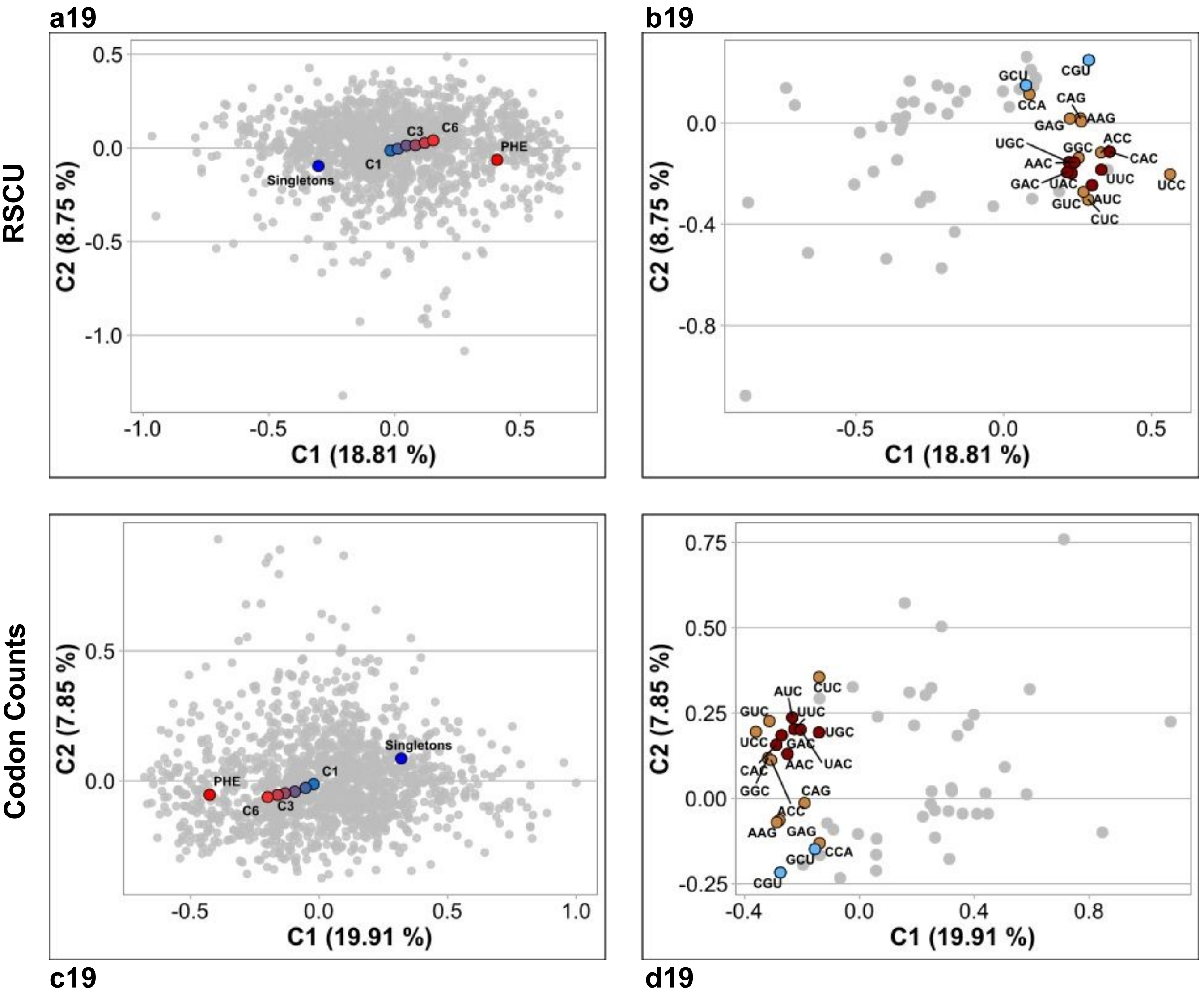

***GROUP D***

*Leisingera methylohalidivorans* DSM 14336

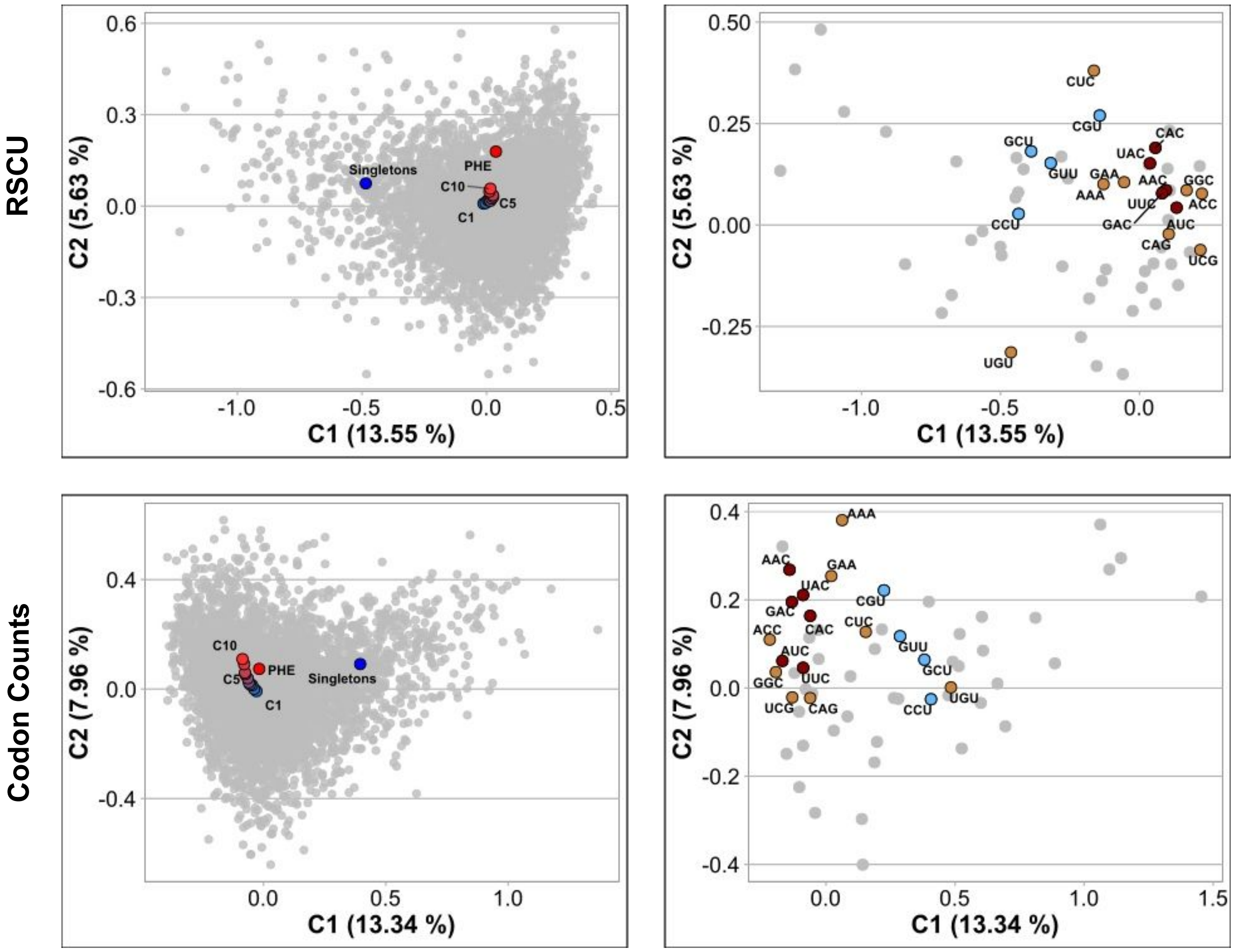


c20 d20

*Oceanibaculum pacificum* MCCC 1A02656 NZ_LPXN01000181

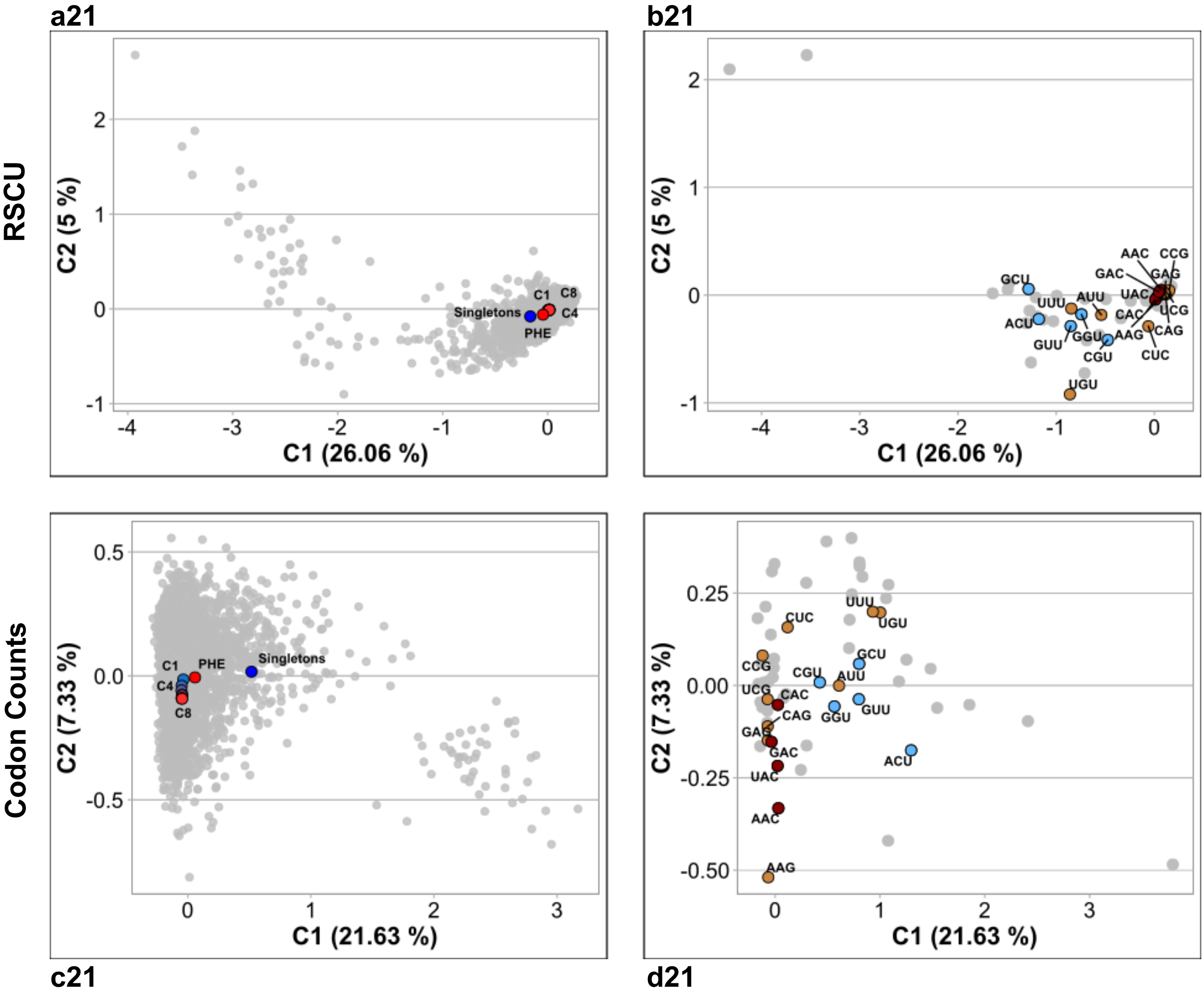

*Arthrobacter enclensis* NIO 1008 NZ_KQ758616

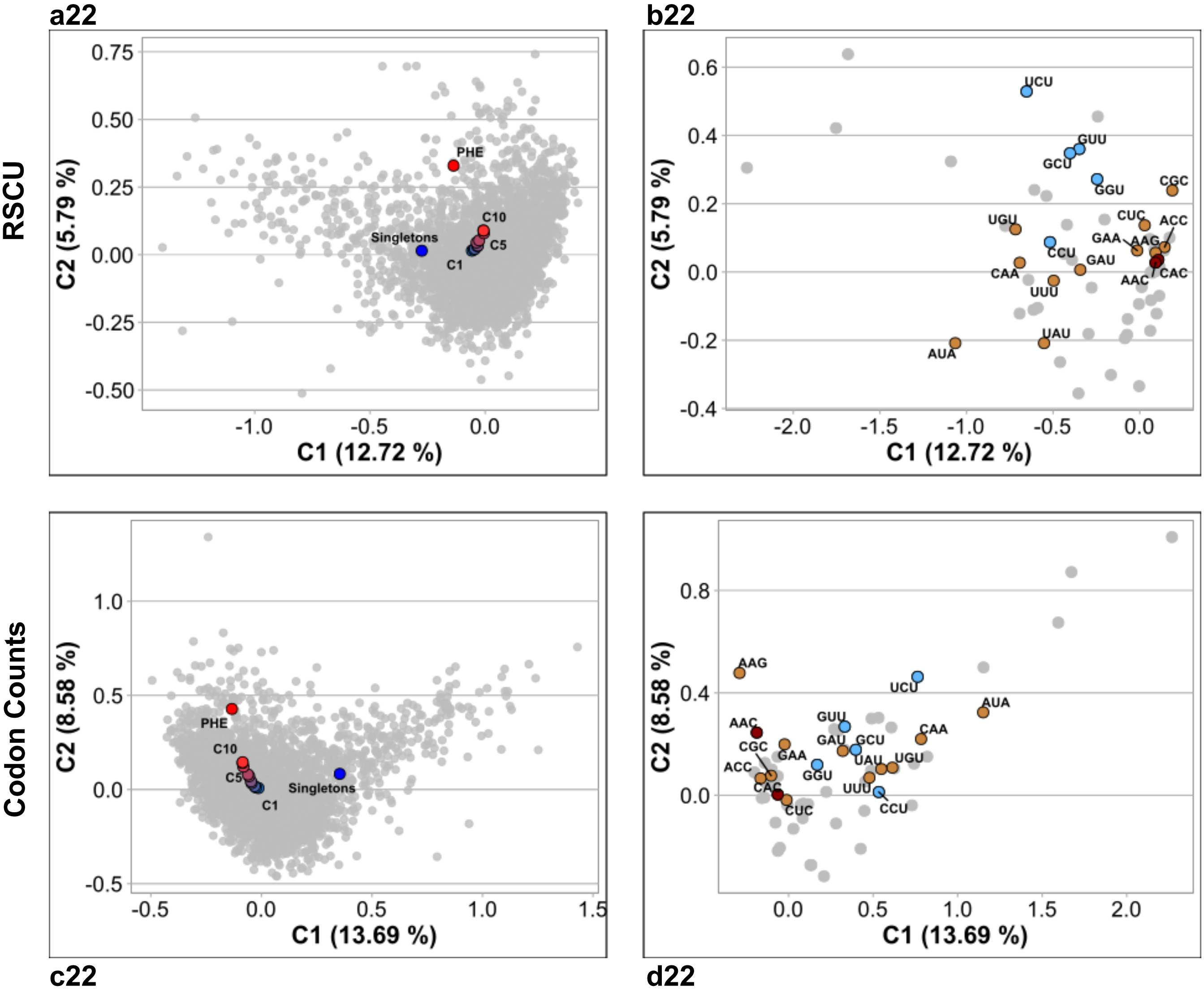

*Rhodanobacter denitrificans* 2APBS1

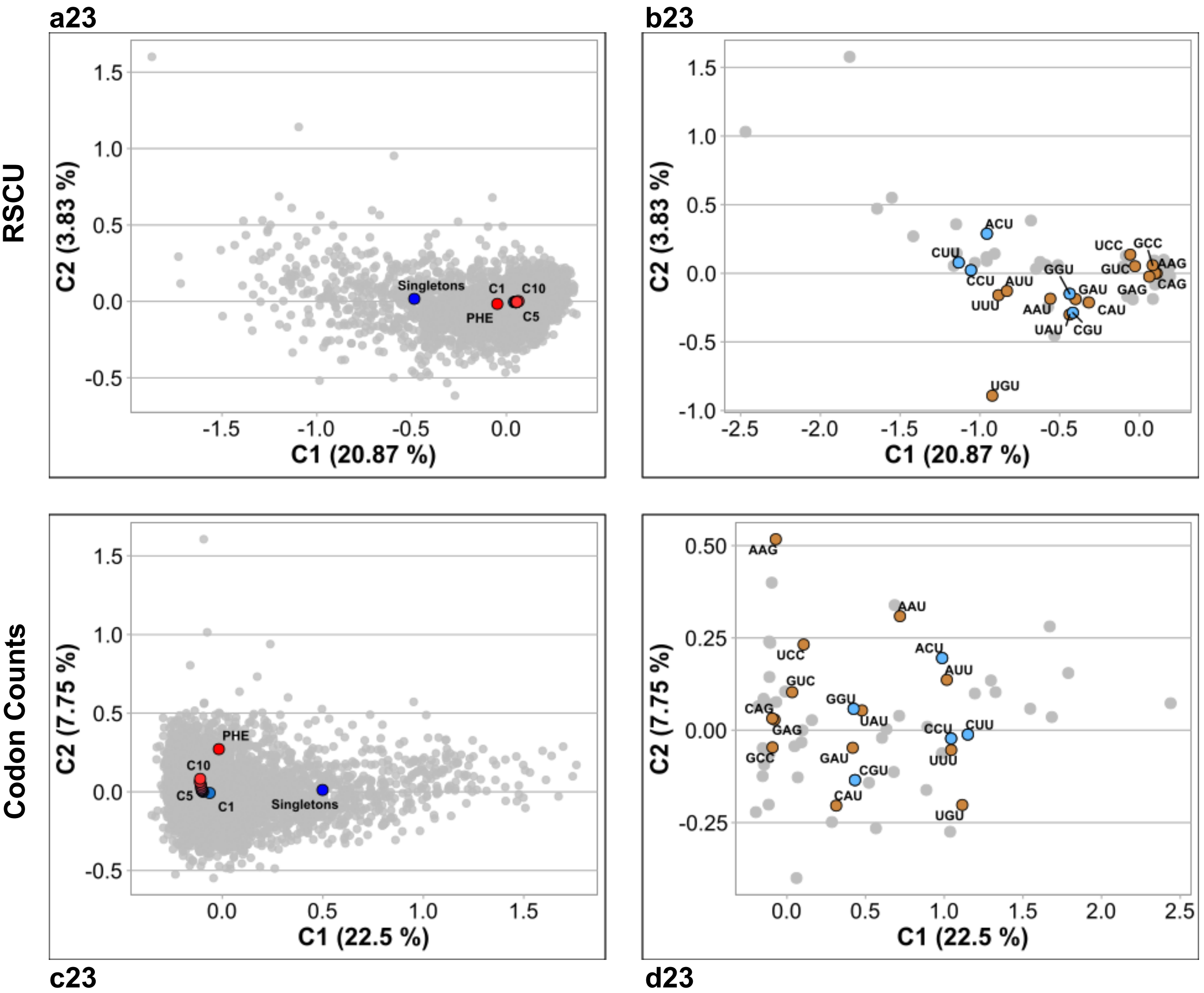

*Microbacterium aurum* KACC 15219 NZ_CP018762

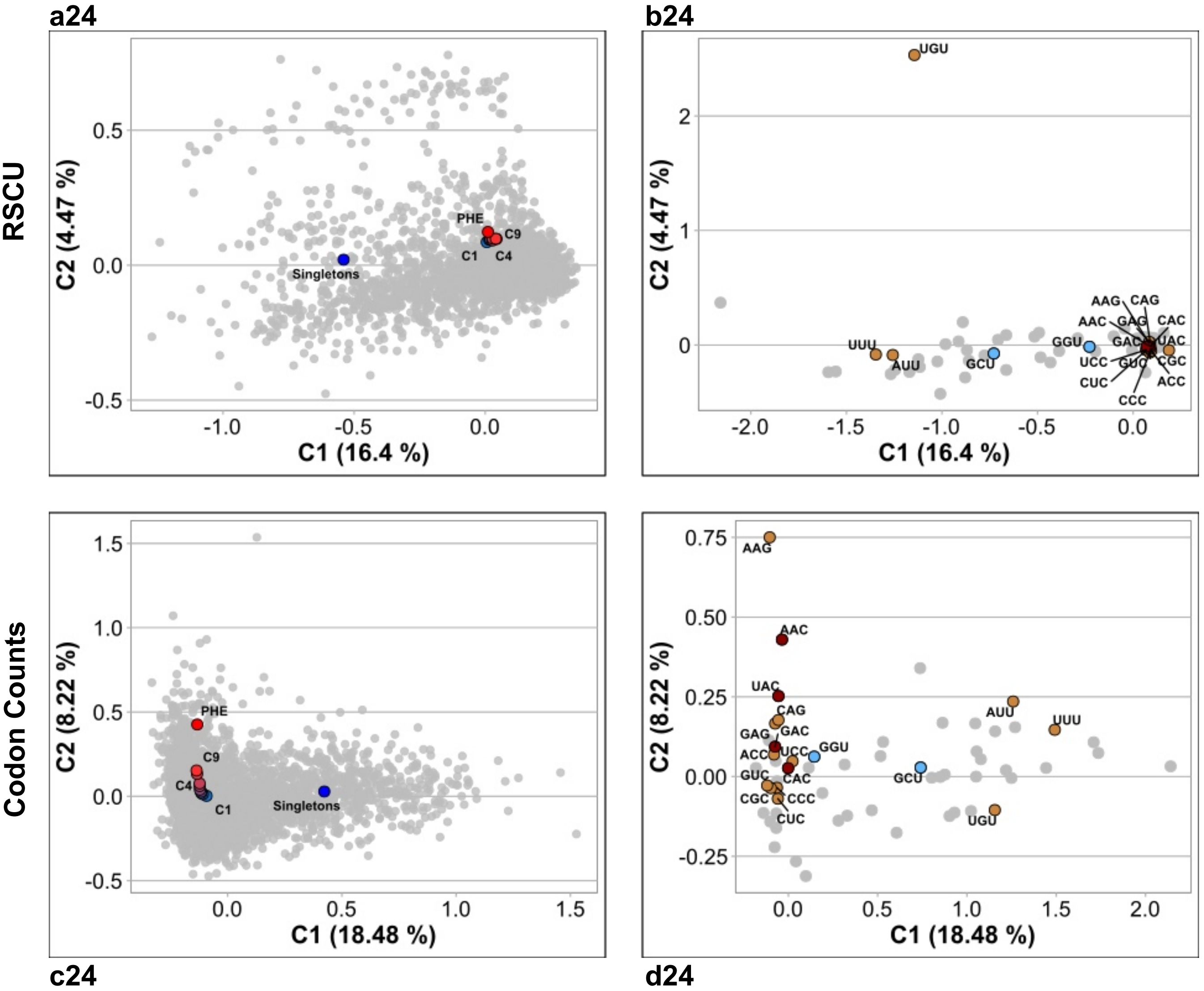

*Roseomonas mucosa* ATCC BAA 692 NZ_JHWD01000081

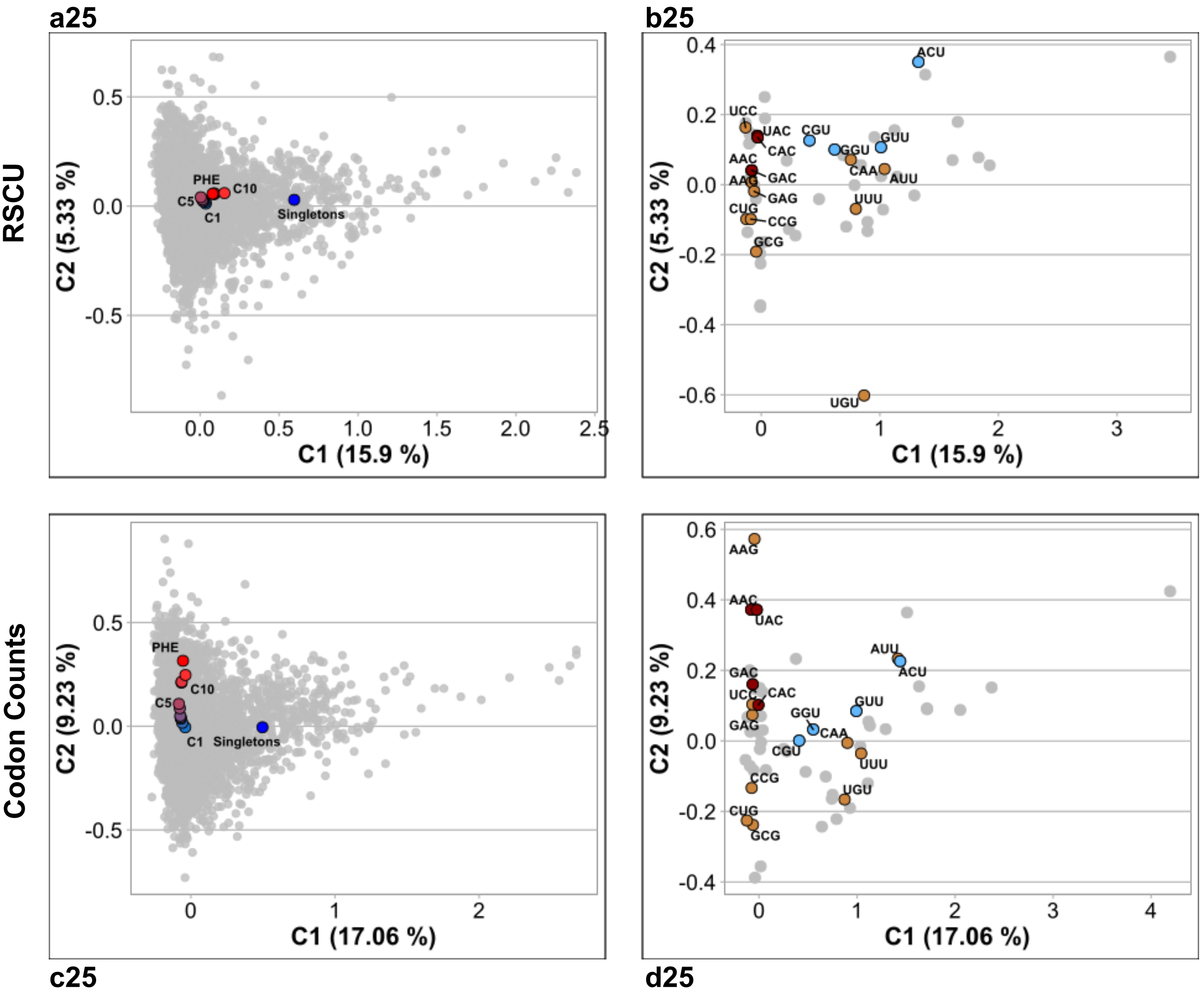

**Fig. S1B. Correspondence-Analysis (CA) plots of core-gene sets with different degrees of conservation throughout the phylogeny of 25 prokaryote families.** The strains that were not included in the main manuscript are represented here and classified as belonging to Group A, B, C, or D. **Panel a1 to a25:** The individual genes (in gray) are represented in the space of the first-two CA components, with the percent variation of components 1 and 2 being indicated on the axes. CAs were computed using either raw codon counts (RCC) or RSCU as the input variables as indicated. Average coordinates (centroids) for different gene sets (i.e. singletons in blue, C1 to Cn in a gradient from blue to red, and PHE in red) were projected on the CA space in the RCC-based CA plots. Similarly, modal codon usages for the same gene sets—calculated as indicated in Materials and Methods—were projected on the RSCU-based CA plots. In C1 to Cn the higher number denote a more ancestral core-gene set within the phylogeny. Table S1_a (tab 1) lists the prokaryote species that were used to construct each Ci gene set by means of the EDGAR software (54, 55). **Panel b1 to b25.** Plots describing codon relative weight in the first two principal-component positions of the CA. Codons that are colored in brown indicate those which present—for each amino acid—the highest CUF enrichment from C1 to PHE (i.e. those codons that better represent translational adaptation) except when some of those codons also correspond to a C-bias (colored in red) or a U-bias (colored in light blue).
