## Supplementary material for "Codon-usage optimization in the prokaryotic tree of life: How synonymous codons are differentially selected in sequence domains with different expression levels and degrees of conservation": Suplemental Figure 2

### FIGURE S2

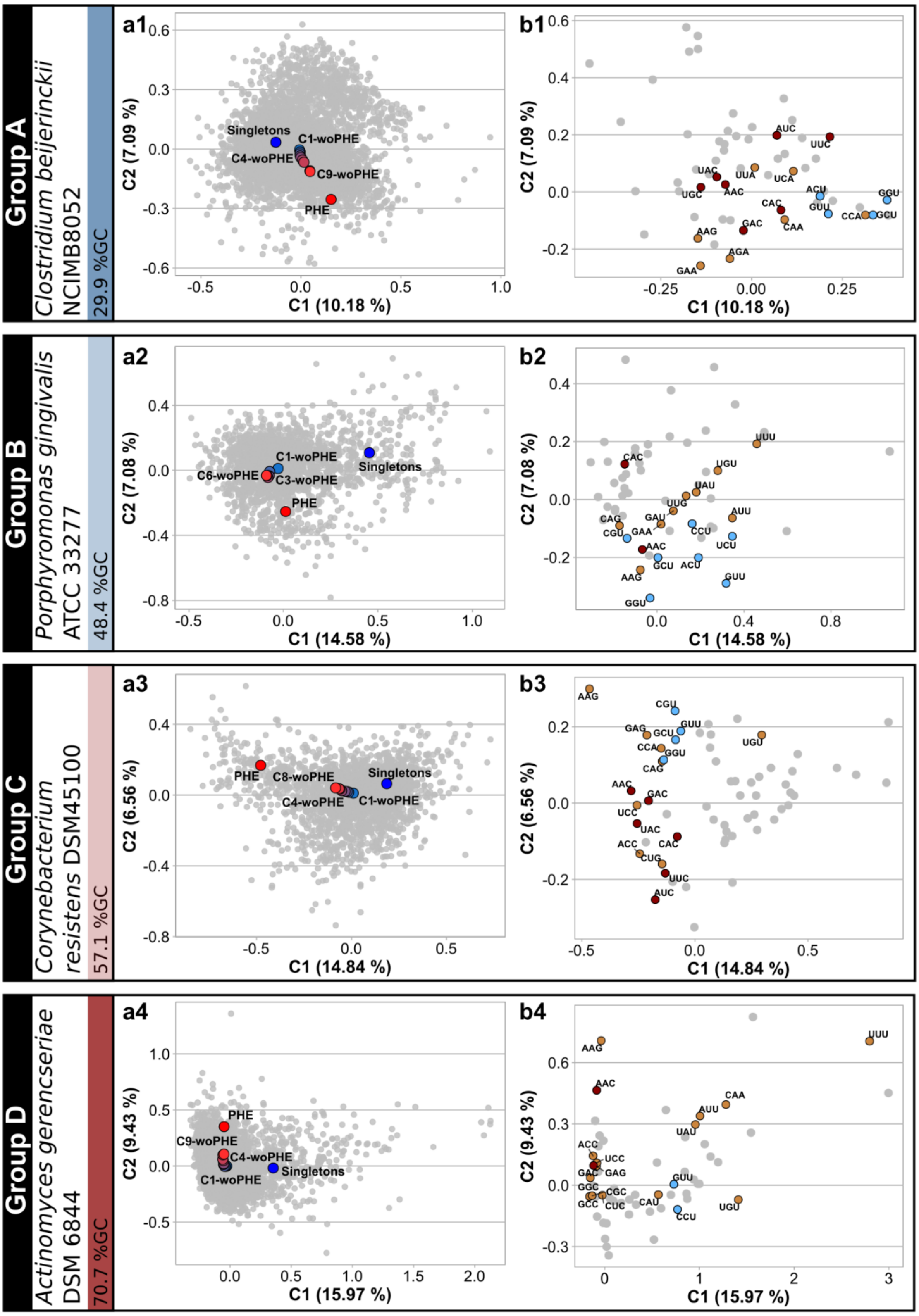


**Fig. S2. RCC-based CA plots of core-gene sets excluding the PHE genes.** PHE genes were extracted from each group of core genes and the resulting gene sets, indicated as Ci-woPHE (Ci-without PHE)(from “1” to “n”, in blue to red circles), were projected on the CA plots as used for the four reference species in Fig. 1. Singletons and PHE are in blue and red, respectively.
