## Supplementary material for "Codon-usage optimization in the prokaryotic tree of life: How synonymous codons are differentially selected in sequence domains with different expression levels and degrees of conservation": Suplemental Figure 3

### Fig. S3

#### Group A

*Methanobrevibacter smithii* ATCC 35061

**a1**

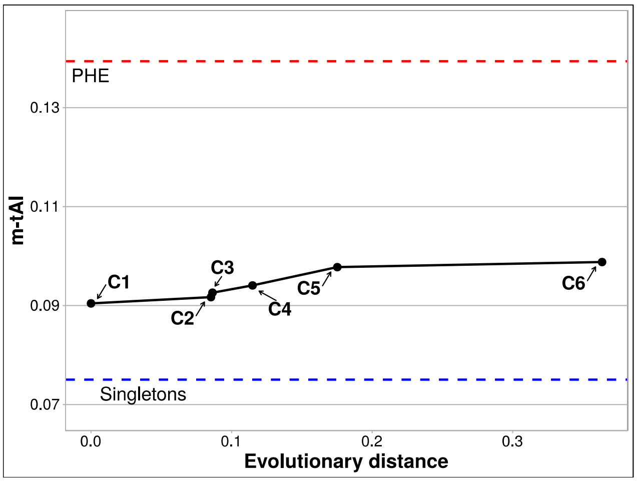


**b1**

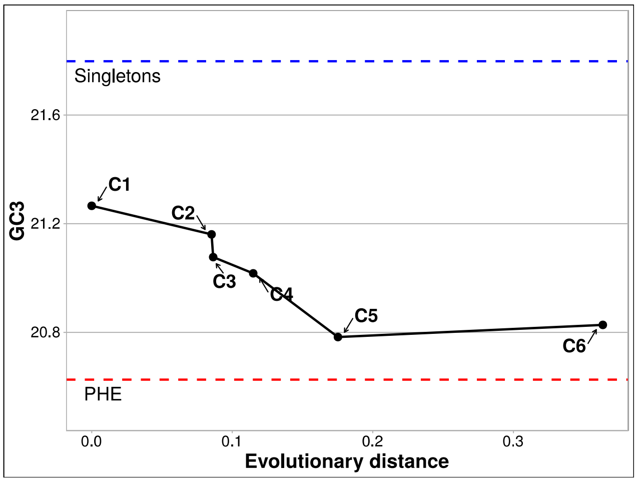


*Tetragenococcus halophilus* NBRC 12172 NC_016052

**a2**

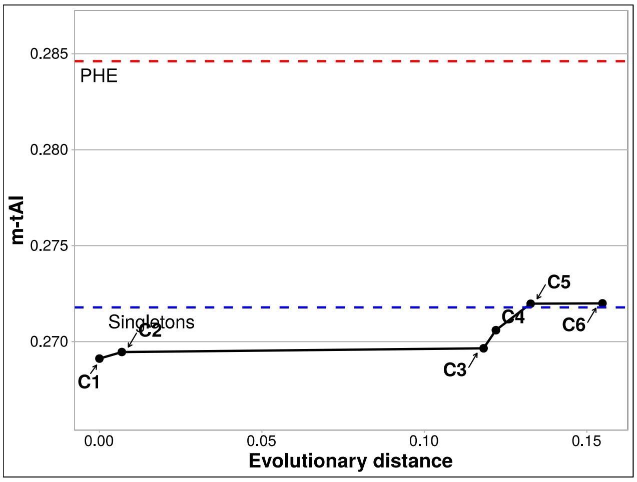


**b2**

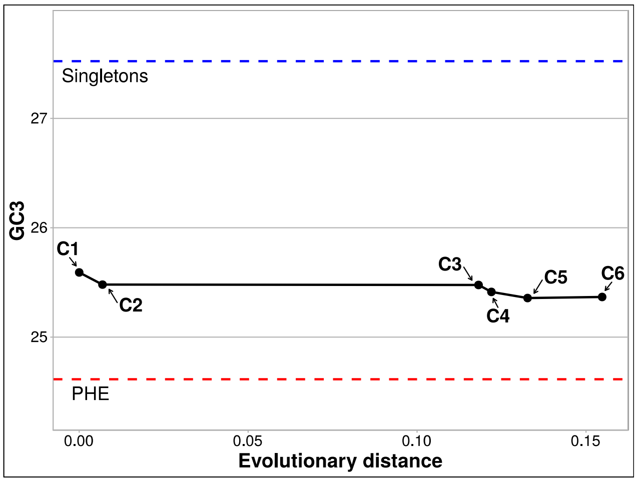


#### Group B

*Sulfurospirillum multivorans* DSM 12446 NZ_CP007201

**a3**

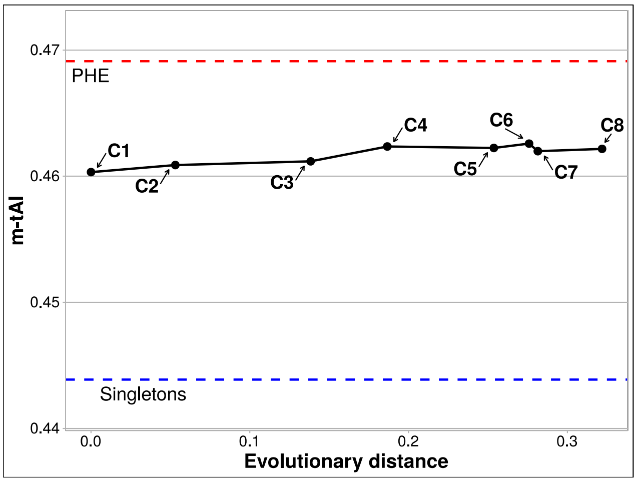


**b3**

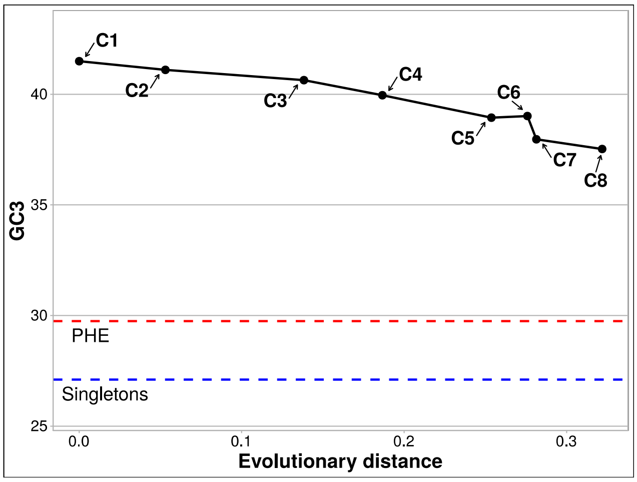

*Streptococcus equi* ATCC 33398 NZ_FTNH01000046

**a4**

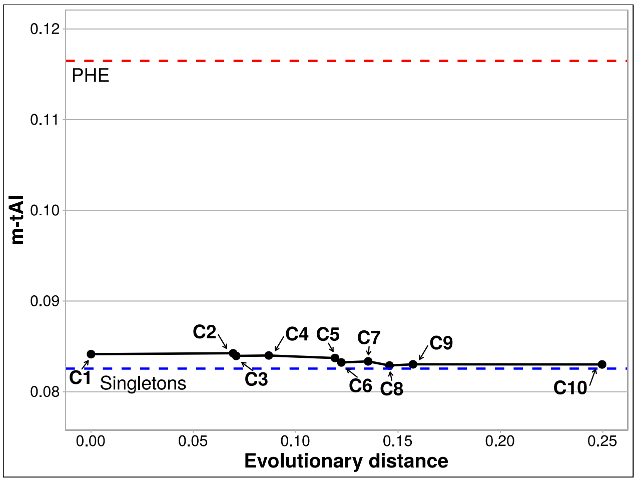


**b4**

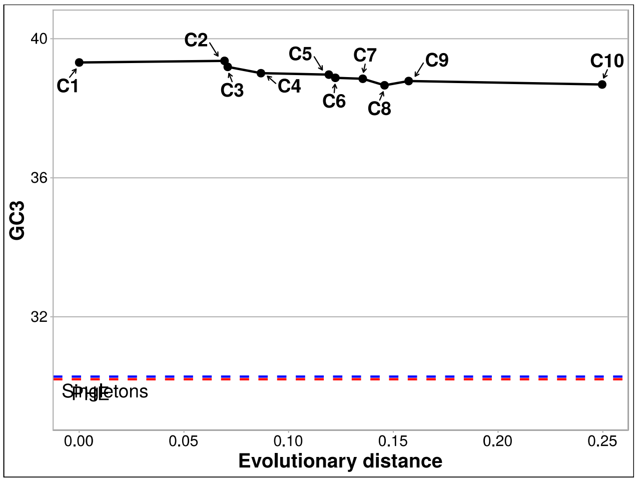


*Bacteroides vulgatus* ATCC 8482 NC_009614

**a5**

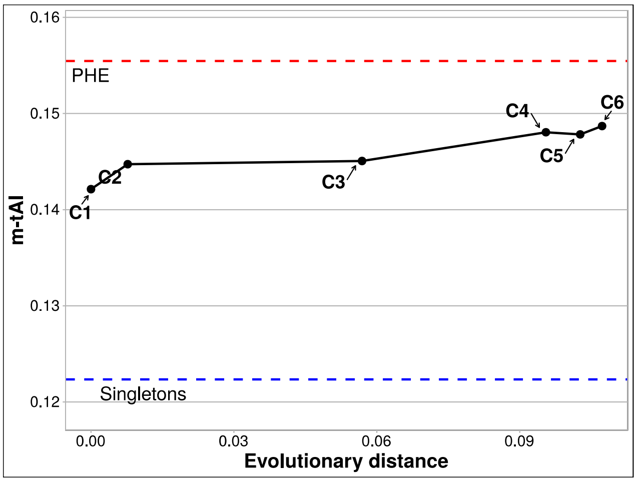


**b5**

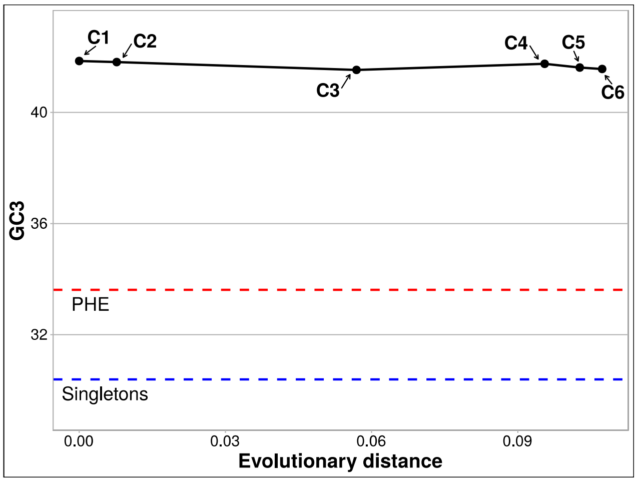


*Bacillus subtilis* subsp spizizenii TU B 10 NC_016047

**a6**

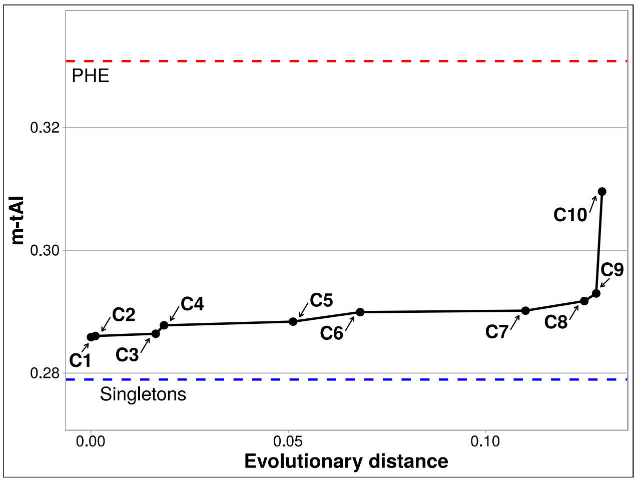


**b6**

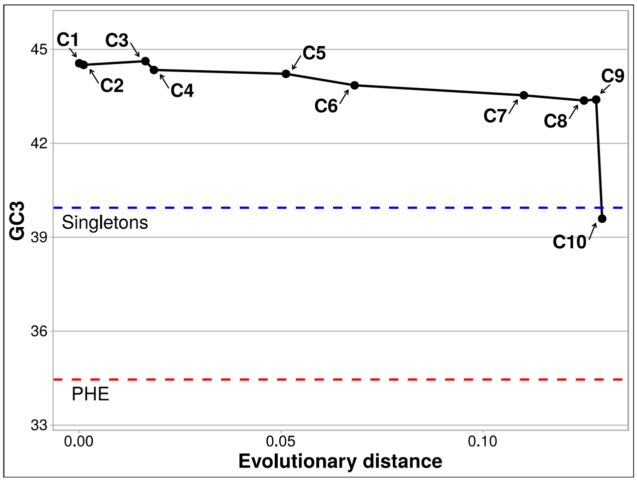

*Moraxella bovis* CCUG 2133 NZ_MUXV01000108

**a7**

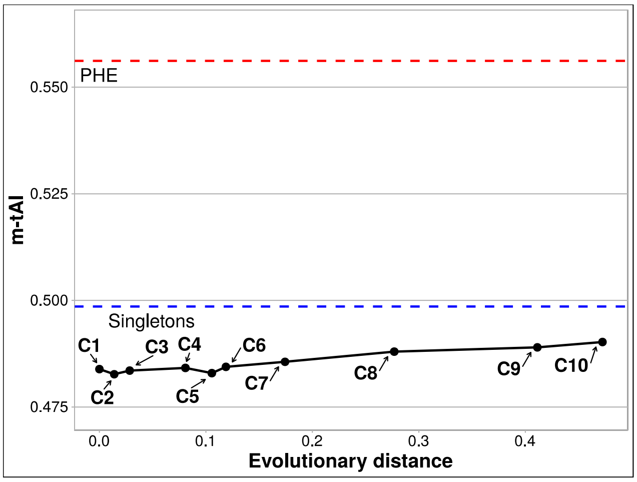


**b7**

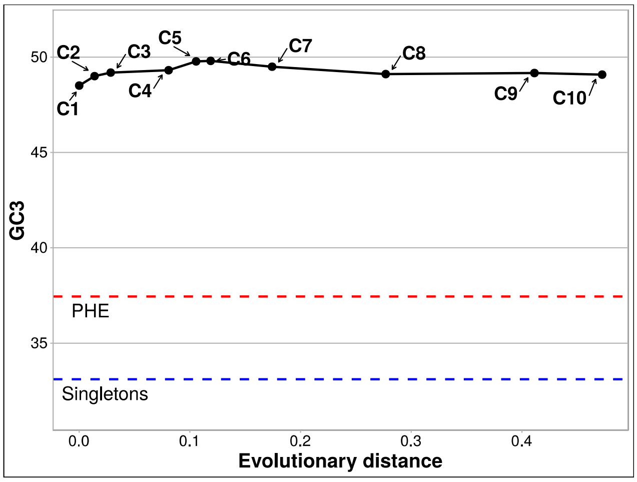


*Chromobacterium violaceum* ATCC12472

**a8**

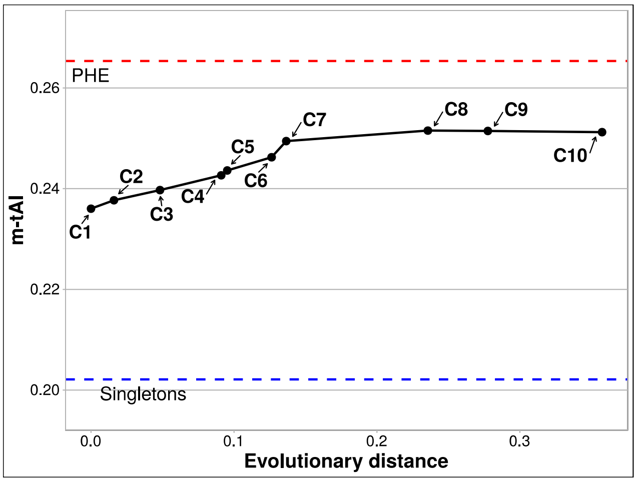


**b8**

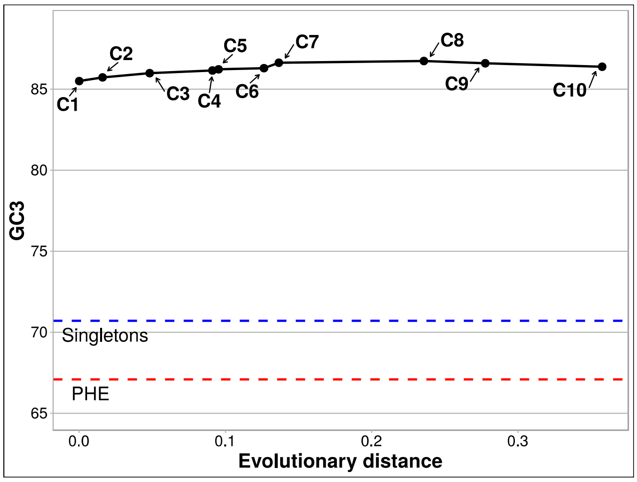


*Paenibacillus graminis* DSM 15220 NZ_CP009287

***a9***

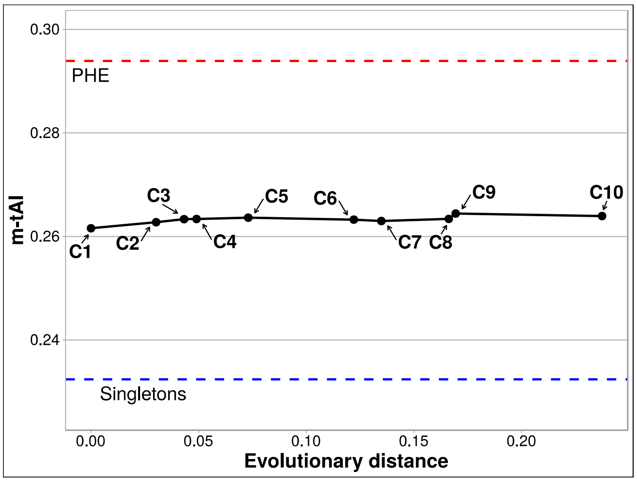


***b9***

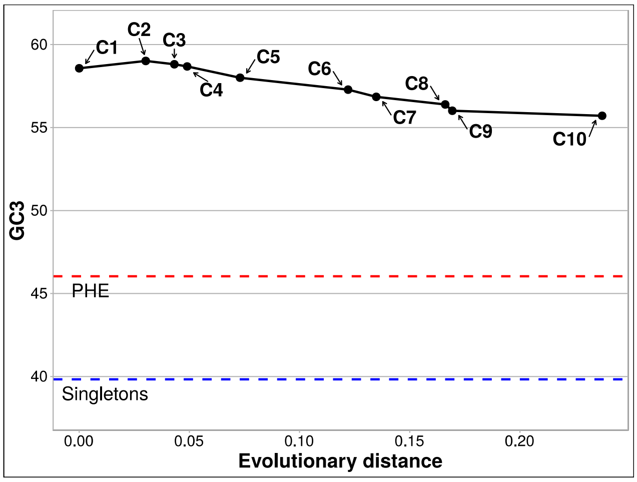

*Photobacterium gaetbulicola* Gung47

**a10**

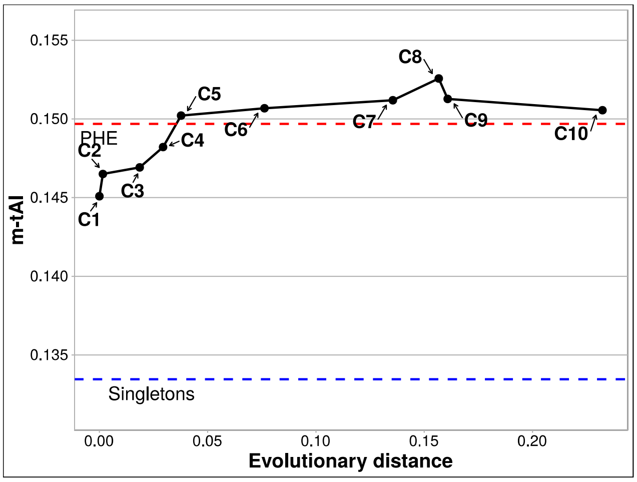


**b10**

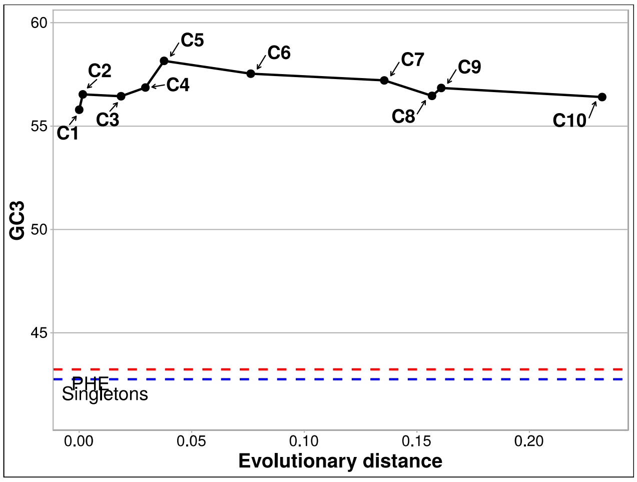


#### Group C

*Treponema succinifaciens* DSM 2489 NC_015385

**a11**

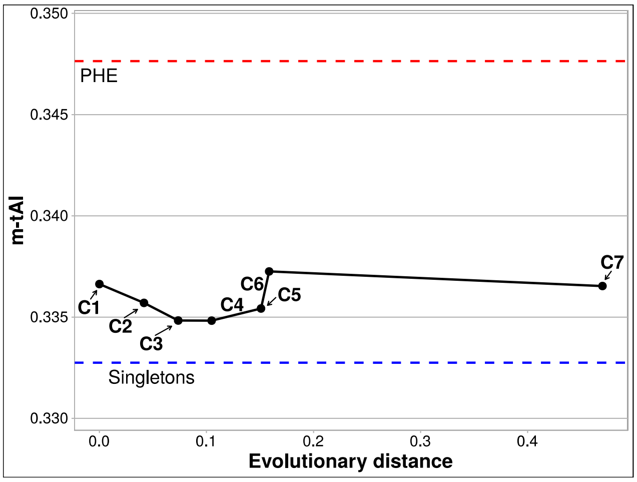


**b11**

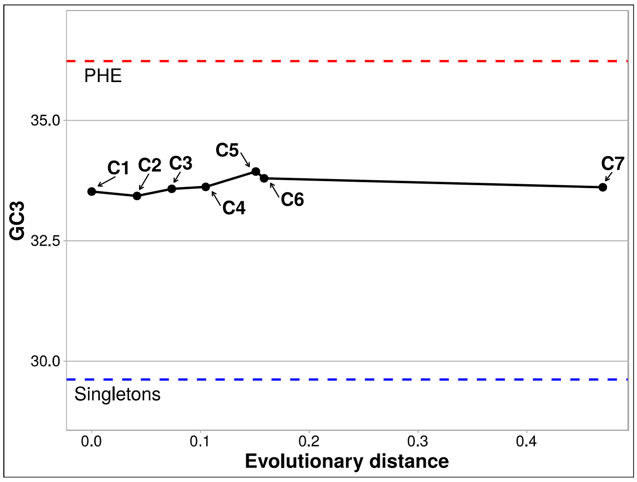


*Prevotella melaninogenica* ATCC 25845 NC_014371

**a12**

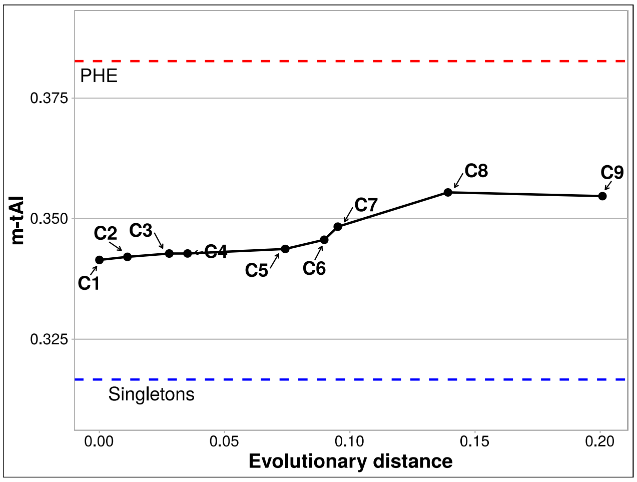


**b12**

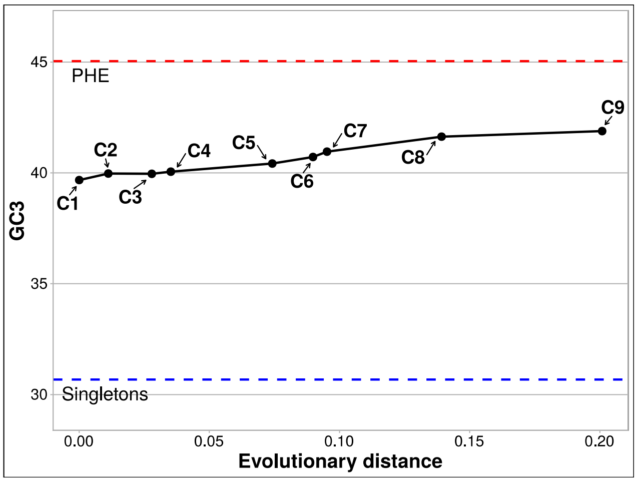

*Yersinia enterocolitica* subsp palearctica Y11 NC_017564

**a13**

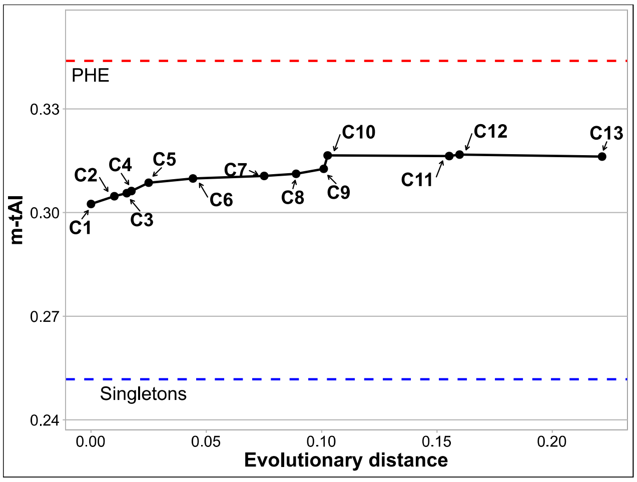


**b13**

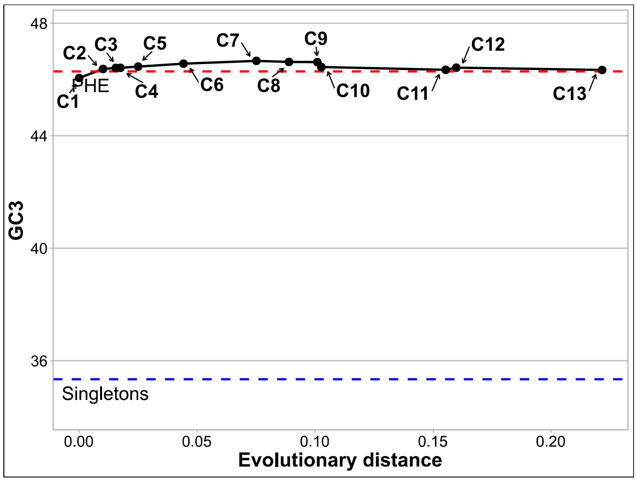


*Methanolacinia petrolearia* DSM 11571 NC_014507

**a14**

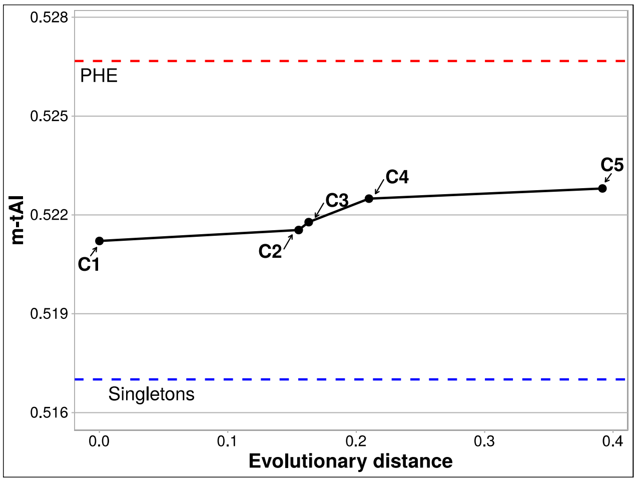


**b14**

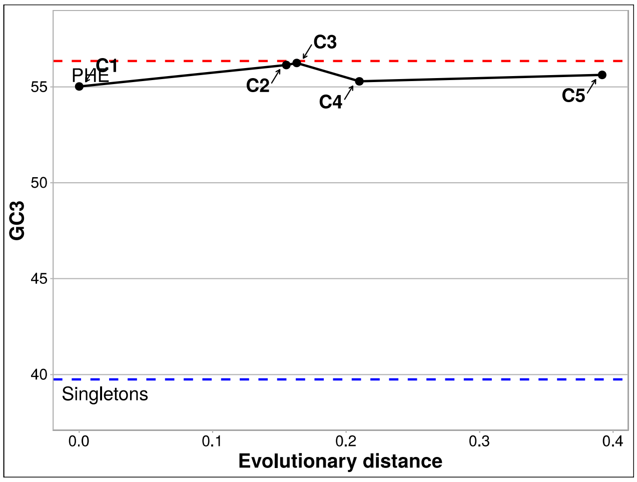


*Bifidobacterium longum* subsp longum JCM 1217 NC_015067

**a15**

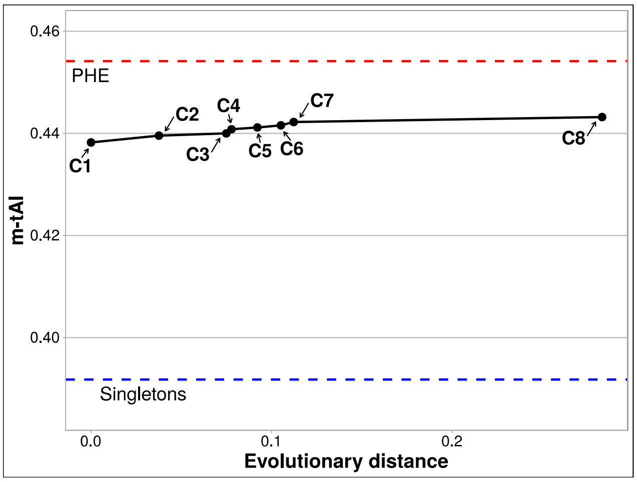


**b15**

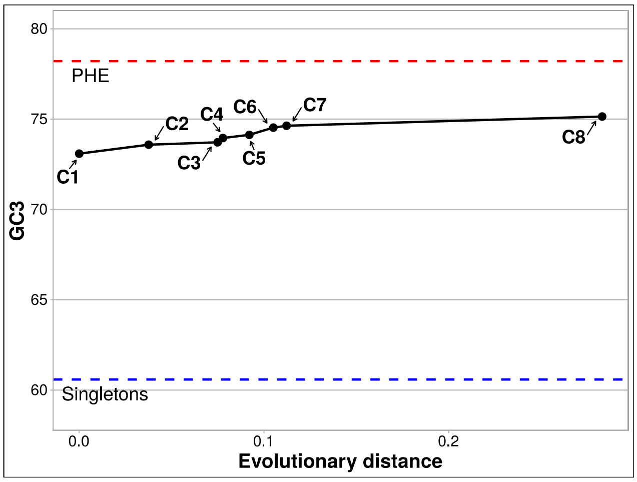

*Bordetella holmesii* ATCC 51541 NZ_CP007494

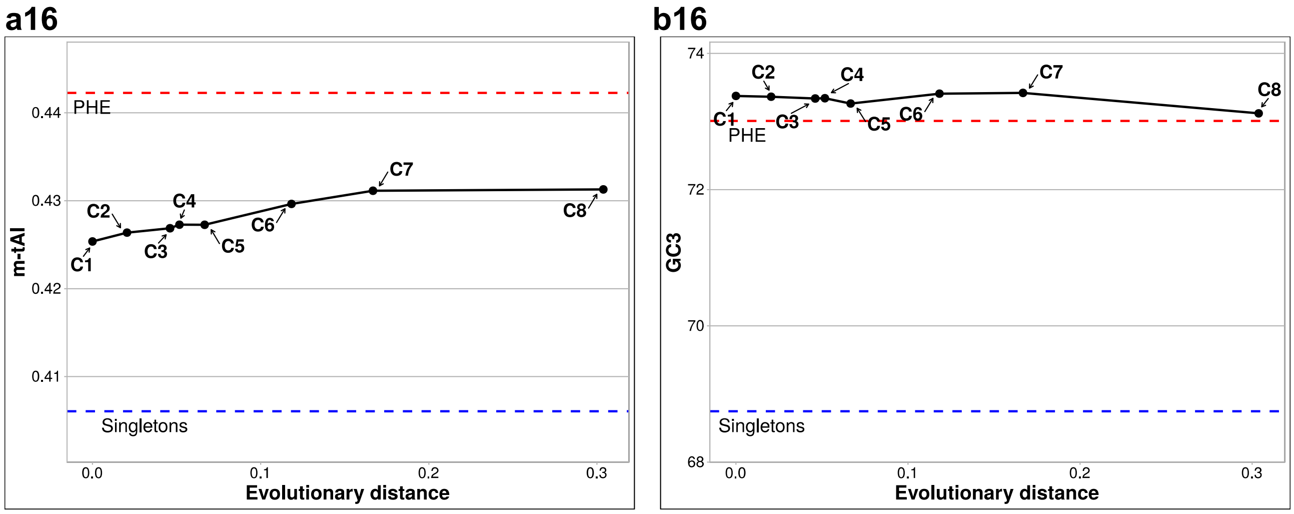


*Mycobacterium fortuitum* subsp fortuitum DSM 46621 ATCC 6841 CP014258

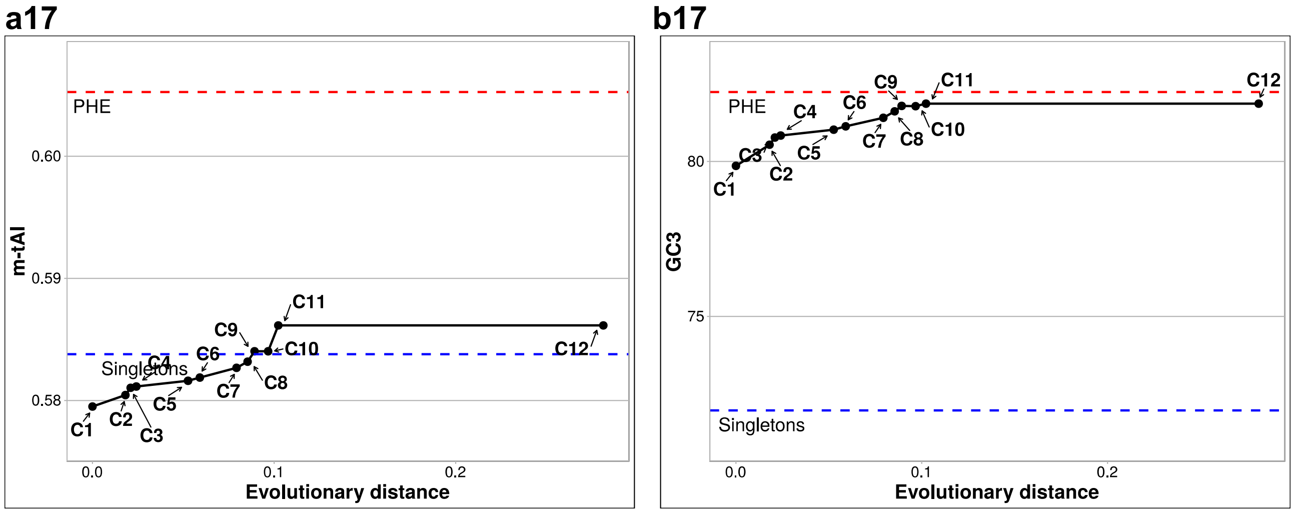


*Sphingomonas parapaucimobilis* NBRC 15100 NZ_BBPI01000001

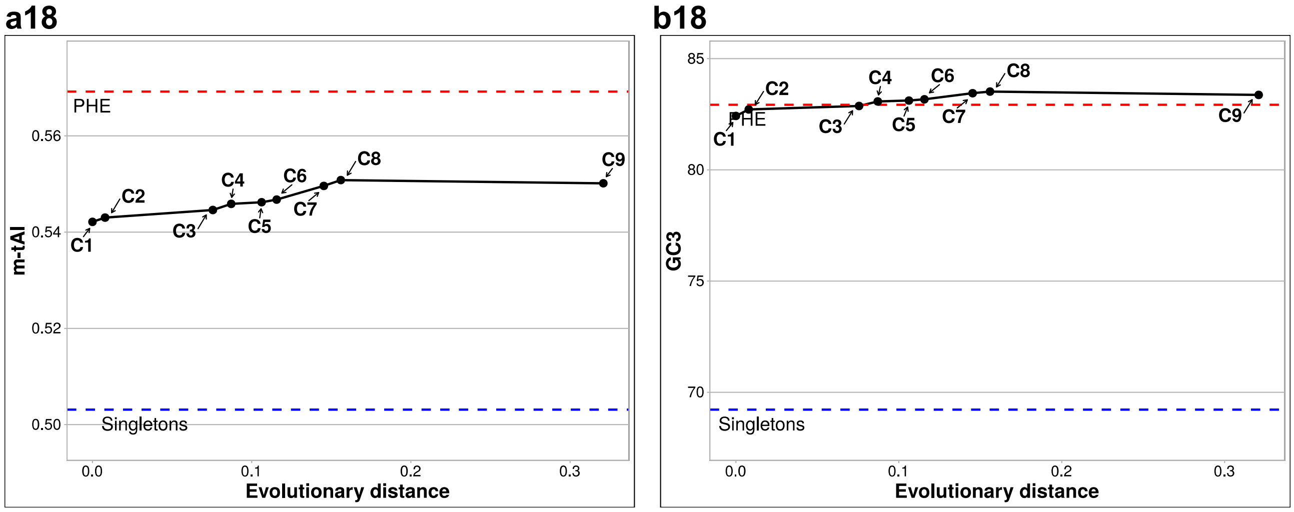

*Atopobium parvulum* DSM20469

**a19**

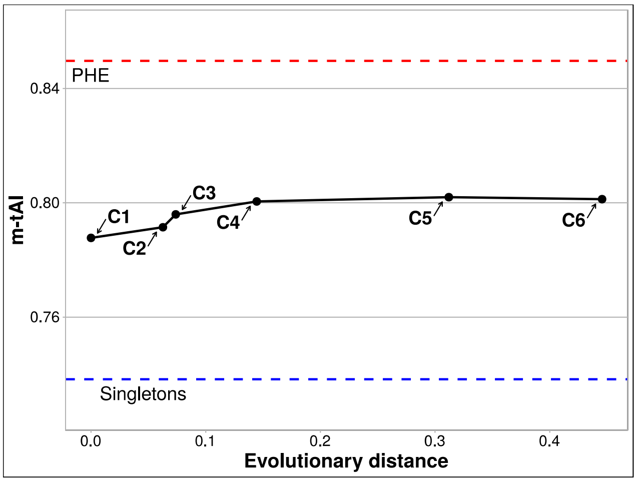


**b19**

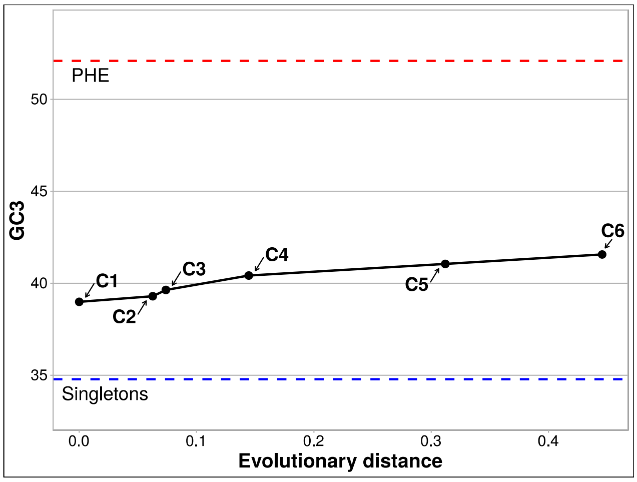


#### Group D

*Leisingera methylohalidivorans* DSM 14336

**a20**

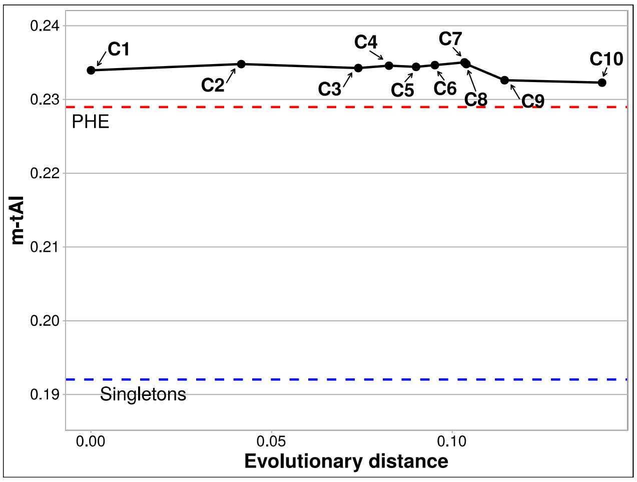


**b20**

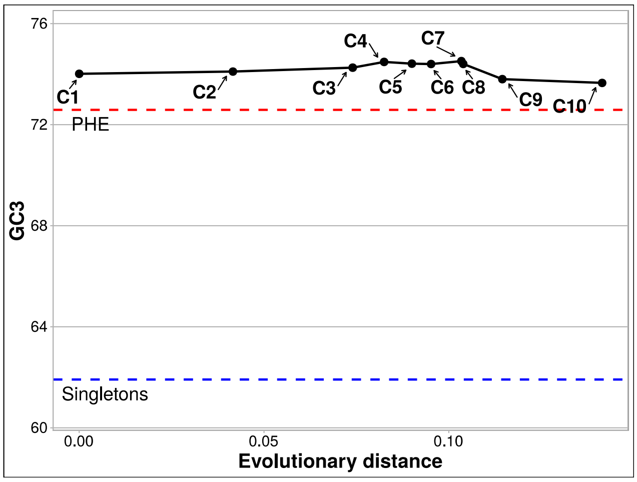


*Oceanibaculum pacificum* MCCC 1A02656 NZ_LPXN01000181

**a21**

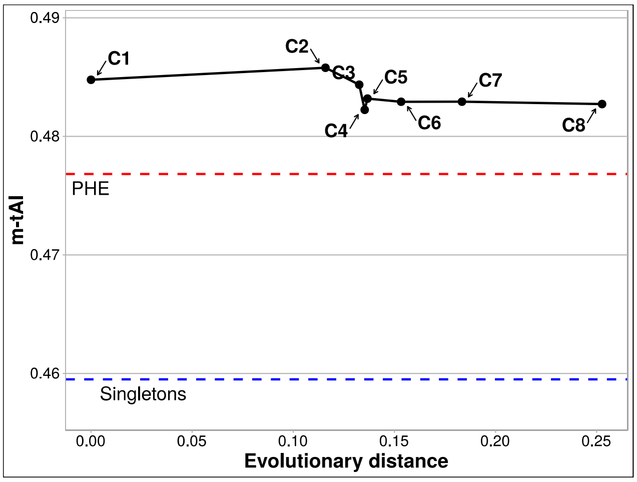


**b21**

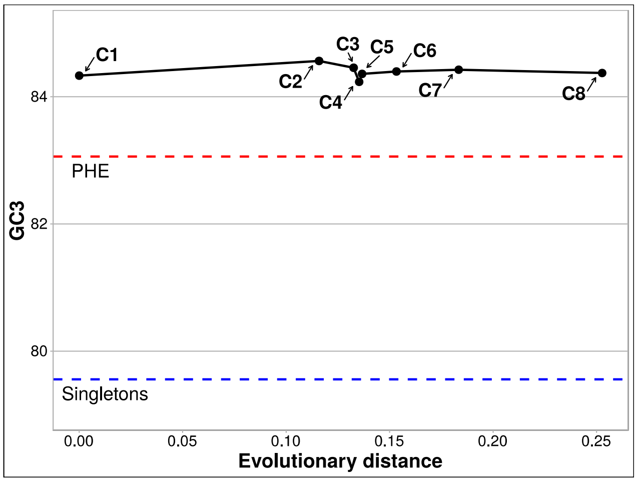

#### *Arthrobacter enclensis* NIO 1008 NZ_KQ758616

**a22**

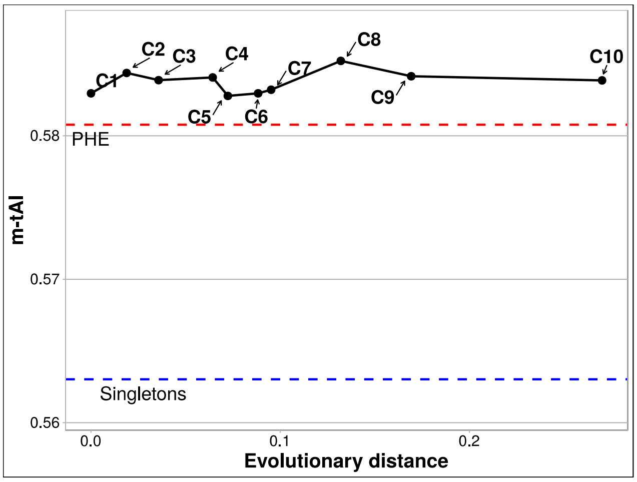


**b22**

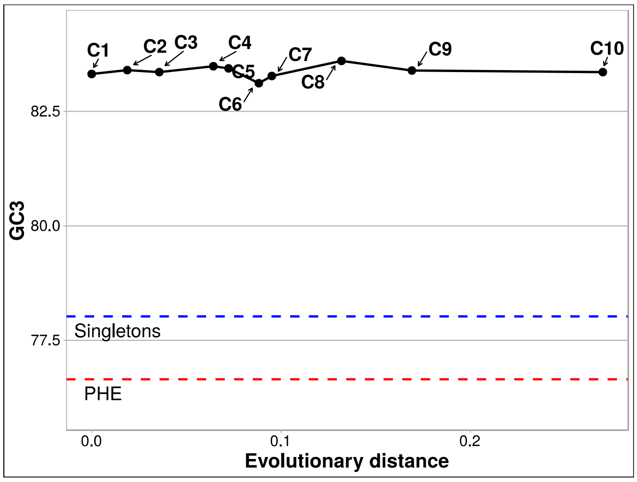


#### *Rhodanobacter denitrificans* 2APBS1

**a23**

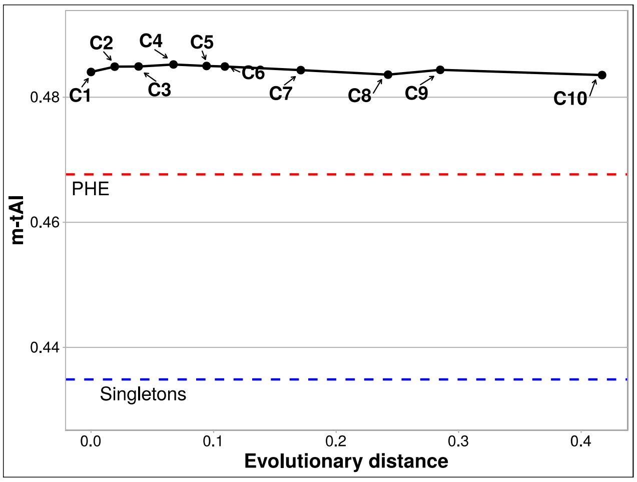


**b23**

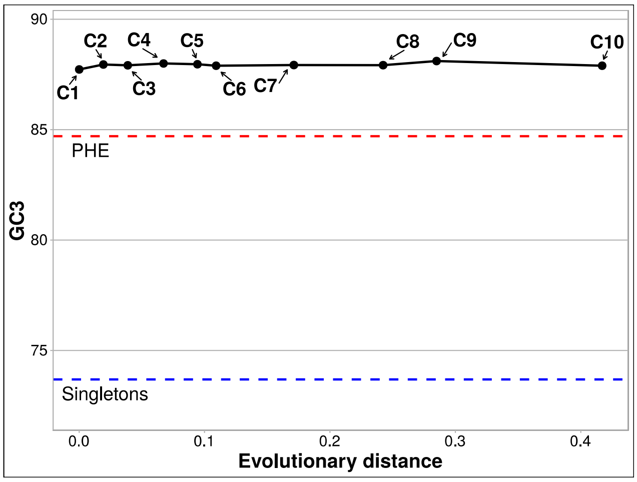


#### *Microbacterium aurum* KACC 15219 NZ_CP018762

**a24**

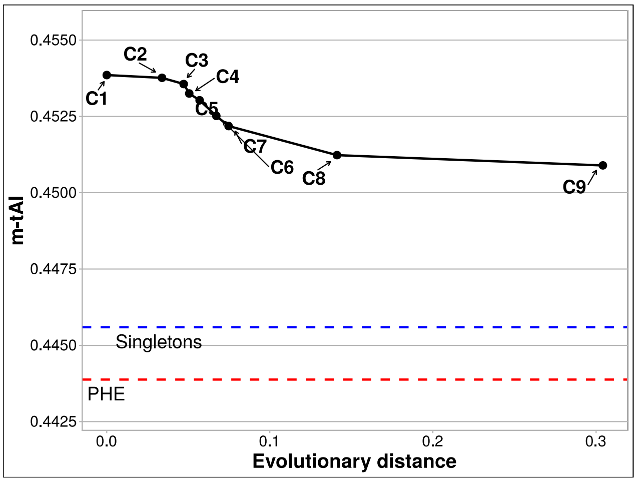


**b24**

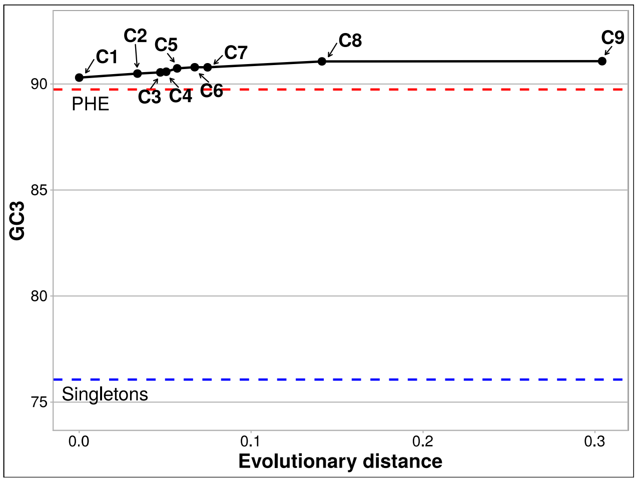

*Roseomonas mucosa* ATCC BAA 692 NZ_JHWD01000081

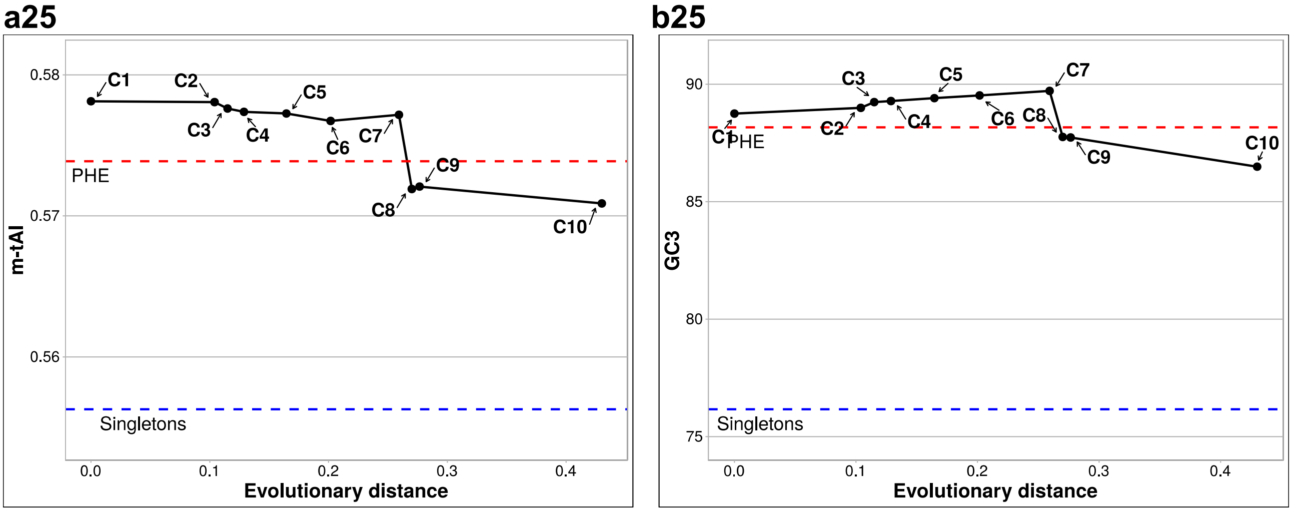

**Fig. S3. Codon-usage adaptations to the cellular tRNA-pool, and changes in the GC3 content of core-gene sets with different degrees of conservation throughout the phylogeny of 25 prokaryote families.** The strains represented here are the same as the 25 in Fig. S1. **Panels a1 to a25.** In each figure, the modal tRNA-adaptation index (m-tAI) calculated for each of the Ci gene sets as described in Materials and Methods is plotted on the *ordinate* as a function of the evolutionary distance (Table S1_a, tab 2) indicated on the *abscissa* as inferred from the corresponding phylogenetic trees included in Table S1_a-c. Higher values of m-tAI indicate an enrichment in the codon-usage frequencies of those synonymous codons better adapted to the host-cell tRNA pool. The C1 to Cn gene sets plotted here are the same as those presented in Fig. 1. The red and blue horizontal dashed lines correspond to the respective m-tAI values calculated for the PHE genes and the singletons. **Panels b1 to b25.** In each panel, the average GC3 content in each core-gene set of increasing ancestry is plotted on the *ordinate* as a function of the evolutionary distance indicated on the *abscissa* as in Panels a1 to a25. The PHE genes and the singletons are represented as red and blue horizontal dashed lines, respectively.
