## Supplementary material for "Codon-usage optimization in the prokaryotic tree of life: How synonymous codons are differentially selected in sequence domains with different expression levels and degrees of conservation": Suplemental Figure 4

### Fig. S4

#### Group A

*S4-1 Methanobrevibacter smithii* ATCC 35061

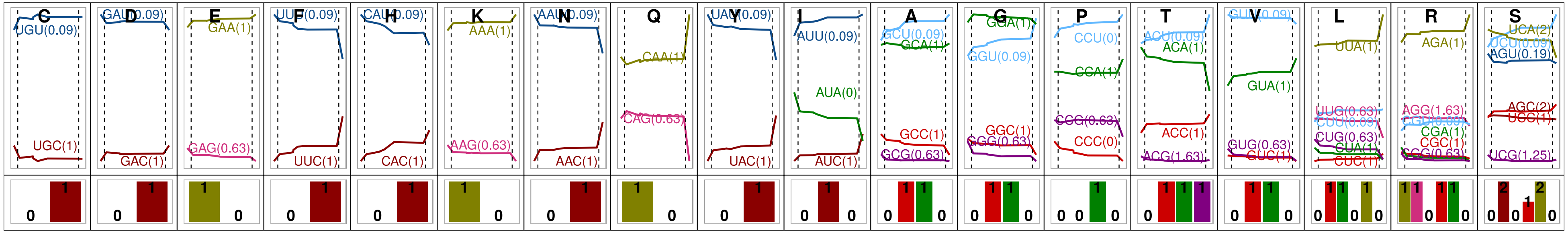

*S4-2Tetragenococcus halophilus* NBRC 12172 NC_016052

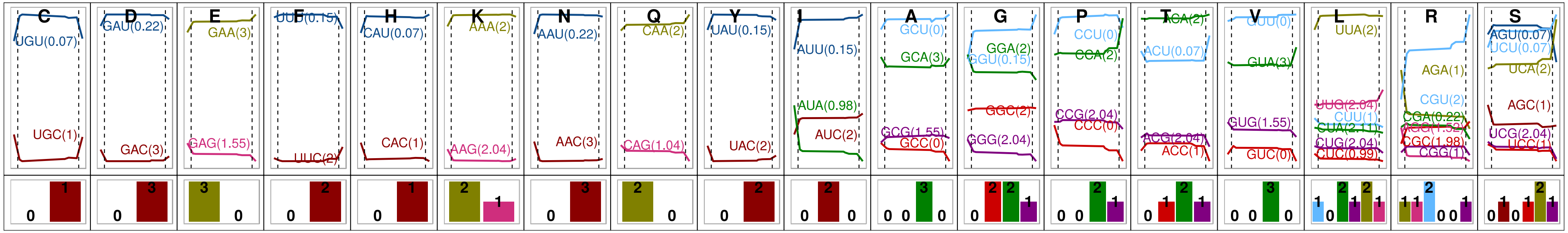

#### Group B

*S4-3 Sulfurospirillum multivorans* DSM 12446 NZ_CP007201

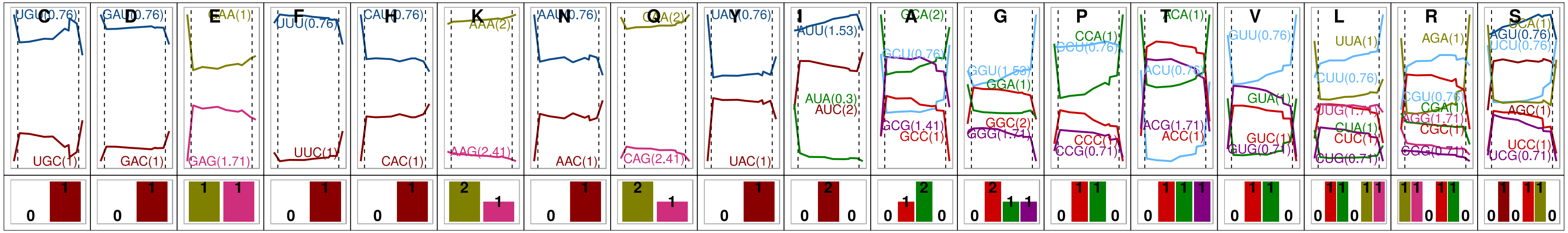

*S4-4 Streptococcus equi* ATCC 33398 NZ_FTNH01000046

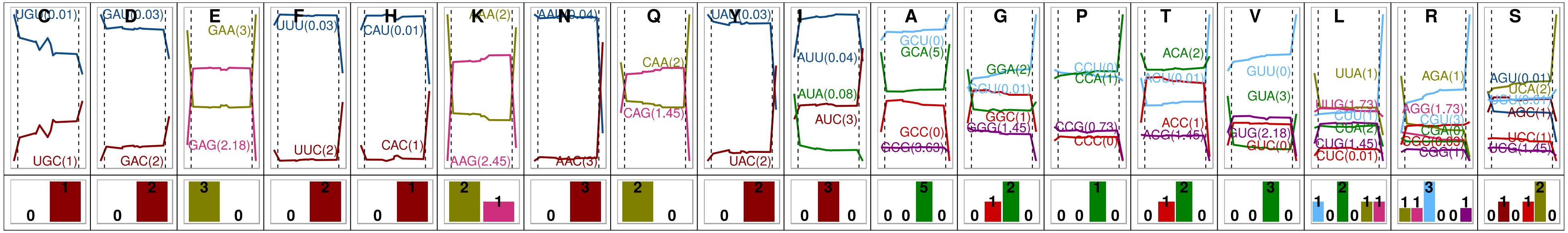


*S4-5 Bacteroides vulgatus* ATCC 8482 NC_009614

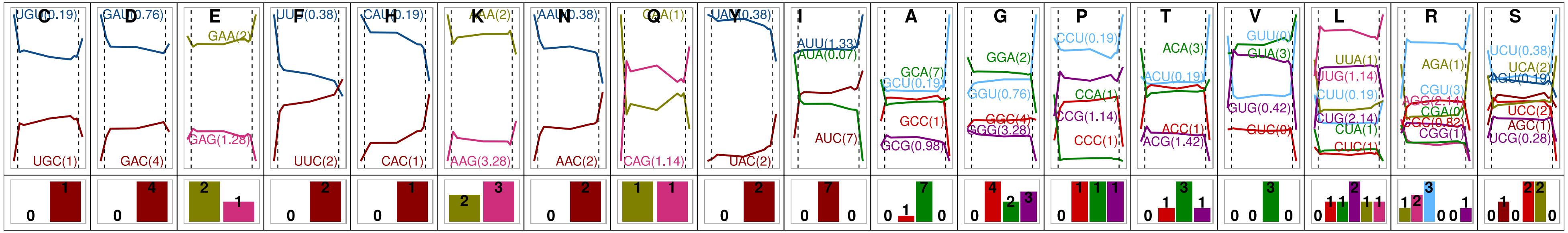


*S4-6 Bacillus subtilis* subsp spizizenii TU B 10 NC_016047

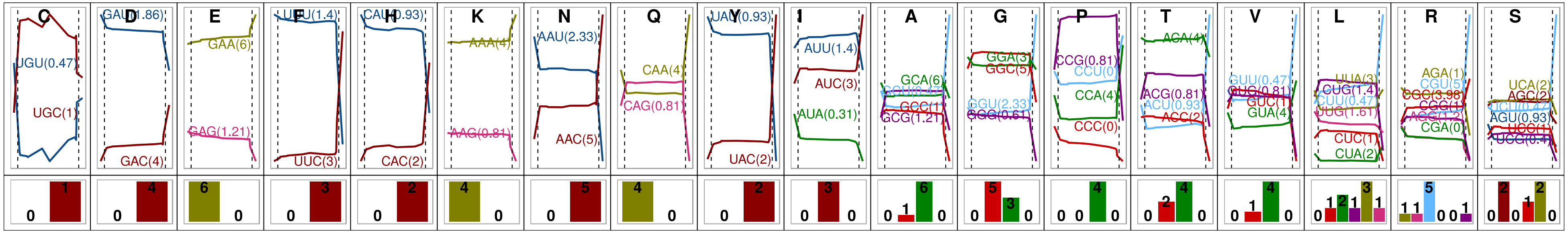


*S4-7 Moraxella bovis* CCUG 2133 NZ_MUXV01000108

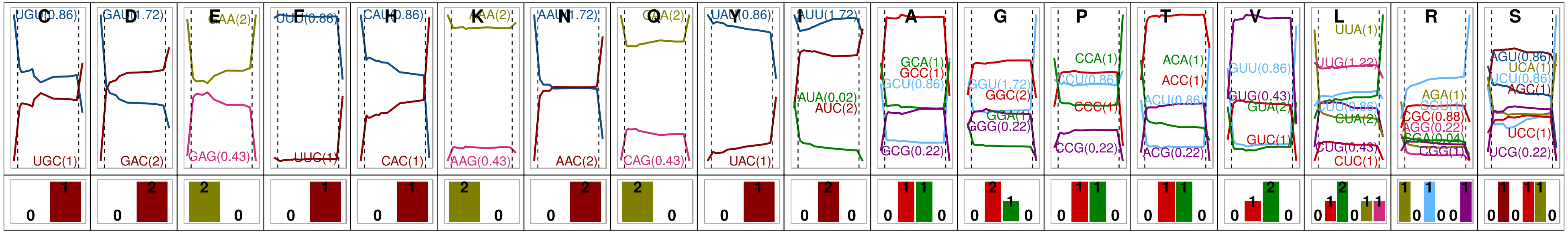

#### *S4-8 Chromobacterium violaceum* ATCC12472

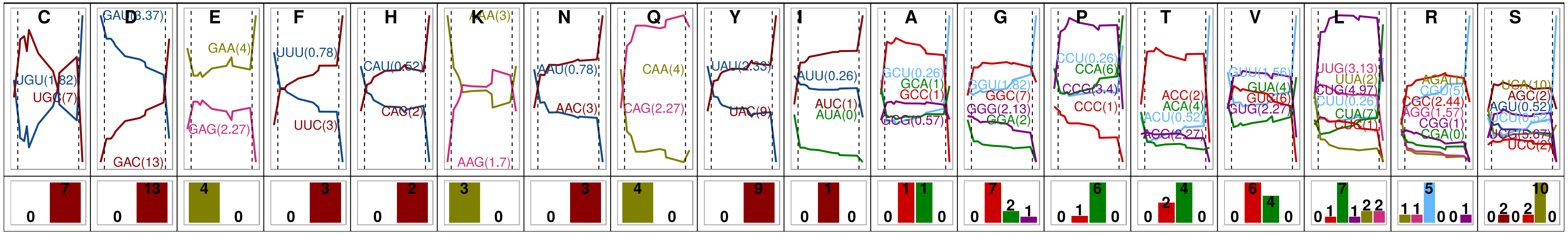


#### *S4-9 Paenibacillus graminis* DSM 15220 NZ_CP009287

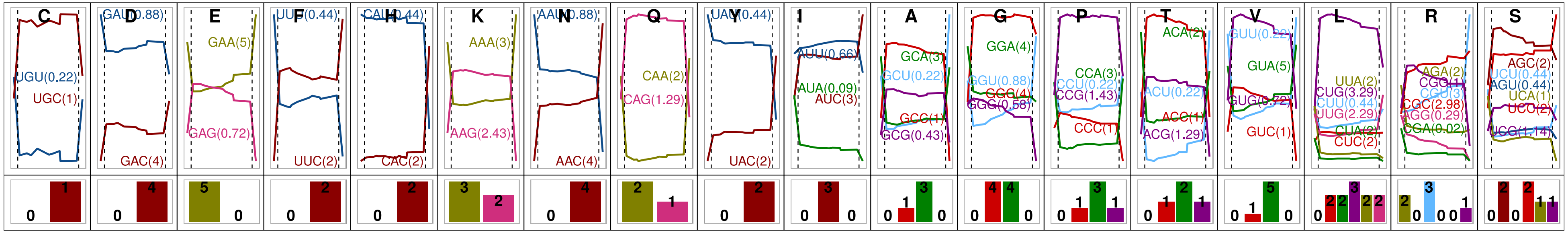


#### *S4-10 Photobacterium gaetbulicola* Gung47

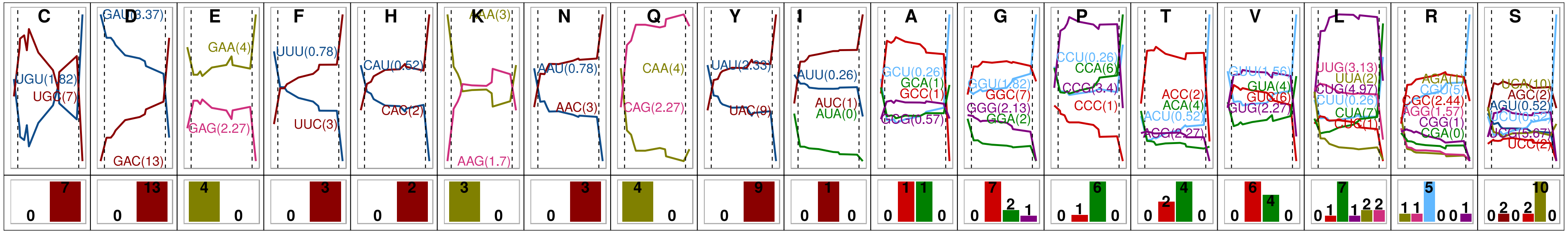

#### Group C

*S4-11 Treponema succinifaciens* DSM 2489 NC_015385

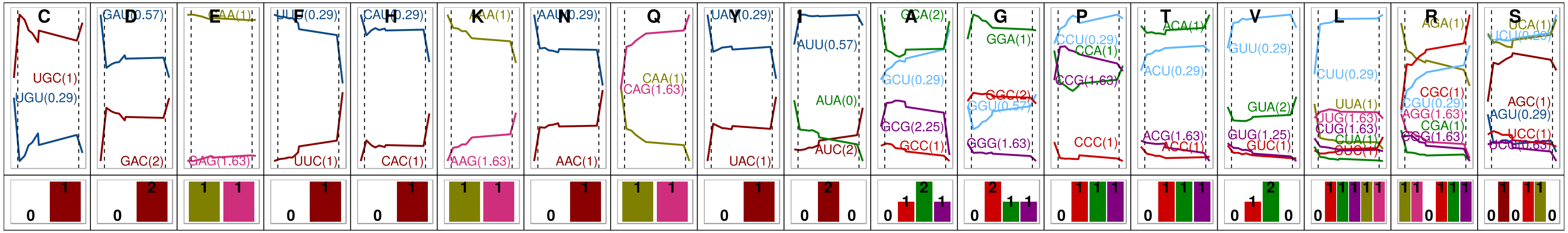


*4-12 Prevotella melaninogenica* ATCC 25845 NC_014371

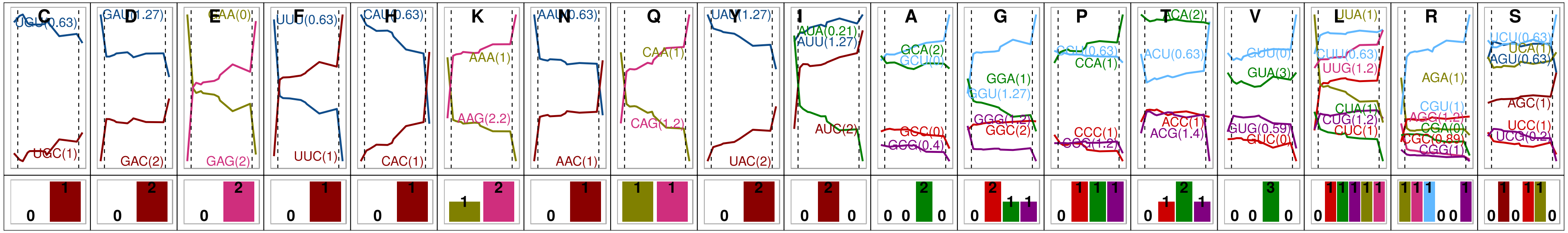


*S4-13 Yersinia enterocolitica* subsp palearctica Y11 NC_017564

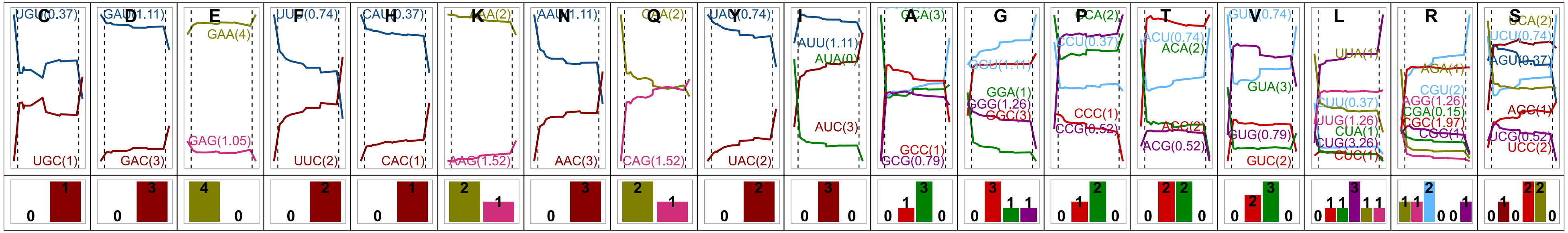

*S4-14 Methanolacinia petrolearia* DSM 11571 NC_014507

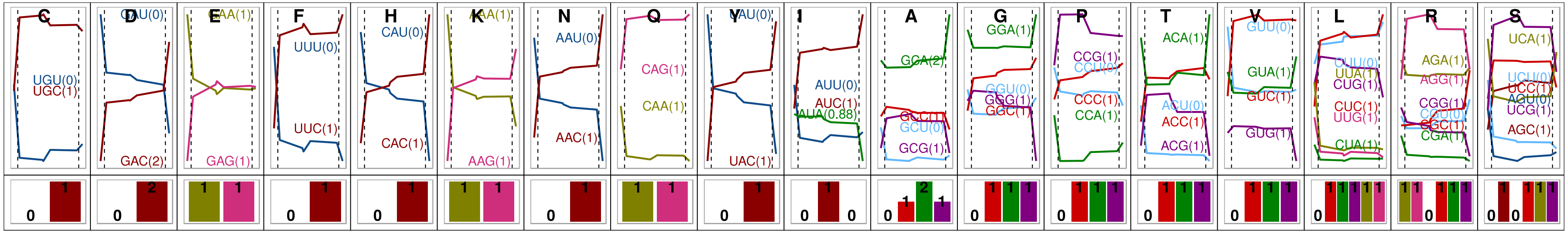

*S4-15 Bifidobacterium longum* subsp longum JCM 1217 NC_015067

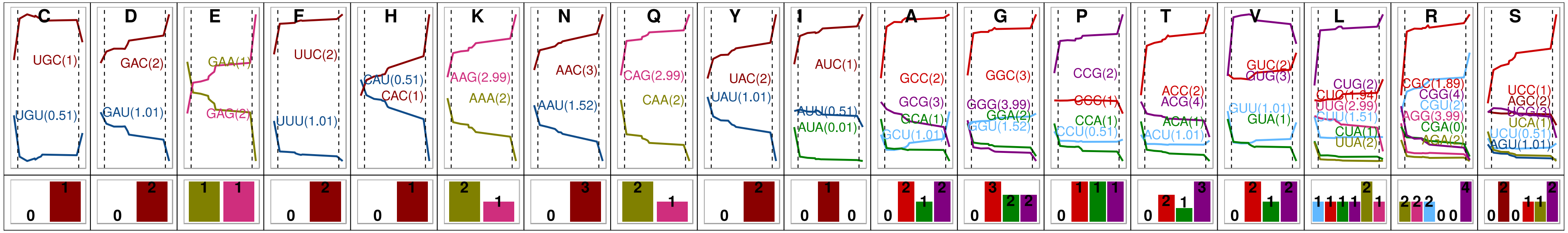

*S4-16 Bordetella holmesii* ATCC 51541 NZ_CP007494

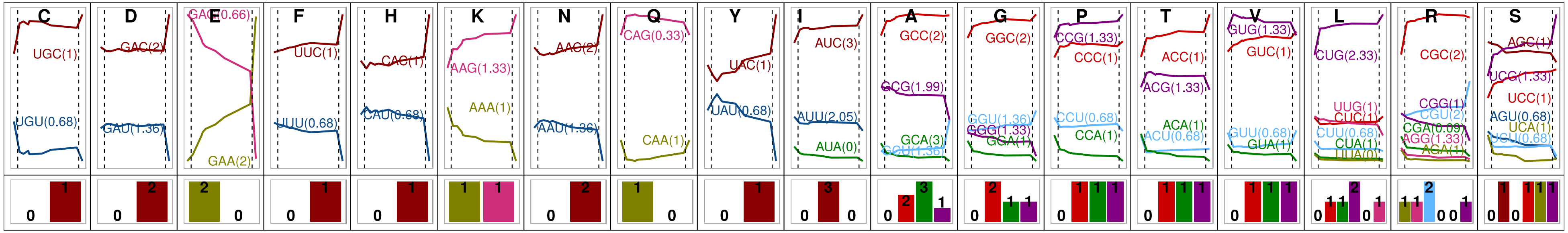

*S4-17 Mycobacterium fortuitum* subsp fortuitum DSM 46621  ATCC 6841 CP014258

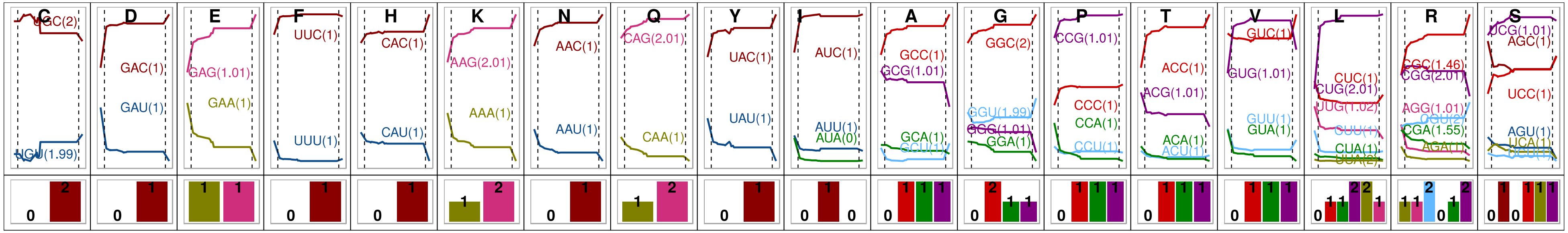


*S4-18 Sphingomonas parapaucimobilis* NBRC 15100 NZ_BBPI01000001

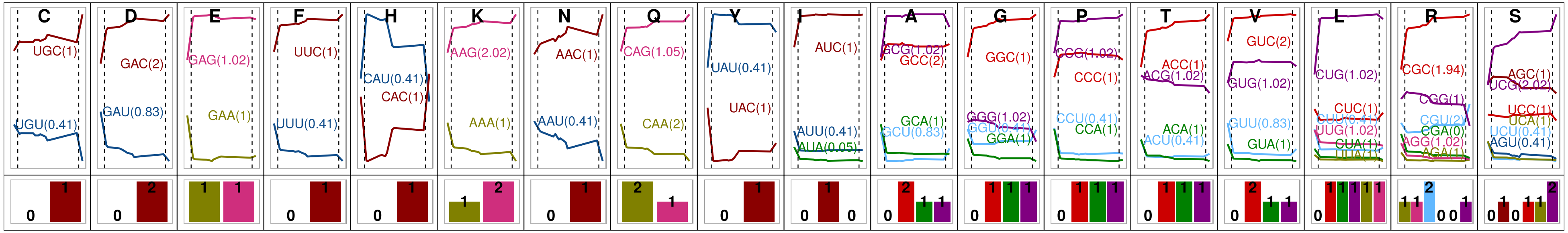


*S4-19 Atopobium parvulum* DSM20469

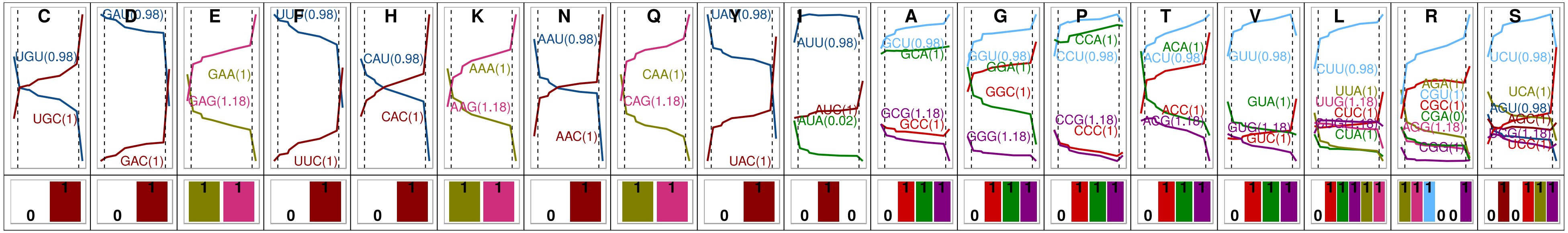

#### Group D

*S4-20 Leisingera methylohalidivorans* DSM 14336

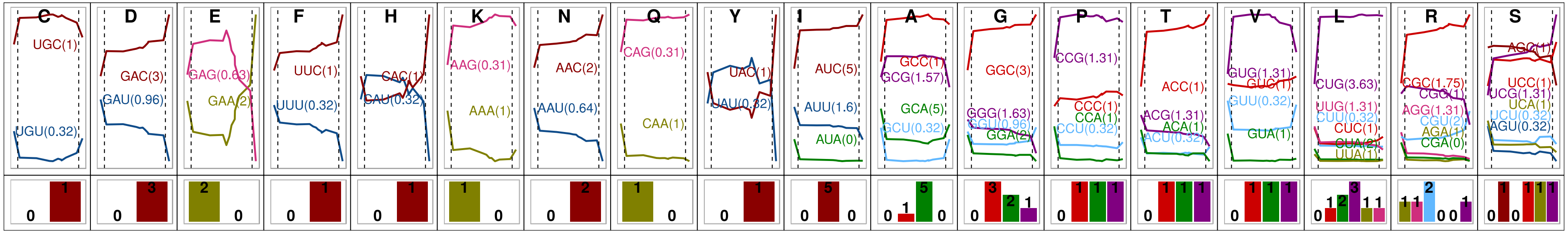


*S4-21 Oceanibaculum pacificum*  MCCC 1A02656 NZ_LPXN01000181

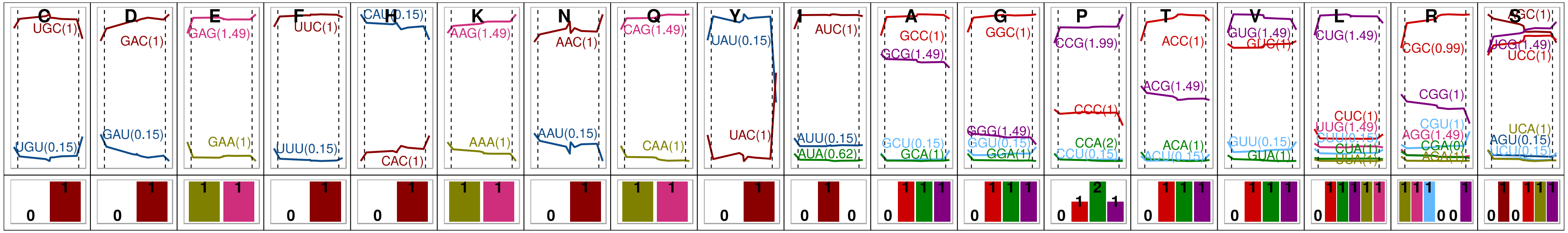


*S4-22 Arthrobacter enclensis*  NIO 1008 NZ_KQ758616

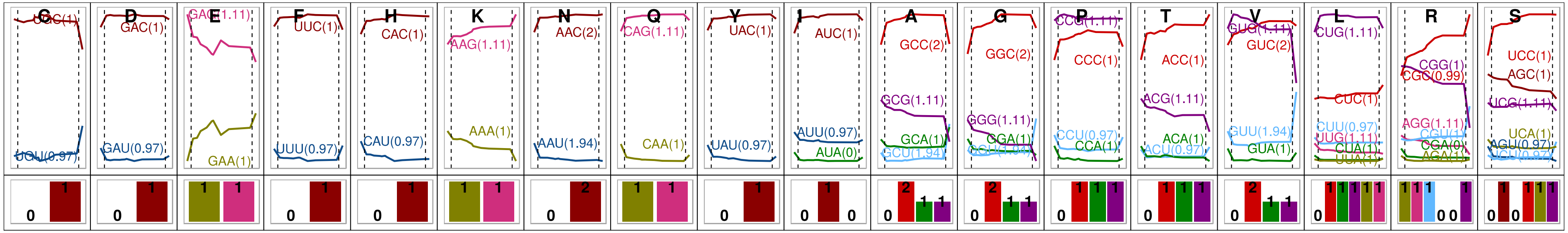

*S4-23 Rhodanobacter denitrificans* 2APBS1

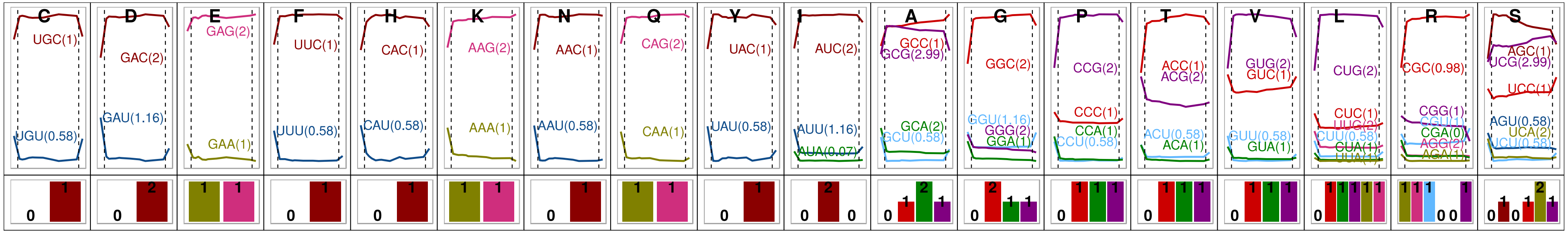


*S4-24 Microbacterium aurum* KACC 15219 NZ_CP018762

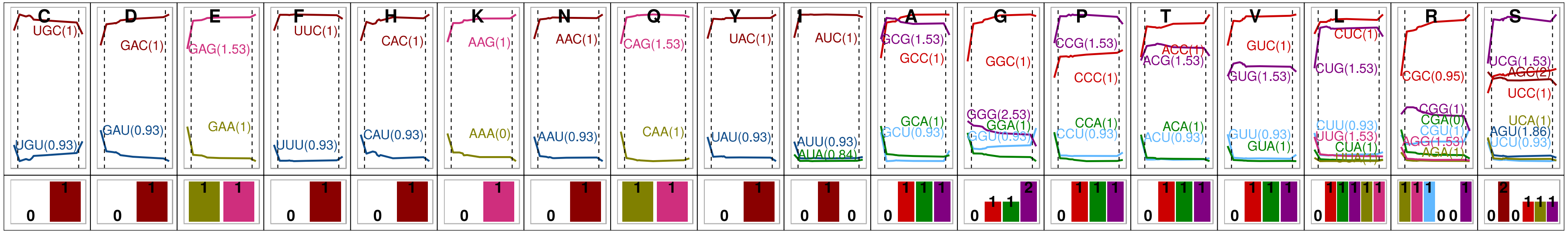


*S4-25 Roseomonas mucosa* ATCC BAA 692 NZ_JHWD01000081

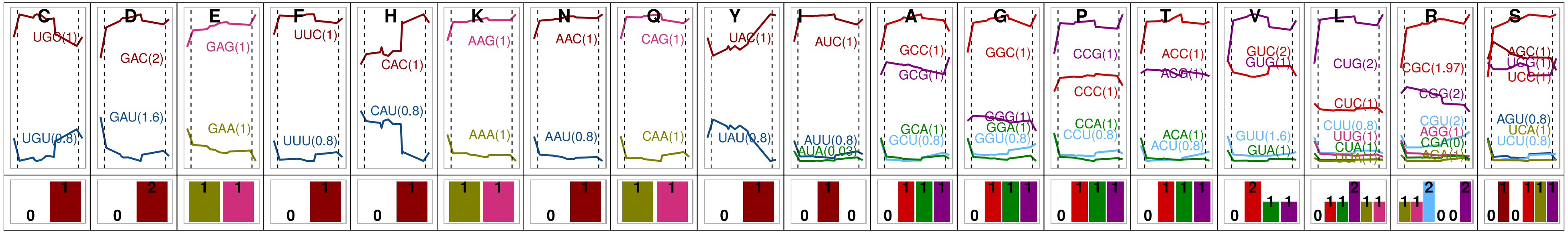

**Fig. S4. Codon usage frequencies and adaptation indices (*Wi*s) of the gene sets analyzed in this work, together with the tRNA-gene–copy numbers for strains of 25 different prokaryote families included in Groups A to D.** For the amino acid denoted by the corresponding single-letter identification code located above each panel, the change in the codon-usage frequencies (CUFs; see Materials and Methods) of the core-gene sets with increasing ancestries (left to right, the C1 to Cn), the PHE genes, and the singletons are plotted in the upper panels as solid horizontal curves for each of the indicated codon triplets between the two vertical broken lines, for the singletons to the left of the first of those lines, and for the PHE genes to the right of the second (with singletons and PHE genes being located at the beginning and the end of the curves, respectively). The CUFs are represented by different colors with the associated codon-adaptation index (*Wi*) being denoted within parentheses beside each triplet. Finally, the presence and gene-copy number (N tRNA) of the cognate tRNA species of a given synonymous codon bearing the exact complementary anticodon is depicted with a number and a bar of proportional height in the lower panel in the same color as the corresponding triplet and curve in the upper panel.
