## Supplementary material for "Codon-usage optimization in the prokaryotic tree of life: How synonymous codons are differentially selected in sequence domains with different expression levels and degrees of conservation": Suplemental Figure 5

### FIGURE S5

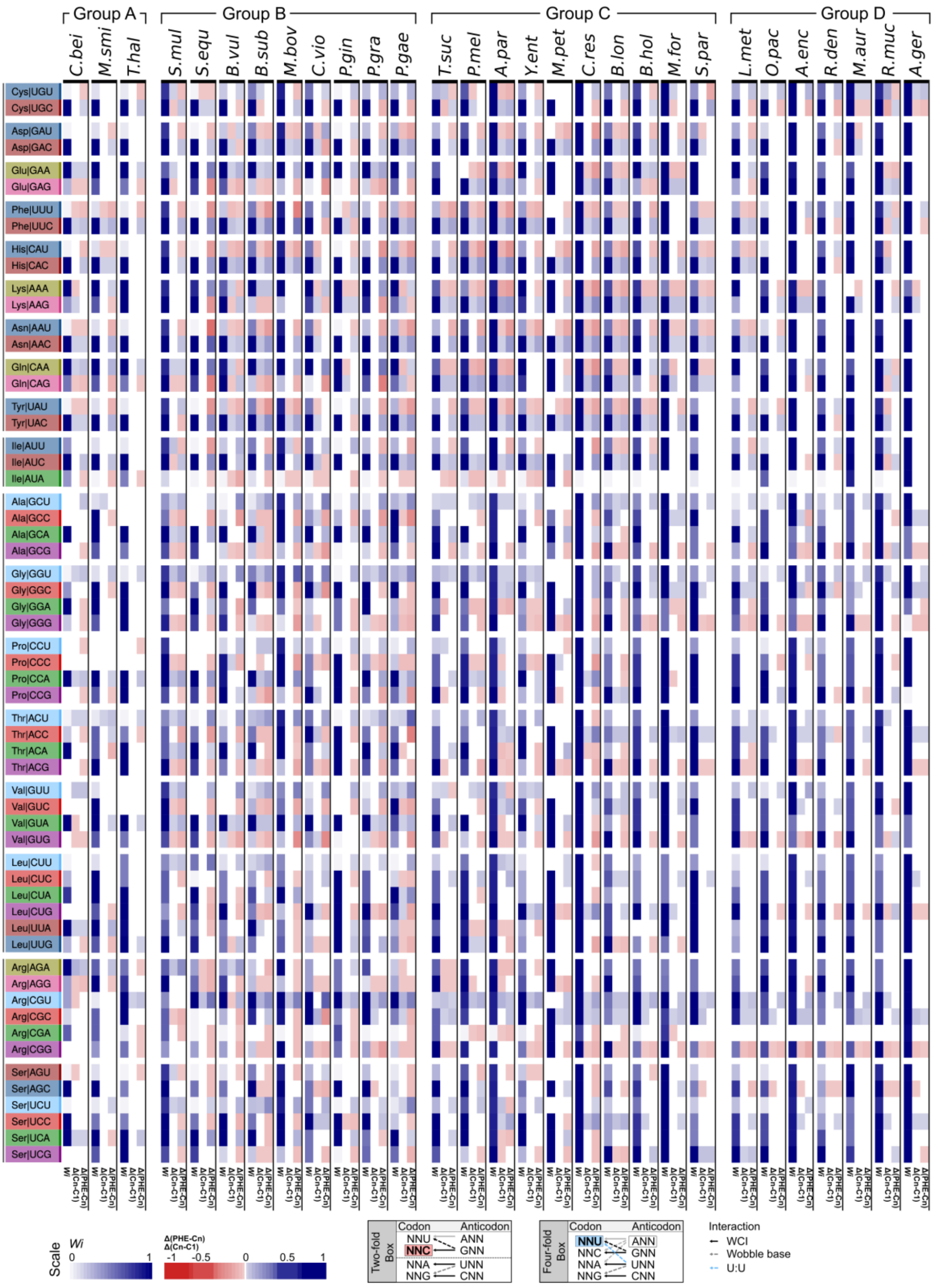


**Fig. S5. General codon-enrichment profiles in reference strains of the 29 prokaryote families analyzed.** The reference species from Groups A, B, C, and D are denoted above the figure. The enrichment in a given i$^{th}$ codon over ancestry was calculated as $_{CODON``i''}CUF^{Cn}$ – $_{CODON``i''}CUF^{C1}$, referred to simply as ∆(Cn-C1). In like manner, the enrichment in the PHE genes over ancestry was calculated as $_{CODON``i''}CUF^{PHE}$ – $_{CODON``i''}CUF^{Cn}$, similarly referred to as ∆(PHE-Cn). The rectangular boxes to the left of the figure indicating the codons in the different amino acids are color-coded according to the nature of the 3' base present as follows: blue-variants, U; red-variants, C; green-variants, A; and violet-variants G. The vertical rectangles at the left of each small panel represent the *Wi*s, with the intensity of the blue color being proportional to the values for each codon. The *Wi* color intensities are furthermore normalized to the maximum *Wi* value in each amino-acid–codon family. The vertical rectangles in the middle and at the right of each small panel represent the ∆(Cn-C1) or ∆(PHE-Cn), with the color intensity being in proportion to either an increased (blue), a decreased (red), or an equal (white) CUF in the gene sets under analysis (Cn *versus* C1, PHE *versus* Cn), as indicated in the color keys below the figure. The variations in codon-anticodon interactions (WCI, wobble base, U:U) along with their participation in the C (red) and U biases (light blue) are illustrated in the boxes below the figure.
